# On the nature of the earliest known lifeforms

**DOI:** 10.1101/2021.08.16.456462

**Authors:** Dheeraj Kanaparthi, Frances Westall, Marko Lampe, Baoli Zhu, Thomas Boesen, Bettina Scheu, Andreas Klingl, Petra Schwille, Tillmann Lueders

## Abstract

Microfossils from the Paleoarchean Eon are the oldest known evidence of life. Despite their significance in understanding the history of life on Earth, any interpretation of the nature of these microfossils has been a point of contention among researchers. Decades of back-and-forth arguments led to the consensus that reconstructing the lifecycles of Archaean Eon organisms is the most promising way of understanding the nature of these microfossils. Here, we transformed a Gram-positive bacterium into a primitive lipid vesicle-like state and studied it under environmental conditions prevalent on early Earth. Using this approach, we successfully reconstructed morphologies and life cycles of Archaean microfossils. In addition to reproducing microfossil morphologies, we conducted experiments that spanned years to understand the process of cell degradation and how Archaean cells could have undergone encrustation minerals (in this case, salt), leading to their preservation as fossilized organic carbon in the rock record. These degradation products strongly resemble fossiliferous features from Archaean rock formations. Our observations suggest that microfossils aged between 3.8 to 2.5Ga most likely were liposome-like protocells that have evolved physiological pathways of energy conservation but not the mechanisms to regulate their morphology. Based on these observations, we propose that morphology is not a reliable indicator of taxonomy in these microfossils.

## Introduction

The Pilbara Greenstone Belt (PGB), Western Australia, and Barberton Greenstone Belt (BGB), South Africa, are the oldest known sedimentary rock successions that have not undergone significant metamorphic alterations (1,2). Hence, these rock formations have been the subject of numerous scientific investigations focused on understanding the biology and biogeochemistry of Archaean Earth (1–5). Over a span of 50 years, these studies have documented various organic structures within these rock formations that resemble fossilized cells and their degradation products, with δ^13^C composition consistent with biologically derived organic carbon (6,7). Although these observations suggest that these organic structures were fossil remnants of Archaean microorganisms, any such interpretation, together with the biological origin of these structures, has been a point of contention among researchers (8–11). Two factors currently limit wider acceptance of their biological origin – the absence of truly analogous microfossil morphologies among extant prokaryotes and an indication of an ongoing biological process, like cell division, among most microfossils (4,12). Moreover, most of the described microfossils are larger than present-day prokaryotes and often exhibit considerable cytoplasmic complexity with intracellular alveolar structures (3,5,13). These complex morphologies and relatively larger cell sizes of supposedly primitive Archaean Eon cells is not in accordance with our current understanding of how biological complexity evolved through Darwinian evolution (14,15).

Apart from the chemical and δ^13^C-biomass composition (13,16–18), one key emphasis of the studies arguing for and against the biological origin of Archaean Eon organic structures involves an extensive morphological comparison with extant prokaryotes or abiotically formed minerals (8,9,19,20). Cell morphology among extant organisms is maintained by a plethora of intracellular processes and is determined by the information encoded in their genome (21). In our opinion, drawing parallels between the present-day prokaryotes and Archaean Eon organisms inherently involves subscribing to the notion that paleo-Archaean life forms possess all the complex molecular biological mechanisms to regulate their morphology as present-day cells. Any such presumptions are not in tune with the current scientific consensus of how life could have originated on early Earth (14,22,23). It is now widely believed that life evolved in the form of protocells devoid of most molecular biological complexity (23). These primitive cells are thought to have undergone slow Darwinian evolution, resulting in present-day cells with intricate intracellular processes (24,25). Given the unlikelihood of Archaean cells possessing complex molecular biological processes, we test the possibility that complex morphologies of Archaean microfossils result from the complete absence of intracellular mechanisms regulating their morphology. (26).

To test this hypothesis, we used a top-down approach of transforming a Gram-positive bacterium (*Exiguobacterium Strain-Molly*) into a primitive lipid vesicle-like state (*EM-P*). Cells in this state can be described as a simple sack of cytoplasm devoid of all mechanisms to regulate their morphology and reproduction. Although it has not been empirically demonstrated, some studies have suggested that cells in this lipid vesicle-like state may resemble primitive protocells (27–29). Given that the reproduction of such cells is shown to be influenced by environmental conditions (28,29), we studied the life cycle of these cells under experimental conditions resembling the native environment of the Archaean microfossils.

While the precise environmental conditions of early Earth remain uncertain, a growing consensus within the scientific community suggests that surface temperatures on Archaean Earth ranged between 26° and 35°C (30–32). Moreover, most, if not all, of the known microfossils from the Archaean Eon are restricted to coastal marine environments (6,33). Coastal marine environments often exhibit higher salinity due to the constant evaporation of seawater. To replicate the high salinities of the coastal marine environments, *EM-P* was cultivated in half-strength tryptic soy broth supplemented with 7% (w/v) Dead Sea Salt (TSB-DSS) at 30°C. We chose Dead Sea Salt over pure NaCl to better emulate complex salt compositions of natural environments.

Given that *EM-P*’s life cycle and the biophysical basis of such a reproduction process is extensively discussed in the previous paper (34), the primary focus of this manuscript will be restricted to the morphological comparison of *EM-P* cells and Archaean Eon microfossils. Below, we present the morphological comparison between *EM-P* cells and Archaean Eon microfossils. In addition to this morphological comparison, we also conducted experiments that spanned years (18 to 28 months) to understand the process of protocell degradation, how they become encrusted in salt, and how they are preserved as fossilized organic carbon in the rock record.

## Results

### Morphological comparison of top-down modified cells with fossilized Archaean cells

When cultured under experimental conditions likely resembling coastal marine environments of Paleoarchean Eon, *EM-P* exhibited cell sizes that were an order of magnitude larger than their original size. They also exhibited complex morphologies and reproduced by a relatively less understood process (28,34). The life cycle of these cells involves reproduction by two methods – via forming internal or a string of external daughter cells (34). *EM-P* reproducing by both these processes bears close morphological resemblance to microfossils reported from the Archaean Eon.

The first step in reproduction by intracellular daughter cells is the formation of hollow intracellular vesicles (Fig. 1A). These vesicles were formed by a process that resembles endocytosis (Fig. 1A). A similar process of vesicle formation was previously reported in protoplasts (29,37). Over time, the number of intracellular vesicles (ICVs) within *EM-P* gradually increased (Fig. 1A-F). No uniformity was observed in the size of ICVs within a cell or the number of ICVs among different cells (Fig. 1E, 1F, S1 & S2). *EM-P* cells with such intracellular vesicles resemble spherical microfossils reported from 3.46 billion-year-old (Ga) Apex chert (35). Like the Apex-chert microfossils, ICVs of *EM-P* were hollow, and organic carbon (cytoplasm) in these cells is restricted to spaces between the vesicles (Fig. 1F-K & S3).

**Fig 1.**
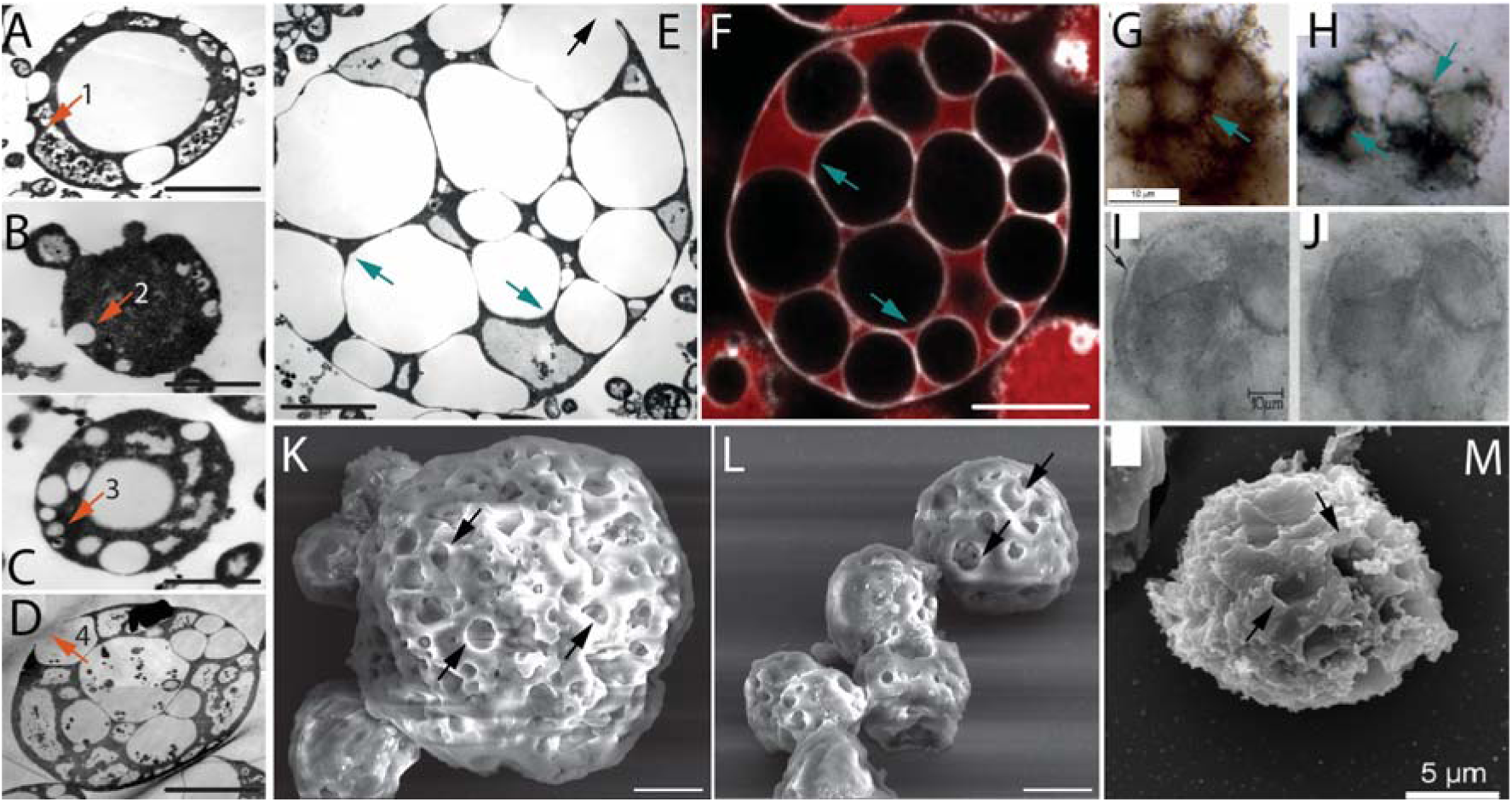
Morphological comparison of the Apex Chert and the Strelley Pool Formation microfossils with *EM-P*. Images A-D are TEM images of *EM-P* cells forming intracellular vesicles (ICVs) and intracellular daughter cells. The numbered arrows in these images point to different stages of ICV formation (see Fig. S1). Images E, F, K & L show TEM, SEM, and STED microscope images of *EM-P* cells with ICVs and surface depressions (black arrows). Cells in image F were stained with universal membrane stain, FM^TM^5-95 (red), and DNA stain, PicoGreen (green). Images G-J & M are spherical microfossils reported from the Apex Chert and the Strelley Pool Formation, respectively (originally published by Schopf et al., 1987 & Delarue *et al.,* 2019)(35,36). Cyan arrows in images E-H point to cytoplasm sandwiched between large hollow vesicles. The arrow in the image I point to the dual membrane enclosing the microfossil. Morphologically similar images of *EM-P* cells are shown in Fig. S3. Black arrows in images K-M point to surface depressions in both *EM-P* and the Strelley Pool Formation microfossils, possibly formed by the rupture of ICV’s as shown in D & E (arrows) (also see Fig. S4-S6). Scale bars: A-D (0.5µm) E, K & L (2 µm), and 5 µm (F).

The three-dimensional STED and SEM images of *EM-P* show numerous surface depressions (Fig. 1K, arrow & S1E). Such depressions are formed either during vesicle formation (Fig. 1A & 1B, arrow) or by the rupture of intracellular vesicles attached to the cell membrane (Fig. 1D, 1E, S1G-K & S4-S6). *EM-P* cells with such surface depressions and intracellular vesicles strongly resemble morphological resemblance to microfossils reported from 3.4 Ga Strelley Pool Formation (SPF) microfossils (36) (Fig. 1M & Fig. S4-S6). Microfossils reported from other sites, such as the Farrel Quarzite (Fig. S7)(38), Turee Creek (Fig. S8)(39), and the Fig Tree Formations (Fig. S9)(40), likely are morphological variants of Archaean *EM-P*-like cells and the Apex Chert microfossils. For instance, *EM-P* with many but relatively smaller intracellular vesicles resemble the Fig Tree microfossils, both in cell size and shape (Fig. S9). On the other hand, *EM-P,* with a relatively larger number of intracellular vesicles squeezed into polygonal shapes, resemble microfossils reported from the SPF (Fig. 1G-I & S4-S6), the Farrel Quartzite microfossils with polygonal alveolar structures (Fig. S7) and the Turee creek microfossils (Fig. S8)(38).

The second step in this method of reproduction involves the formation of daughter cells into the ICVs (Fig. S10)(34). Daughter cells were formed in the intracellular vesicles by a process resembling budding (Fig S10). Over time, these bud-like daughter cells detached from the vesicle wall and were released into the ICV (Movie 1). Due to a gradual loss of cytoplasm to the daughter cells, we observed a gradual reduction of the cytoplasmic volume of the parent *EM-P* cells and a corresponding increase in the number of daughter cells within the ICVs (Fig. S11, Movie 2-4). Over time, *EM-P* cells transformed into hollow vesicles with multiple tiny daughter cells (Fig. S11e-h, Movie 3&4). These intracellular daughter cells were released into the surroundings by a two-step process. In the first step, cells underwent lysis to release the ICVs (Fig. 2, S12 A & B and Movies. 5-9). In the second step, the vesicle membrane underwent lysis (Fig. S12C-E) to release the daughter cells (Fig. S12F-H). *EM-P* cells undergoing this method of reproduction closely resemble fossilized microbial cells discovered from several Archaean Eon rock formations (Fig. 2 & S13-S23)(36,41,42). For instance, microfossils reported from Mt. Goldsworthy Formation exhibit cells with ICVs containing daughter cells, cells that underwent lysis to release these ICVs (Fig. 2 & S13-S17), and the subsequent rupture of the vesicle membrane and release of the daughter cells (Fig. S14D-F)(34).

**Fig 2.**
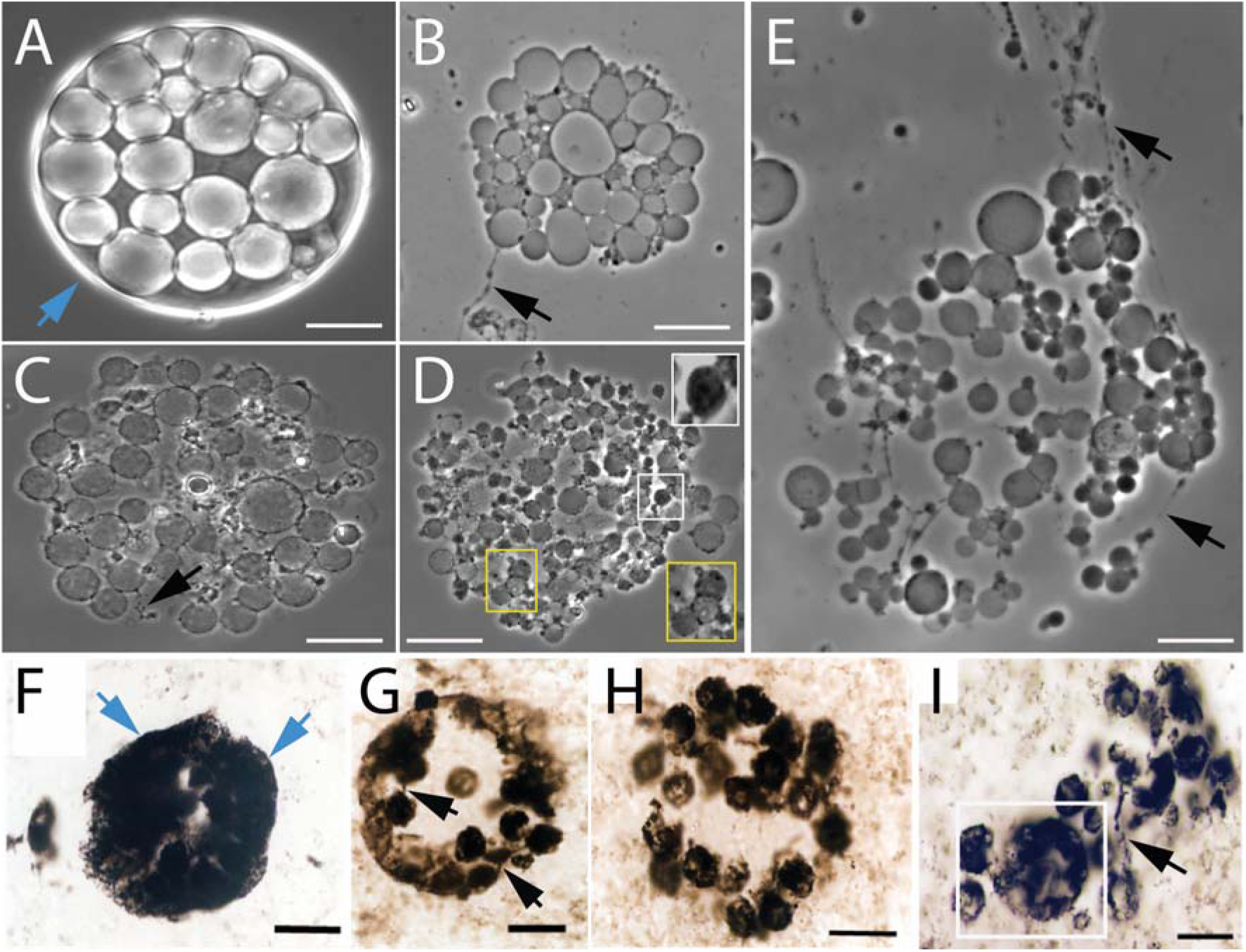
Morphological comparison between the Mt. Goldsworthy microfossils and *EM-P*. Images A-E show the process of cell lysis and release of intracellular vesicles in *EM-P*. Image A shows an intact cell with intracellular vesicles. Images B-E show lysis and gradual dispersion of these vesicles. Insert in image D shows enlarged images of individual ICVs. Images F-I show spherical microfossils reported from the Mt. Goldsworthy formation (originally published by Sugitani et al., 2009)(41). The arrow in this image, A & F, points to a cell surrounded by an intact membrane. The black arrow in these images points to filamentous extensions connecting individual vesicles. The boxed region in images D & I highlights a similar discontinuous distribution of organic carbon in ICVs and microfossils. Also see Fig. S13-S17. Scale bars: 20μm (A-E) & 20μm (F-I).

The microfossils reported from sites like the Farrel Quartzite (Fig. S7)(38), the Strelley Pool Formation (Fig. S18-S20), the Waterfall locality (Fig. S21-S22)(41,44,45), the Turee Creek (Fig. S23)(42), and Dresser Formation (Fig. S24-S28)(10), bear close morphological resemblance with morphologies of *EM-P* cells reproducing by this process. *EM-P* cells exhibited all the distinctive features of the Dresser formation microfossils, like the presence of hollow regions within the cell (Fig.S25) and discontinuous or thick-porous cell walls (Fig. S24 & S27). The step-by-step transformation of *EM-P* cells into these morphologies is shown in Fig. S27. Due to the lysis and release of daughter cells, most late stationary growth phase *EM-P* cells were deflated with numerous surface depressions. The morphology and surface texture of such deflated *EM-P* cells resemble the morphologies of microfossils reported from the Kromberg Formations (Fig. S29)(46).

Reproduction by external daughter cells happens by two different processes. Tiny daughter cells initially appeared as buds attached to the cell membrane (Fig. S30, S31 & Movie 10). These buds subsequently grew in size and detached from the parent cell. Depending on the size of the daughter cells (buds)*, EM-P* cells appear to have been reproducing either by budding or binary fission. *EM-P* cells that appear to have been reproducing by budding resemble microfossils reported from the North-pole formations (Fig. S31). As observed in our incubations, microfossils from this site are a mix of individual spherical cells, spherical cells with pustular protuberances, cells with bud-like structures, and cells undergoing binary fission (Fig. S31). Other *EM-P* morphotypes, like individual spherical cells, hourglass-shaped cells undergoing fission, and cells in dyads, bear close morphological resemblance to microfossils reported from both the Swartkopie (Fig. S32) and the Sheba formations (Fig. S33 & S34)(12,33,48). Like the Sheba Formation microfossils, the cells undergoing binary fission were observed to be in close contact with extracellular organic carbon (clasts of organic carbon in the Sheba Formation). This extracellular organic carbon likely represents the intracellular constituents released during the lysis of cells, as described above (Movies 5-8 & Fig. S34).

In some cases, the above-described buds did not detach from the cell surface but transformed into long tentacles (Movies 11&12). These initially hollow tentacles (34) gradually received cytoplasm from the parent cells and gradually transformed into “string-of-spherical daughter cells” (Fig. 3, S31A-C & S33)(34). Subsequently, these filaments detached from the parent cell, and due to the constant motion of daughter cells within these filaments, the “string-of-spherical daughter cells” fragmented into smaller and smaller strings and ultimately into multiple individual daughter cells (Movies 13-15). Apart from the cells that received cytoplasm (daughter cells), we also observed hollow spherical structures within these tentacles that did not receive cytoplasm from the parent cells (Fig. 3F & 3G, black arrows).

**Fig 3.**
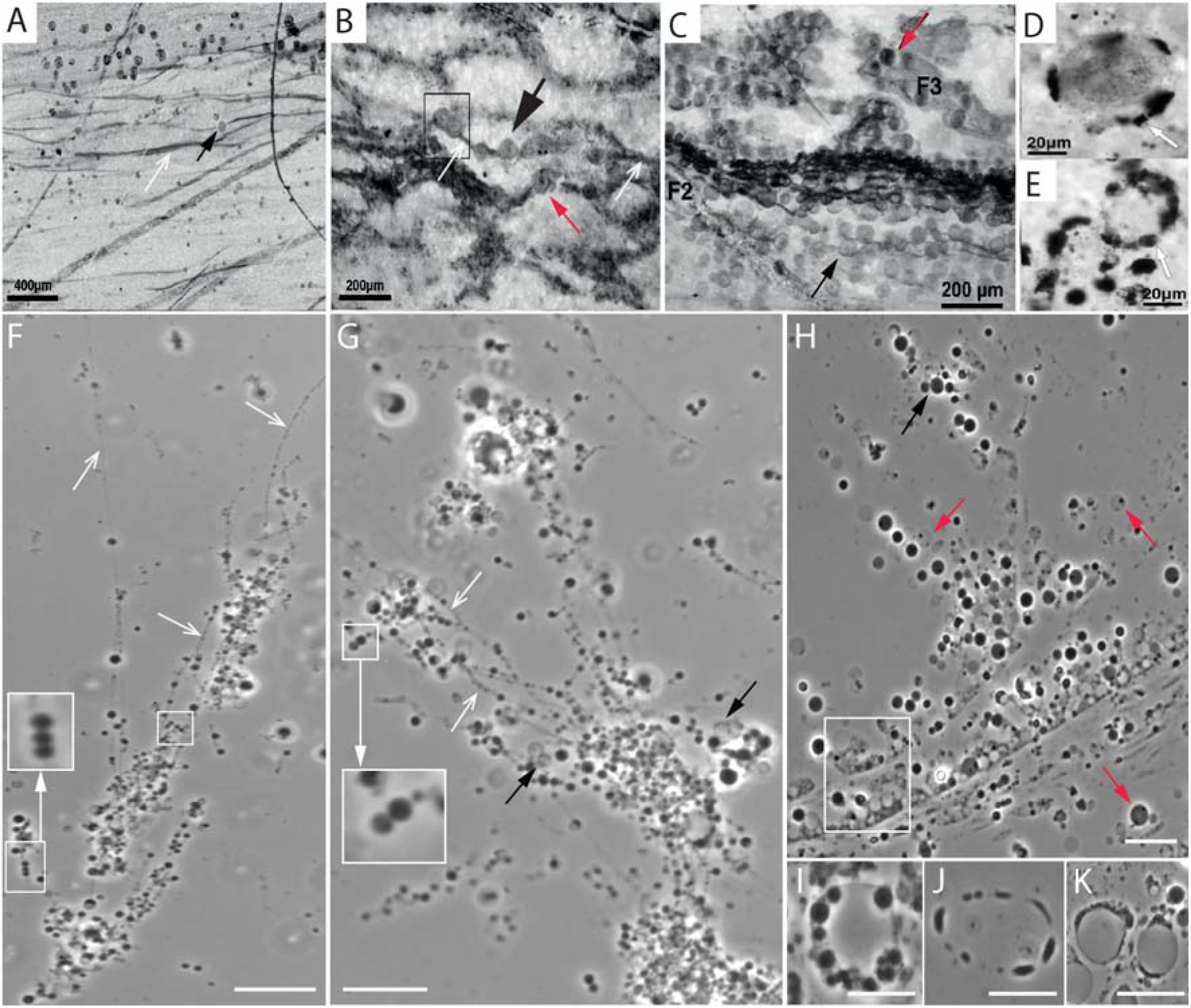
Morphological comparison of the Cleaverville microfossils with *EM-P*: Images A-E are the microfossils reported from Cleverville formation (originally reported by Ueno *et al.,* 2006). Images F-K are the *EM-P* cells morphologically analogous to the Cleverville Formation microfossils. Open arrows in images A, B, F & G point to the membrane tethers connecting the spherical cells within the filamentous extensions. Red arrows in the images point to the cells that have a similar distribution of organic carbon within the cells. Boxed and magnified regions in images B, F & G highlight the arrangement of cells in the filaments in pairs. The boxed region in image H highlights the cluster of hollow vesicles in *EM-P* incubations similar to the hollow organic structures in the Cleverville Formation, as shown in image C. Images D, E, and I-J show spherical cells that were largely hollow with organic carbon (cytoplasm) restricted to discontinuous patches at the periphery of the cell. Scale bars: 20μm (F-K).

All *EM-P* morphotypes observed undergoing this reproduction process bear close morphological resemblance to microfossils reported from the Cleaverville Formation (Fig. 3). All distinctive features of the Cleaverville microfossils, like the arrangement of cells as pairs within a string (Fig. 3H, 3F & 3G), microfossils with a discontinuous layer of organic carbon at the cell periphery (Fig. 3E, 3D & 3I-K), were also observed in *EM-P*. Several hollow spherical structures devoid of organic carbon were reported from the Cleaverville formation (Fig. 3). As in *EM-P*, these structures could have been the hollow membranous structures that didn’t receive the cytoplasm from the parent cells. Similar structures were also reported from other microfossil sites, such as the organic structures reported from the Onverwacht Group (Fig. S35)(6).

In addition to the Cleaverville microfossils, *EM-P* cells in our incubations also resemble filamentous structures with spherical inclusions reported from the Sulphur Spring Formations (Fig. S36-S38)(17). Based on the morphological similarities, we propose that these structures could have been the filamentous extensions with spherical daughter cells observed in *EM-P*. Similar but smaller filamentous structures were reported from the Mt. Grant Formation (Fig. S39)(39) and could have been the shorter fragments of similar “strings-of-daughter cells” (Movie 14). The spherical structures with sparsely distributed organic carbon reported from the Sheba Formations could also have been such cells undergoing fragmentation into smaller and smaller filaments (Fig. S33). In tune with this proposition, the Sheba Formation microfossils, like the *EM-P* cells, exhibited an un-uniform distribution of organic carbon within the cells and filamentous overhangs (Fig. S33, Movie 16).

Five to seven days after the start of the experiment, most cells in the incubations were the daughter cells. We observed three distinct types of daughter cells – a string of daughter cells (Fig. 3, S39 & Movie 14), daughter cells that were still attached to the membrane debris of the parent cell (Fig. 4, 5, S40-S48 & Movie 17) and individual daughter cells (Fig. 3h & Movie 15). All these daughter cell morphotypes resemble – a cluster of tiny spherical globules reported from the SPF (Fig. S18), a string of daughter cells reported from Mt. Grant formation (Fig. S39)(39,41), and spherical daughter cells still attached to membrane debris of parent cell like the ones reported from the Sulphur Spring Formations, Mt.Goldsworthy, the Farrel Quartzite, the Moodies Group, the Dresser Formation, and the SPF (Fig. 4 & S40-S48)(10,41,43,45,51).

**Fig. 4.**
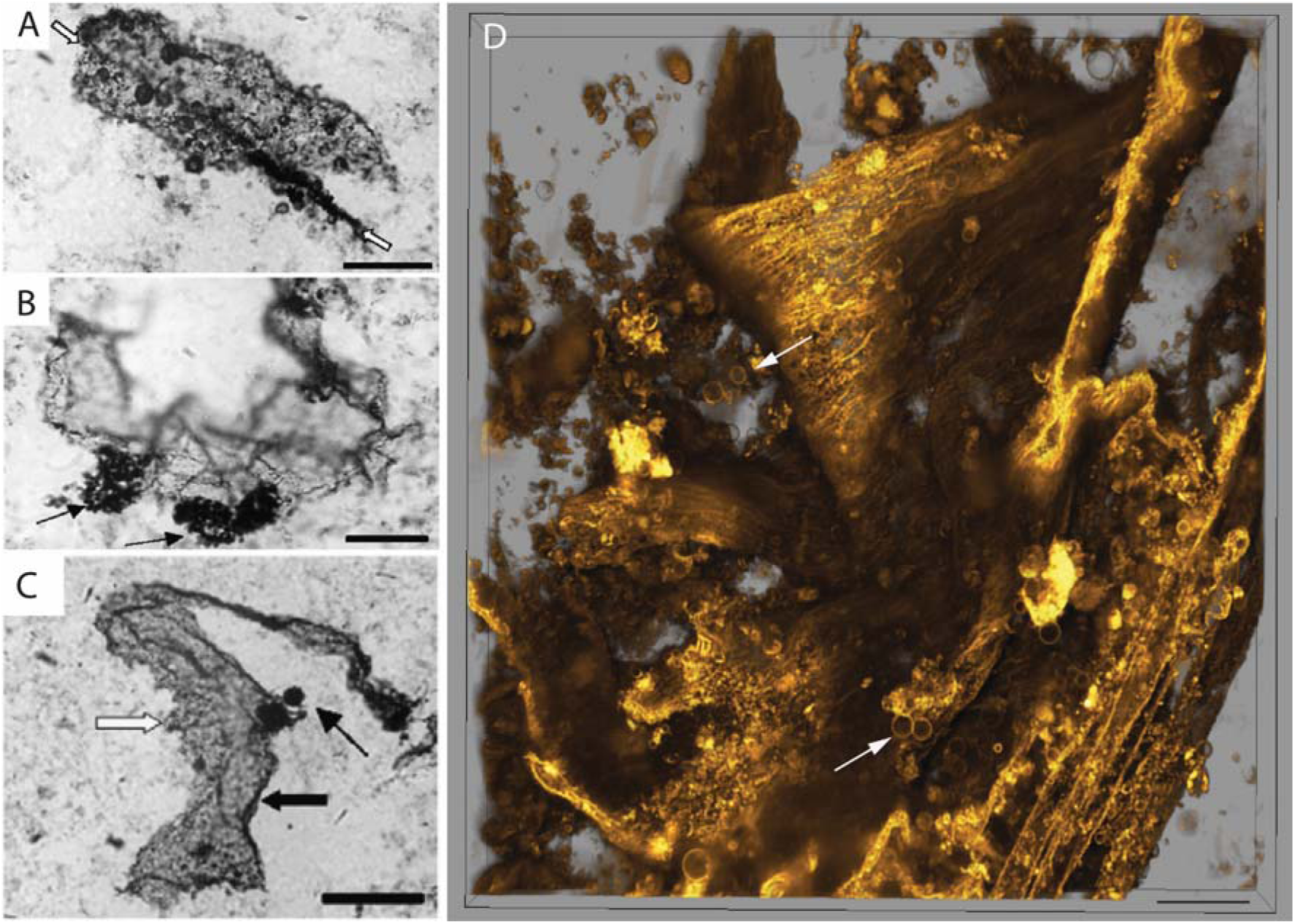
Morphological comparison between *EM-P* and the Mt. Goldsworthy microfossils. Images A, B & C are organic structures reported from the Mt. Goldsworthy Formation (Sugitani *et al.,* 2009)(43). Image D shows morphologically analogous film-like membrane debris observed in *EM-P* incubations. Arrows in images A-D point to either clusters or individual spherical structures attached to these film-like structures. Scale bar: 50μm (A-C) & 10μm (D).

**Fig. 5.**
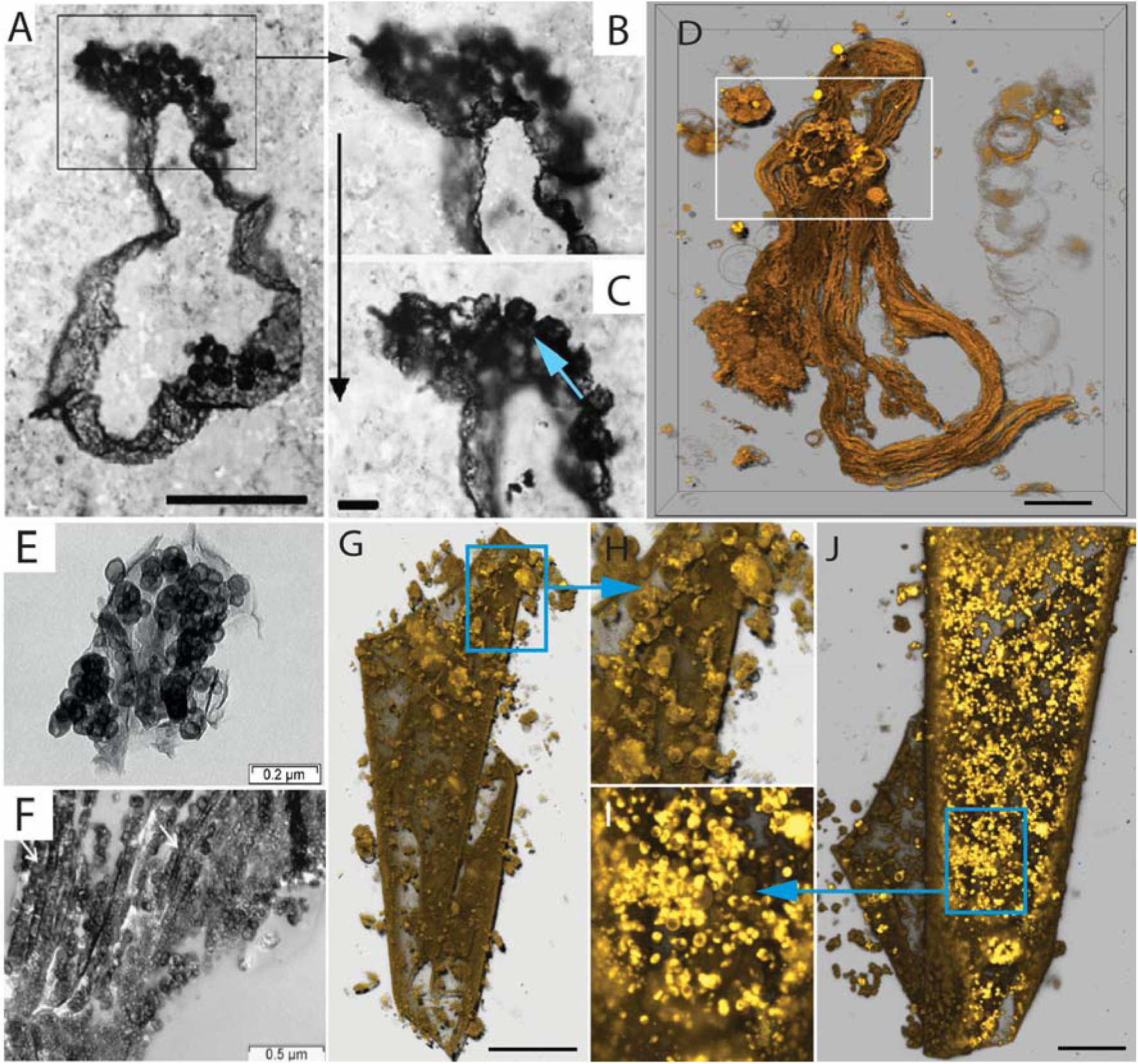
Morphological comparison of the Mt. Goldsworthy and the Sulphur Spring microfossils with *EM-P.* Image A-C are microfossils reported from the Mt. Goldsworthy Formation (39)(Sugitani *et al.,* 2007). Image D is the 3D-rendered STED microscope images of morphologically analogous membrane debris of *EM-P* cell with attached daughter cells (highlighted region) (also see Fig. S40). Images E & F are microfossils reported from the Sulphur Spring site (59)(Duck *et al.,* 2007), showing spherical structures attached to membrane debris. Images G & J are the morphologically analogous structures observed in *EM-P* incubations. Images H & I show the magnified regions of G & J showing spherical *EM-P* daughter cells attached to membrane debris (also see Fig. S40-S47, Movie 17). Cells and membrane debris in these images were stained with the membrane stain FM^TM^5-95 (yellow). Scale bars: A (50μm), G & J (20μm).

### The gradual transformation of *EM-P* cells into lamination-like structures and their comparison with Archaean organic structures

Fossilization and preservation of individual cells is considered unlikely due to the absence of rigid structures. However, recent studies indicate such a process could happen under favorable environmental conditions (52,53). However, the prevailing observations suggest that a significant portion of cell biomass undergoes taphonomic alteration and is preserved as degraded organic matter. To understand the possibility of *EM-P*-like cells forming structures similar to those observed in Archaean rocks, we studied the morphological transformation of individual *EM-P* cells and biofilms over 12 to 30 months. Below, we present the step-by-step transformation of individual cells into organic structures and their morphological resemblance to Archaean organic structures.

*EM-P* grew in our incubations as a biofilm at the bottom of the culture flask (Fig. 6 & Fig. S49). The rapid biofilm formation by *EM-P* can be attributed to the presence of extracellular DNA released during the lysis of *EM-P* cells (Fig. S50)(34). DNA released by such processes is known to promote biofilm formation (54). Over time, increased cell numbers resulted in biofilms comprising multiple layers of closely packed individual spherical cells (Fig. 6 & S50). A subsequent increase in the number of cells led to lateral compression and transformation of spherical cells into a polygonal shape (Fig. 6). By the late stationary growth phase, most cells underwent such a transformation, resulting in a honeycomb-like biofilm. The step-by-step transformation of individual spherical cells into these structures is shown in Fig. 6. Morphologically similar organic structures were reported from several paleo-Archaean sites, like the North Pole Formation (Fig. S51-S62)(47). A similar taphonomic degradation of organic matter associated with a biofilm was demonstrated by previous studies (Westall et al., 2006, Figure 10B)(55).

**Fig 6:**
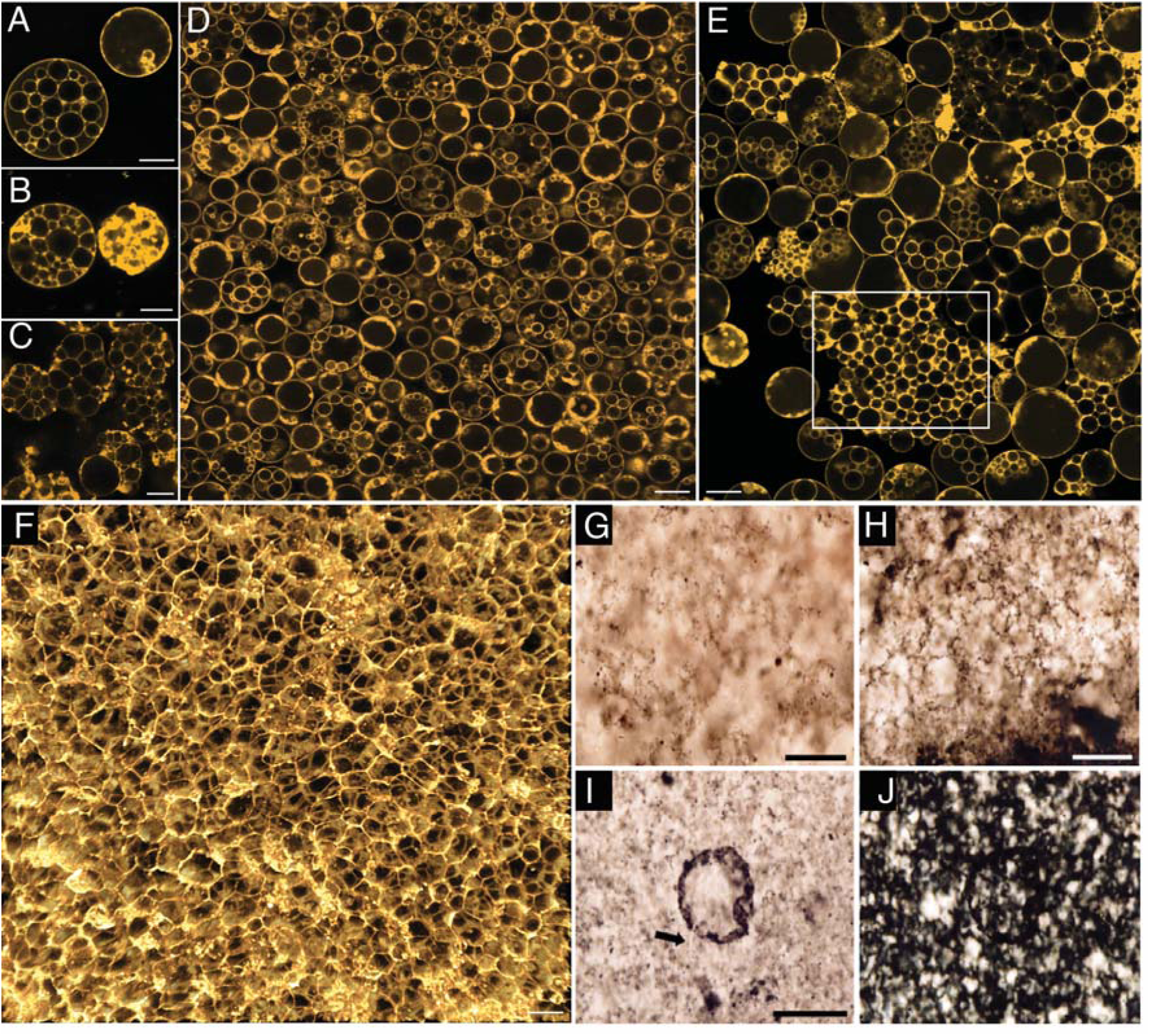
Sequential steps involved in the formation of honeycomb-shaped mats: Images A-C show single *EM-P* cells that gradually transformed from spherical cells with intracellular vesicles into honeycomb-like structures. Images D-E show a similar transformation of biofilms composed of individual spherical cells into honeycomb-like structures. Cells in these images are stained with membrane stain, FM^TM^5-95 (red), and imaged using a STED microscope. Images G-J are the microfossils reported from the SPF (originally published by Sugitani et al., 2007)(39). Scale bars: A-F (10μm), G & H (20μm), and I (50μm).

Large aggregations of spherical cells devoid of internal organic carbon were reported from the North-Pole Formation (47) (Fig S52). These structures closely resemble the aggregations of hollow ICVs released after the lysis of *EM-P* cells (Fig. 2A-2E & S22). As observed in *EM-P*, the distribution of organic carbon in the North-Pole Formation microfossils is restricted to the periphery of the spherical cells (Fig. S52). Along with the morphological and organizational similarities, *EM-P* also exhibited all the accessory structures associated with the North Pole formation microfossils, such as the large clots of organic carbon (Fig. S51 arrows), filamentous structures originating from the spherical cells, and spherical clots of organic carbon within these filamentous (Fig. S52-S55)(47). Based on the similarities, we propose that the large clots of organic carbon would have been the membrane debris formed during the lysis and the release of intracellular vesicles (Fig. S51, Movie 5-9). As observed in *EM-P*, the filamentous structures associated with the microfossils could also have formed during the release of intracellular vesicles (Movie 5-9). The organic carbon clots within the filamentous structures could have been fossilized daughter cells (Fig. S53 & S54, arrows).

Apart from the aggregations of hollow spherical cells, honeycomb-like structures were also reported from several microfossil sites, like the SPF, the Nuga Formation, the Buck Reef Chert, the Moodies Group, and the Turee Creek formations (Fig. 6, S57-S62)(42,56–58). As observed in *EM-P*, these structures could have been formed by the lateral compression of cells or hollow vesicles within the biofilm (Fig. 6). In tune with our proposition, Archaean honeycomb-like structures are often closely associated with spherical *EM-P*-like cells (Fig. 6).

Spherical microfossils from the Pilbara and Barberton Greenstone Belts were often discovered within layers of organic carbon (48,51,60,61). Over a period of 2-6 months, we observed cells in our incubations gradually being enclosed by membrane debris. These structures were formed by a multi-step process (Fig S63). First, *EM-P* grew as multiple layers of cells within a biofilm (Fig. S63A). Second, the lysis of these cells led to the formation of a considerable amount of membrane debris (Fig. S63B, S64 & Movie 17&18). Subsequently, this membrane debris coalesced to form large fabric-like structures (Fig. S65). These membrane fabrics were then expelled from the biofilm (Fig. S63D, S63E & Movie 18). Over time, these expelled membrane fabrics grew in surface area to form a continuous layer of membrane enclosing a large population of cells (Fig. S65-S69 & Movie 19). This fabric-like membrane debris enclosing biofilms observed in *EM-P* incubations bear close morphological resemblance to microfossils reported from Chinaman Creek in the Pilbara (Fig. S69), and Mt. Goldsworthy Formation (Fig. 4 & 5)(39,43,62).

Parallel layers of organic carbon termed laminations were reported from several Archaean microfossil sites (33,51,60,63). Structures similar to these laminations were observed in our incubations. As described above, the reproduction in *EM-P* involves the lysis of cells to facilitate the release of the intracellular daughter cells, resulting in a considerable amount of cell debris (Movie 9 & 17). The parallel layers of organic carbon in our incubations (Fig. 7 & 8) are formed by lysis and collapse within individual biofilm layers (Fig. S49). Another way the organic carbon layers could have formed is by the lateral compression of honeycomb-like biofilms (Fig. S71-S74). Sequential steps resulting in the formation of such structures are shown in Fig. S72. Such layers of cell debris closely resemble different types of laminated structures reported from the Barberton Greenstone Belt and Pilbara Iron Formations, like the α, and β laminations (Fig. 7, 8 & S69-S75)(51,60,61).

**Fig. 7:**
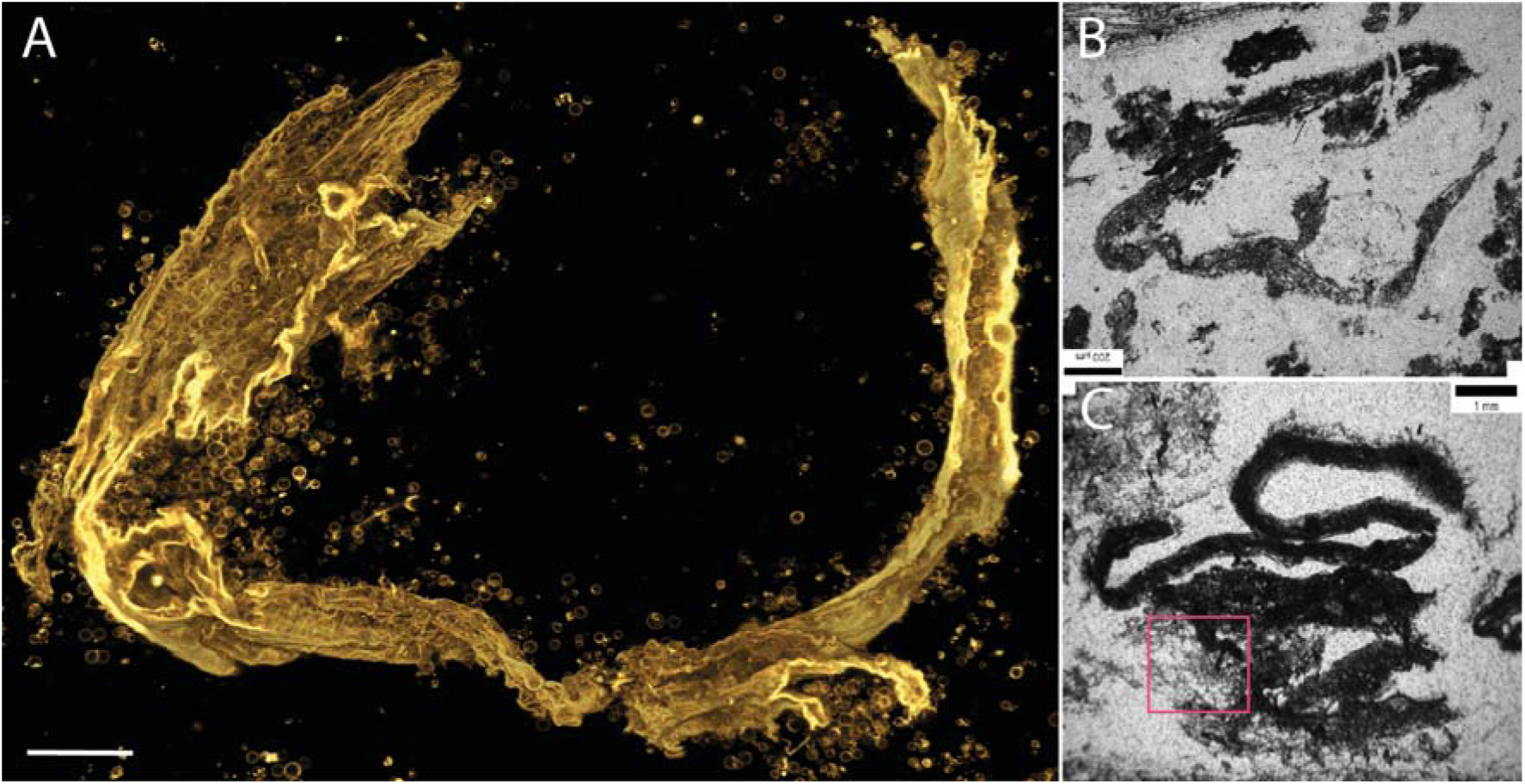
Morphological comparison of the Buck Reef Chert. β**-laminations with *EM-P’s* membrane debris.** Image A shows a 3D-rendered image of *EM-P’s* membrane debris. Cells in the image are stained with membrane stain Nile red and imaged using a STED microscope. Images B & C show β-type laminations reported from Buck Reef Chert (originally published by Tice *et al.,* 2009)(61). The boxed region in image-a highlights the membrane-forming rolled-up structures containing spherical daughter cells, as described in the case of BRC organic structures. Scale bars: 50μm.

**Fig. 8:**
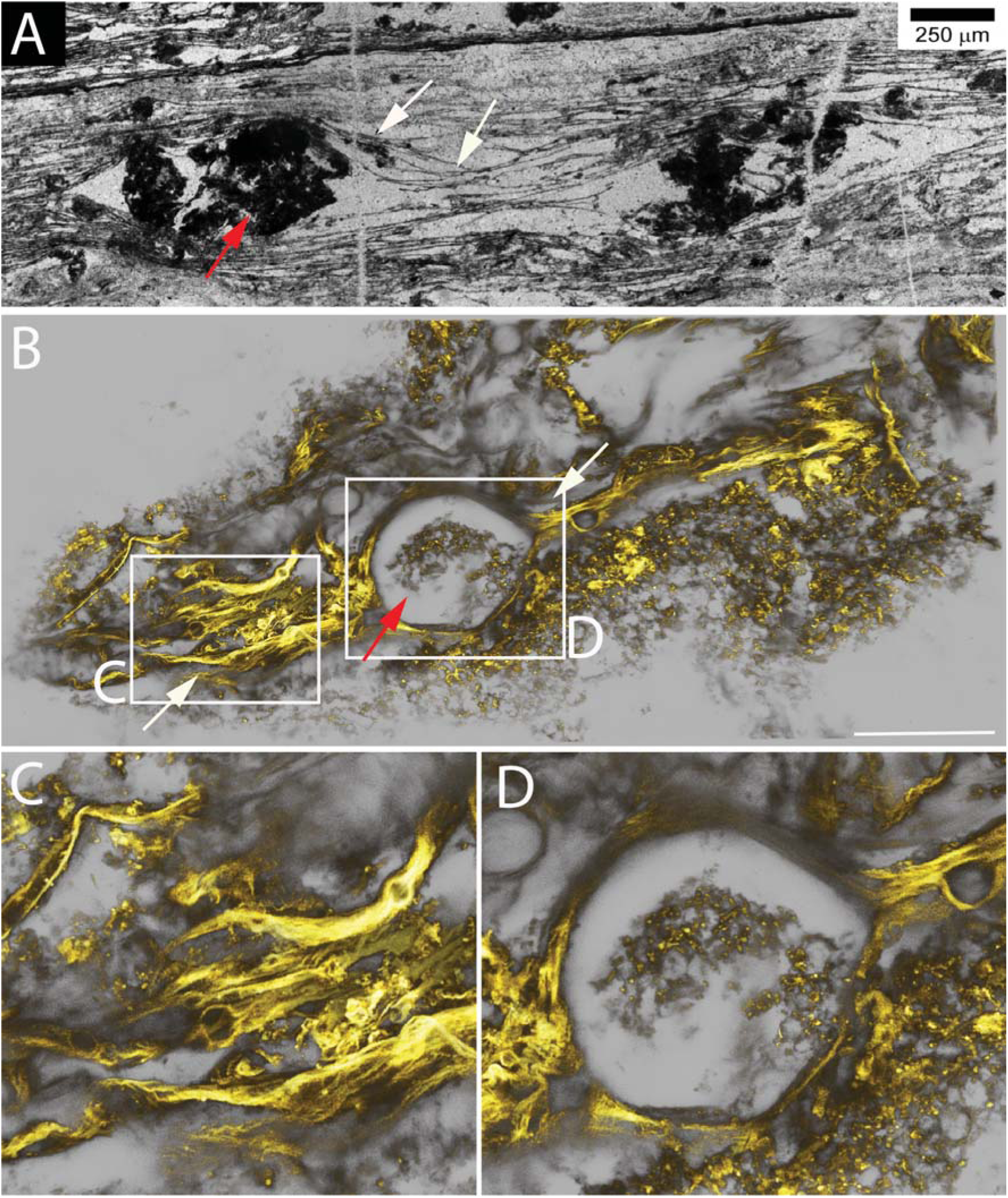
Morphological comparison between laminated structures reported from the Moodies Group and structures formed by *EM-P.* Image A shows laminated structures reported from the Moodies Group (originally published by Homann *et al.,* 2015)(63). They show parallel layers of organic carbon with lenticular gaps. Together with the quartz, these lenticular gaps consist of clumps of organic carbon. Image B is a 3D-rendered confocal image of analogous membrane debris formed by *EM-P*. Images C & D are the magnified regions of C. Like Moodies formation, filamentous membrane debris bifurcating to form spherical\lenticular gaps can be seen in several regions (S75, S76, & S77). Some spherical/lenticular gaps were hollow, and some had an organic structure within them, even exhibiting a honeycomb pattern (arrow), suggesting the presence of large spherical *EM-P* cells with intracellular vesicles (D, & S77). Membranes were stained with Nile red, and imaging was done using a STED microscope. The scales: 50μm.

Also similar to the Archaean laminations, we observed layers of cell debris in our incubations have lenticular gaps (Fig. 8, S69, S70-S71 & S76-S79). Within these lenticular gaps, we observed intact *EM-P* cells or honeycomb patterns, suggesting that lenticular gaps within otherwise uniformly parallel laminations were formed due to non-uniform lysis or incomplete deflation of cells within individual layers of *EM-P* cells (Fig. 7, S75-S79 & Movies 20 & 21). Although in the case of Archaean laminations, these lenticular gaps were thought to have been formed by the entrapment of air bubbles (51), based on our results, we argue that there could have been more than one way such structures could have formed. Other distinctive features of the lamination, like raised mounds or swirls (Fig. S81 - S83)(51,61), were also observed in batch cultures of *EM-P*. Given these morphological similarities, we propose that some of the laminated and other diaphanous filamentous structures could have been formed by the cell debris of the *EM-P*-like cells that inhabited these sites during the Archaean Eon. We will discuss these possibilities in more detail below.

Over a period of 3-12 months, we observed the biofilms solidifying into a solid crust (Fig. S84 & S85). The SEM-EDX characterization of these solidified biofilms showed the presence of potassium and magnesium minerals on the surface, suggesting that these structures were formed by the gradual adsorption of positively charged cations on the negatively charged biofilms (Fig. S84). Most Archaean microfossils are restricted to coastal marine environments. Compared to open oceans, these coastal marine environments harbor higher concentrations of salt due to higher evaporation rates. Hence, these microfossils could have undergone a similar encrustation process as observed in our incubations. Moreover, solidified *EM-P* biofilms resemble the mineral-encrusted structures reported from the Kromberg Formation (Fig. S85)(64). Like the Kromberg Formations structures, solidified *EM-P* biofilms are composed of desiccation cracks and spherical cells beneath the surface (Fig. S85).

When these salt-encrusted cells were transferred into fresh media, we observed a gradual increase in cell numbers (results not shown). However, given that these cells are encrusted in a layer of salts, we observed early growth-phase cells breaking out of the thick salt crust, resulting in stellar morphologies (Fig. S86 & S87). The observed morphologies of these cells closely resemble the morphologies of microfossils reported from the Strelley Pool and other North Pole cherts (Fig. S86 & S87)(41,47). All distinctive features of these microfossils, like the stellar-shaped cells undergoing binary fission and a string of daughter cells extending out of such stellar cells (Movie 22), were also observed in our incubations.

## Discussion

Advances in microscopic (FIB-SEM) and analytical (NanoSIMS) techniques over the past few decades have facilitated better imaging and precise determination of chemical and isotopic compositions of microfossils (13,65). Nevertheless, there is considerable disagreement among researchers regarding the interpretation of this information (9). Given the importance of morphology in determining the biogenicity and taxonomic affiliation of the microfossils, reconstructing the lifecycles of Archaean Eon organisms is considered crucial to understanding the nature of these microfossils (43). Our study is the first to reconstruct all known spherical microfossil morphologies and their lifecycles from extant bacteria. Furthermore, we have shown that many of the taphonomic structures observed in our study closely resemble the controversial structures observed in rocks of the Palaeo-Mesoarchaean age (3.6-3.0 Ga) and even in the Neoarchaean (3.0-2.4 Ga). These similarities help us answer long-standing questions regarding the origin and the nature of Archaean microfossils.

The nature of Archaean organic structures is currently being debated among researchers (7,9,10). While some studies suggest that these structures could be remnants of Archaean microorganisms, others suggest that they may have been abiotic minerals that formed due to volcanic activity (10). The argument for this proposition is based on the fact that these organic structures share more similarities with inorganic mineral structures than with extant prokaryotes. To establish the biogenicity of a microfossil, it is essential to either find a convincing morphological analog among extant bacteria or establish a biogenic process through which they are formed (3). The biogenicity of microfossils reported from several sites like the Swartkoppoie Formation, Kitty’s Gap Chert, and the Josephdal Formation is widely accepted among the scientific community due to the discovery of spherical microfossils in different stages of their lifecycle (Fig. S32)(12,55,64). However, such a step-by-step biological process through which Archean Eon organic structures could have formed has never been demonstrated empirically.

The biological origin of microfossils reported from several sites, like the Dresser Formation, to date, remains a matter of debate (10). Several morphological features of these organic structures, like the presence of organic carbon only at the periphery, the absence of internal cell constituents, the presence of pyrite and silicate minerals inside the cells, and the presence of a thick porous or discontinuous cell wall, were all argued as claims for their abiotic origin (10). Justifiably, these morphological features have never been observed in any living organism. Nevertheless, all spherical microfossils reported from the Dresser Formation resemble *EM-P* cells, especially those with a single large ICV (Fig. S24-S27). What was thought to have been a thick, porous cell wall in the microfossils could have been the cytoplasm with tiny ICVs sandwiched between the cell and vesicle membrane (Fig. S24 & S27). Similarly, the hollow cells with a discontinuous cell wall could either have been the ICVs released by cell lysis or the late-growth stage cells with little cytoplasm (Fig. S26). In such cells, the presence of cytoplasm is restricted to discontinuous patches around the periphery of the cells (Fig. S26). The sequence of steps leading to these cell morphologies that resemble the Dresser Formation microfossils is shown in Fig. S27. A closer inspection of the Dresser Formation microfossils shows the ICVs membrane rupture and the daughter cell release (Fig. S28). Morphologies indicating this method of reproduction among microfossils is not unique to the Dresser Formation. Microfossils with similar morphological features were reported from sites like the Strelley Pool, the Waterfall region, and Mt. Goldsworthy Formation (Fig. S3-S14)(3,43). These similarities suggest that microfossil morphologies observed in the Dresser Formation are in tune with other microfossils of similar geological time periods, suggesting their biological origin.

Spherical structures half-coated with pyrite are reported from the Dresser Formation (Fig.S25)(10). These structures could have been the iron-reducing *EM-P*-like cells with hollow ICV constituting half their volume. The selective co-localization of pyrite and carbon could be explained by the Fe(III) reduction happening at the cell surface. The Fe(II) produced from this metabolic reaction could have reacted with environmental sulfide, converted to insoluble pyrite, and precipitated onto the cell surface. Given the absence of this metabolic process within the hollow ICV, these structures remained pyrite-free (Fig.S25). In addition to the Dresser Formation, organic structures coated with pyrite have also been reported from other microfossil sites like the Sulphur Spring Formations (17). The selective presence of pyrite on these microfossils could also be explained by a similar mechanism (Fig. S36-S38). Apart from pyrite, minerals like anatase were reported to have been present within the cells (10). The presence of anatase within these cells could be explained by the transport of these minerals into the cells during the ICV formation (Fig. 1B&D). ICVs are formed by a process similar to endocytosis, which involves the intake of salt-rich media and minerals into the cells (Fig. S88). Like the Dresser Formation microfossils, we often observed the presence of salts and minerals within *EM-P*s vesicles (Fig. S88). Moreover, the presence of minerals within cells is not unique to the Dresser Formation microfossils (10) and was reported previously from several bonafide microfossils from Gunflint Iron Formations (66). Additionally, we observed remarkable similarities between the *EM-P* cell debris and all the organic structures closely associated with the microfossil, such as the wavy lamellar and pumice-like structures (Fig. S59, S61 & S89). The step-by-step transformation of cell debris into pumice-like structures is shown in Fig. 6. Based on these morphological similarities between the Dresser Formation organic structures and *EM-P*, we hypothesize that these organic structures are the fossil remnants of *EM-P*-like bacteria rather than mineral aggregates.

Morphological similarity between microfossils from far-flung sites like Western Australia and Southern Africa could be explained by the similarity in the environmental conditions in both sites (5). This relationship between cell morphology, reproductive processes, and environmental conditions was discussed extensively in our previous work (34,67). The experimental conditions that we employed in our study are likely similar to the environmental conditions faced by Archaean organisms from both these sites at the time of their fossilization. All sites from which microfossils were reported are shallow intertidal regions. Evidence for periodic evaporation and flooding with sea water was presented from the Barberton and Pilbara Greenstone Belts (6,68), suggesting that the original microorganisms experienced high salinities. The salinities of our experiments are broadly similar to those of Archaean oceans (5-10% w/v)(31). To our knowledge, the exact salt composition of the Archaean Ocean has not been elucidated. Hence, we used a complex mixture of salts (DSS) as a proxy to reproduce these salinities in our experiments. Salts like Mg^+2^, Ca^+2^, Na^+,^ and K^+^ or their oxides were also reported to be present and constitute 1-5% by weight in both Pilbara and Barberton greenstone belt microfossil sites (6,68). Moreover, these salts were shown to be closely associated with microfossils (68). The spatial distribution of these salts resembles the spatial distribution pattern of organic carbon, possibly indicating the chelation of these salts to the cell membrane, which is also in agreement with our observations (9). The presence of potassium phyllosilicates and NaCl crystals within the microfossils (68) is also in agreement with our hypothesis that internal structures of the microfossils should have formed by invagination of cell-membrane taking in salt-rich water (Fig. S86). As observed in the microfossils (68), salt crystals on the cell surface, within the membrane invaginations, or cell debris were often observed in *EM-P* (Fig. S89).

The above-presented results suggest that Archaean Eon cells are likely primitive lipid-vesicle-like protocells that lack a cell wall. From a physiological perspective, it would have been unlikely for primitive cells to possess a cell wall given the substantial number of genes required to synthesize individual building blocks, to mediate its assembly, and its constant modification to facilitate cell growth and reproduction (21,69,70). Furthermore, a cell wall could impede the transport of physiologically relevant compounds in and out of the cells. To overcome this limitation, present-day microorganisms (with a cell wall) had to develop extensive molecular biological processes for transporting nutrients and metabolic end products across the cell wall (71,72). This could not have been the case for primitive Archaean life forms. Hence, rather than drawing parallels between the microfossils and life as we know it today, we propose that these microfossils could have been liposome-like protocells, as proposed by the theory of chemical evolution (26). Indeed, it has been recently shown that liposome-like molecules could be produced in some of the hydrothermal settings proposed for the emergence of life (73). To the best of our knowledge, this is the first study to provide a link between theoretical propositions and geological evidence for the existence of protocells on early Earth.

According to the theory of chemical evolution, biological organic compounds are formed by abiotic processes (74). These compounds then self-assembled to form lipid vesicles, which grew in complexity and eventually evolved into self-replicating protocells (75,76). These protocells are believed to have undergone Darwinian evolution, resulting in the emergence of bacteria, archaea, and eukaryotes (77). It was previously thought that the fragility of protocells made it unlikely for them to be preserved in rock formations. However, later studies showed the preservation of cellular features by a rapid encrustation of cells with cationic minerals (52,53). The rapid encrustation and preservation of cells observed in our study (Fig. S84-S85) is in accordance with the proposition that environmental conditions influence the extent of cellular preservation. Our study aligns with the interpretations from these studies that environmental conditions play a pivotal role in determining the extent of cellular preservation.

### Conclusion

For the first time, our investigations have been able to reproduce morphologies of most Archaean microfossils from wall-less, extant cells. Apart from reproducing the morphologies, we also presented a step-by-step biological process by which Archaean organic structures could have formed. Based on these results, we propose that Archaean microfossils were likely liposome-like cells, which had evolved mechanisms for energy conservation but not for regulating cell morphology and replication. In an earlier study, we have shown that the morphologies of such primitive cells are determined by environmental conditions (34,67) rather than the information encoded in their genome. Given this lack of intrinsic ability to regulate their morphology, we argue that morphological features such as cell size, shape, or cytological complexity are reliable factors in interpreting either the phylogeny or the physiology of microfossils (at least from Archaean Eon). Rather than attempting to assign present-day taxonomies to these microfossils, we suggest that these microfossils represent primitive protocells proposed by the theory of chemical evolution. To the best of our knowledge, ours is the first study to provide paleontological evidence of the possible existence of protocells on the Palaeoarchaean Earth.

## Methods

### Isolation of cells and their transformation to protoplasts

*Exiguobacterium* strain-Molly (*EM*) was isolated from submerged freshwater springs within the Dead Sea (78). The taxonomic identification of the isolate to the genus *Exiguobacterium* was determined by 16S rRNA gene sequencing (79,80). *EM* cells were transformed into protoplasts following a previously documented protocol(28). The resulting *EM-P* cells were cultured in half-strength TSB with 7% Dead Sea Salt (7%DSS-TSB).

### Microscopic observation of *EM-P* cells

Morphology *EM-P* was routinely assessed using an Axioskop 2plus microscope (Carl Zeiss, Germany) with a Plan-NEOFLUAR 100X/1.3 objective. Images were captured using a Leica DSF9000 camera (Leica Microsystems, Mannheim, Germany). STED microscopy was performed with an inverted TCS SP8 STED 3X microscope (Leica Microsystems, Mannheim Germany) using an 86x/1.2 NA water immersion objective (Leica HC PL APO CS2 -STED White). Fluorophores were excited with 488, 561nm, 594nm, or 633m laser light derived from an 80 MHz pulsed White Light Laser (Leica Microsystems, Mannheim Germany). For stimulated emission, either a pulsed 775 nm laser or a 592nm CW laser (Leica Microsystems, Mannheim, Germany) was used depending on the fluorophore. Photon counting mode and line accumulation were used for image recording, and Huygens Professional (SVI, Hilversum, The Netherlands) performed image deconvolution on selected images and movies.

Spinning Disk Microscopy was performed using an Olympus SpinSR10 spinning disk confocal microscope (Olympus, Tokyo, Japan) equipped with a 100x/NA1.35 silicone oil immersion objective (Olympus UPLSAPO100XS, Tokyo, Japan), a CSU-W1-Spinning Disk-Unit (Yokogawa, Tokyo, Japan) and ORCALFlash 4.0 V3 Digital CMOS Camera (Hamamatsu, Hamamatsu City, Japan).

Transmission electron microscopy was conducted utilizing a Zeiss EM 912 (Zeiss, Oberkochen, Germany) equipped with an integrated OMEGA filter, operating at 80 kilovolts (kV). Image acquisition was carried out using a 2k x 2k pixel slow-scan CCD camera (TRS, TrÖndle Restlichtverstrkersysteme, Moorenweis, Germany) with ImageSP software (SysProg, Minsk, Belarus).

## Acknowledgments

We want to thank Gabriella Berthal for her excellent technical support and Christian Sibert for providing the Dead Sea samples from which *EM* was isolated. We thank the Advanced Light Microscopy Facility at EMBL, Heidelberg, Ulf Schwartz from Leica Microsystems, and colleagues at the departments of Ecological Microbiology (Bayreuth University) and of Cellular and Molecular Biophysics (Max Planck Institute for Biochemistry) for their support throughout the work.

## Data Availability Statement

Dr. Dheeraj Kanaparthi will share all data, materials, and methods upon reasonable request.

