## Supplement File 1 for "On the nature of the earliest known lifeforms"

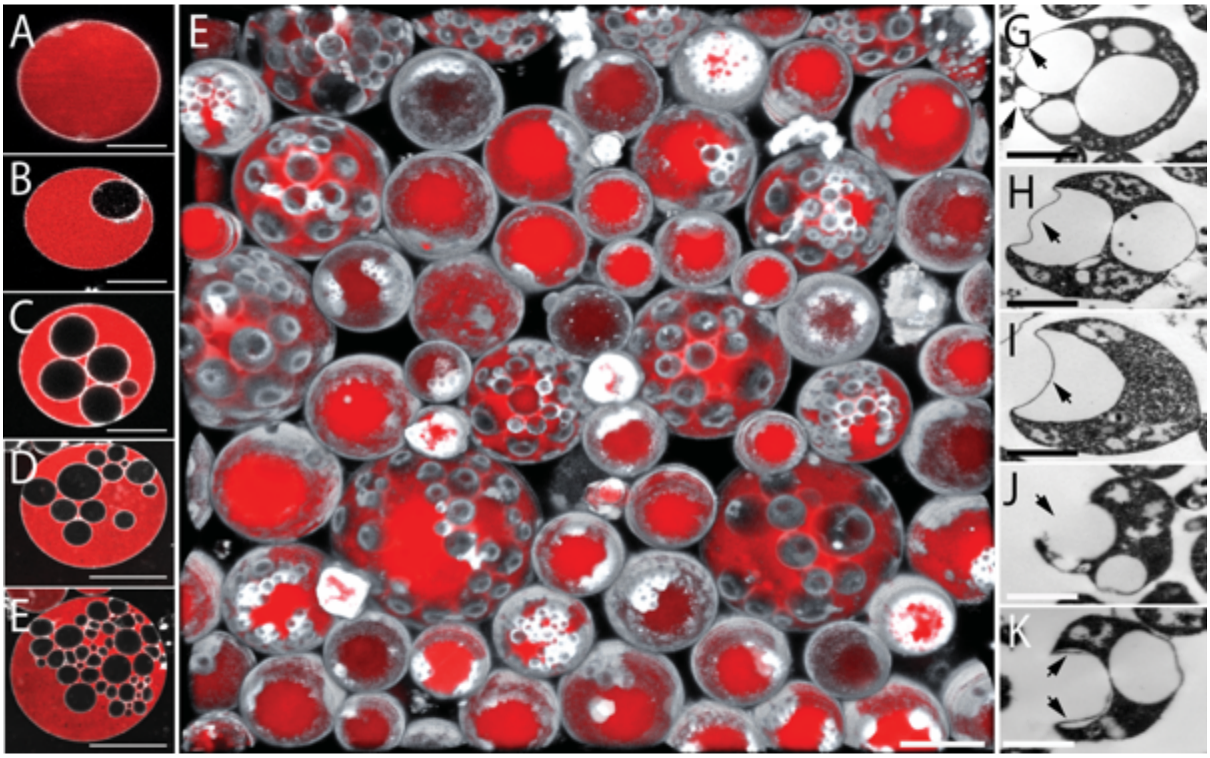


**Fig S1.** ***EM-P* cells with intracellular vesicles (ICV)**

Images A-E show STED microscope images of *EM-P* cells in different stages of intracellular vesicle formation. Cells in these images were stained with FM$^{TM}$5-95 (membrane, white) and PicoGreen$^{TM}$ (DNA, red). Image B shows the first stage of vesicle formation by a process similar to endocytosis. Images C-E show a gradual increase in the number of intracellular vesicles. Image-e shows the 3D-rendered STED microscope image of *EM-P* cells with hollow invaginations on the cell surface. Images G-K show TEM images of *EM-P* cells with intracellular vesicles. Arrows in these images point to the gradual collapse of ICVs, leading to the formation of spherical invaginations on the cell surface. Scale bars: 5µm (A-E), 10µm (F), and 250nm (G-K).

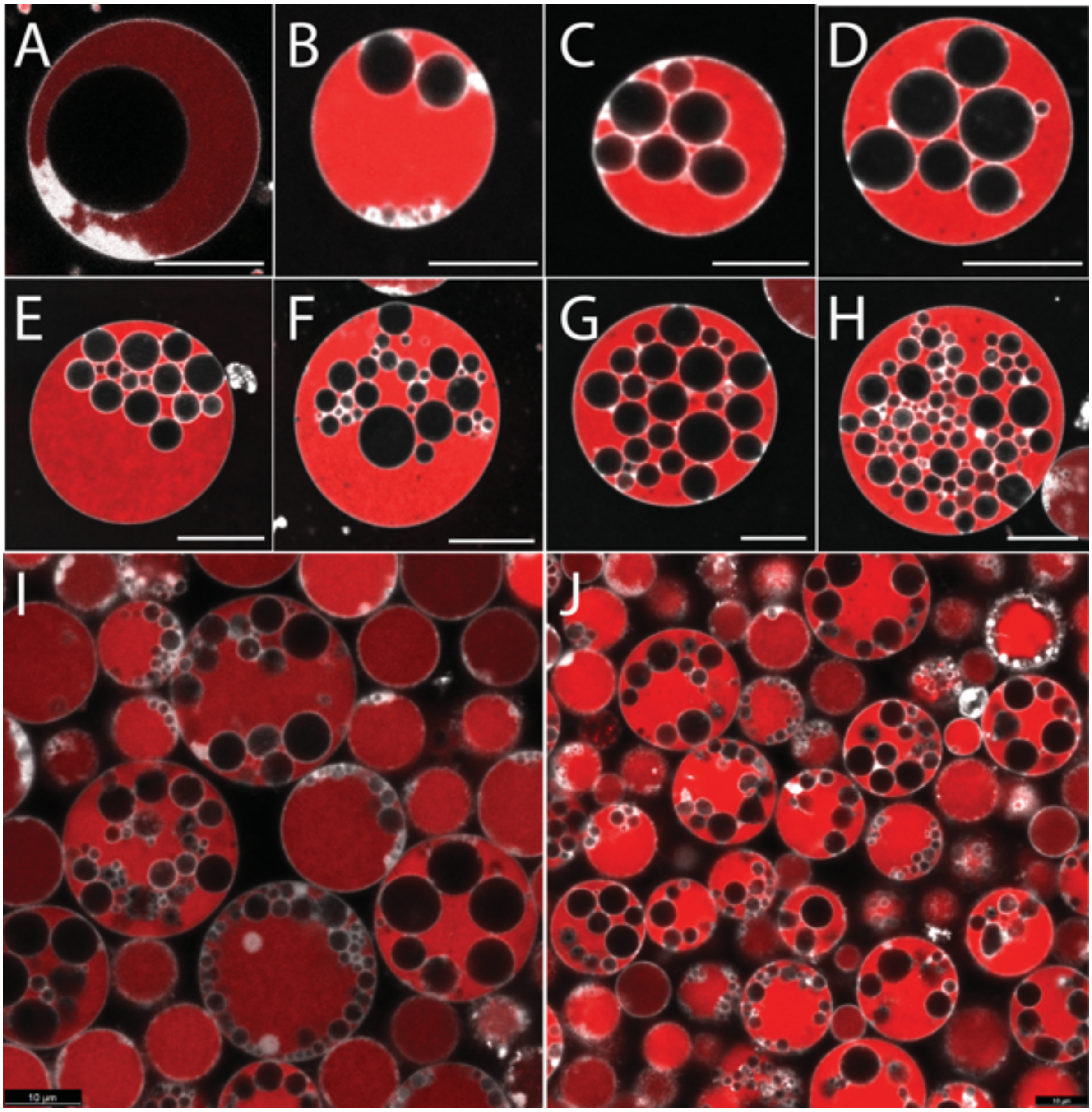


**Fig S2. Variation in the morphology of *EM-P* cells and intracellular vesicles**

Images A-J are STED microscopy images of *EM-P* cells with intracellular vesicles. These images show variations in the size and number of vacuoles observed within *EM-P*. Cells in these images were stained with FM™5-95 (membrane, white) and PicoGreen™ (DNA, red). Scale bars: 5µm (A-G) & 10µm (H-J).

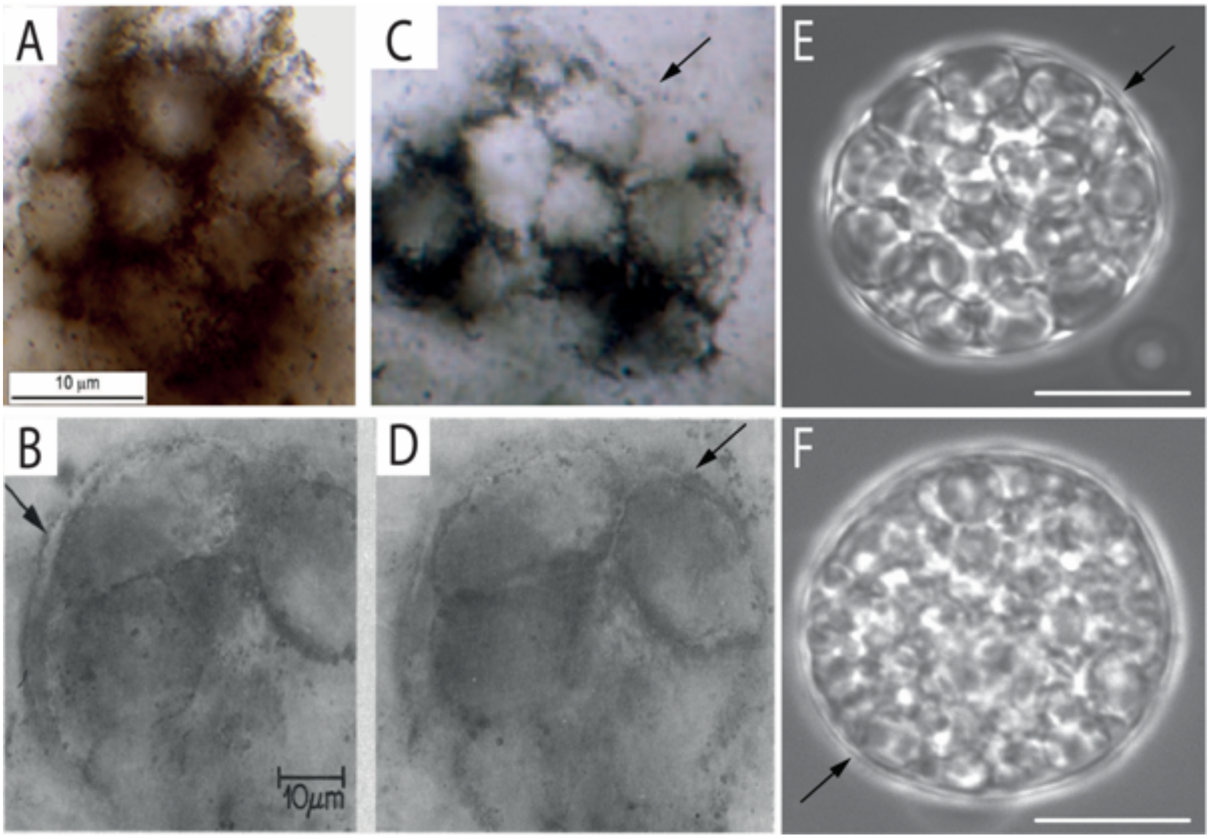


**Fig. S3. Morphological comparison of the Apex Chert microfossils with *EM-P*.**
Images A-D show spherical microfossils with hollow intracellular compartments and two layers of the outer cell membrane (arrows) (originally published by Schopf *et al.,* 1987)(34). Images E & F are morphologically analogous to *EM-P* cells with hollow intracellular compartments. Like microfossils, *EM-P* cells also appear to have a dual outer membrane due to the juxtaposition of vesicle membrane against cell membrane (arrows). Scale bar: 10µm (E & F).

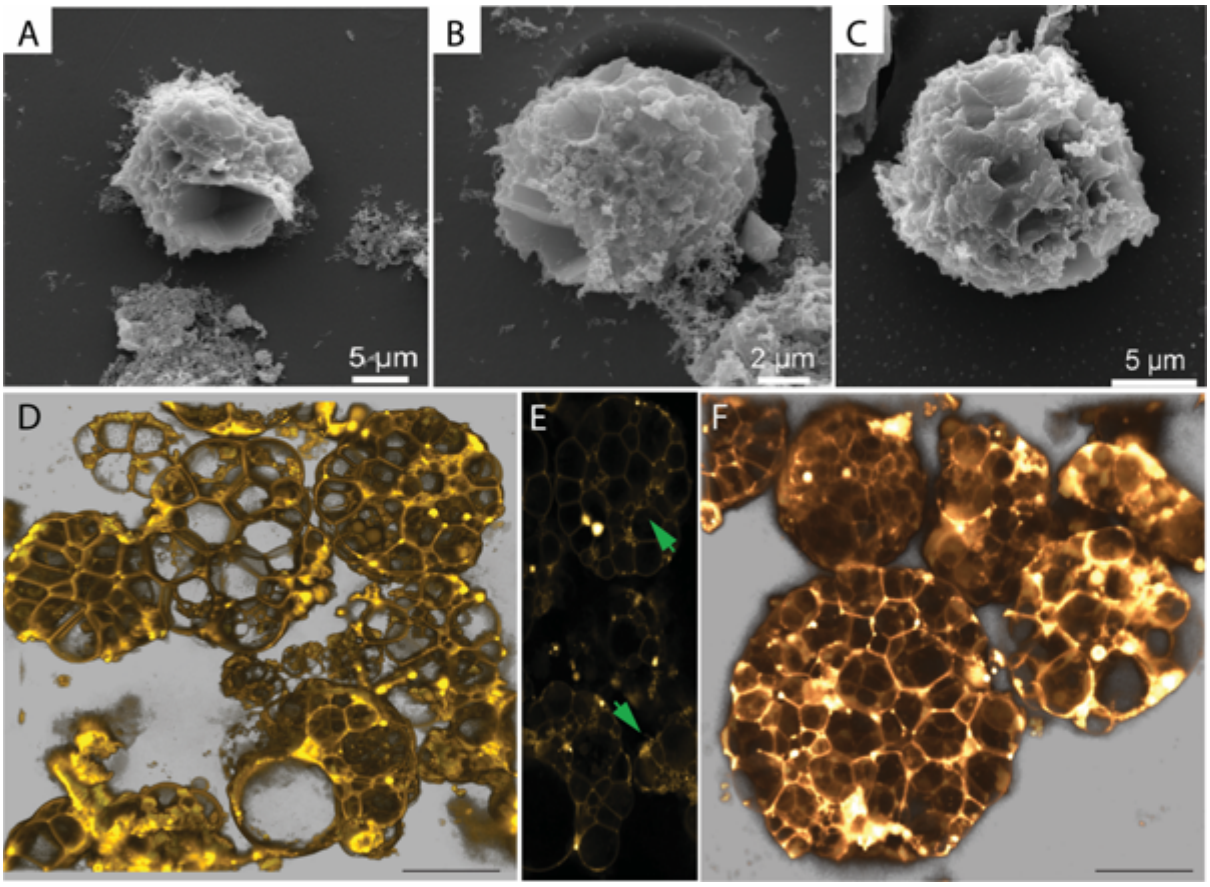


**Fig. S4. Morphological comparison between *EM-P* and the Strelley Pool Formation (SPF) microfossils.** Images A-C are SEM images of spherical microfossils reported from the SPF site (originally published by Delarue *et al*., 2019)(35). Images D & F are 3D rendered STED microscope images of similar *EM-P* cells with hexagonal internal vacuoles. Image E is the individual Z-stack image of cells shown in image D. Arrows in this image point to regions of cells with large and small polygonal vacuoles. Cells in images D-F were stained with membrane stain, FM$^{TM}$5-95. A comparison of SPF microfossils with *EM-P* was also shown in Fig. S5 & S6. Scale bar: 10µm (D & F).

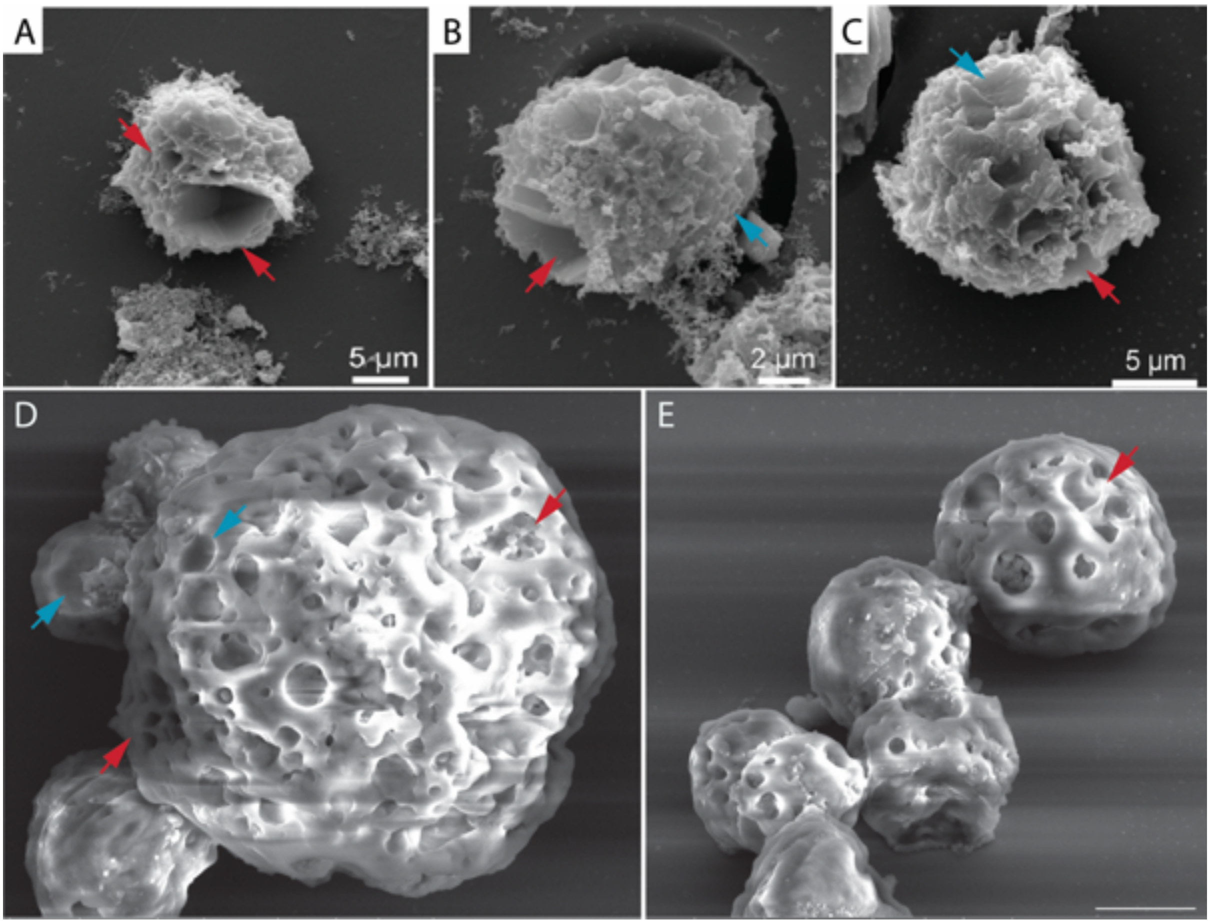


**Fig. S5. Morphological comparison between *EM-P* and SPF microfossils.**

Images A-C are SEM images of spherical microfossils reported from the SPF site (originally published by Delarue *et al*., 2019)(35). Images D & E are SEM images of morphologically analogous *EM-P* cells with hexagonal internal vacuoles. Red and cyan arrows in all images point to either surface depressions or inward cell membrane bending in both SPF microfossils and *EM-P* cells. As shown in the figure, we assume these structures are formed due to invagination, complete lysis, or individual vacuole membranes. S6D & S6E. Scale bar: 0.5µm (E).

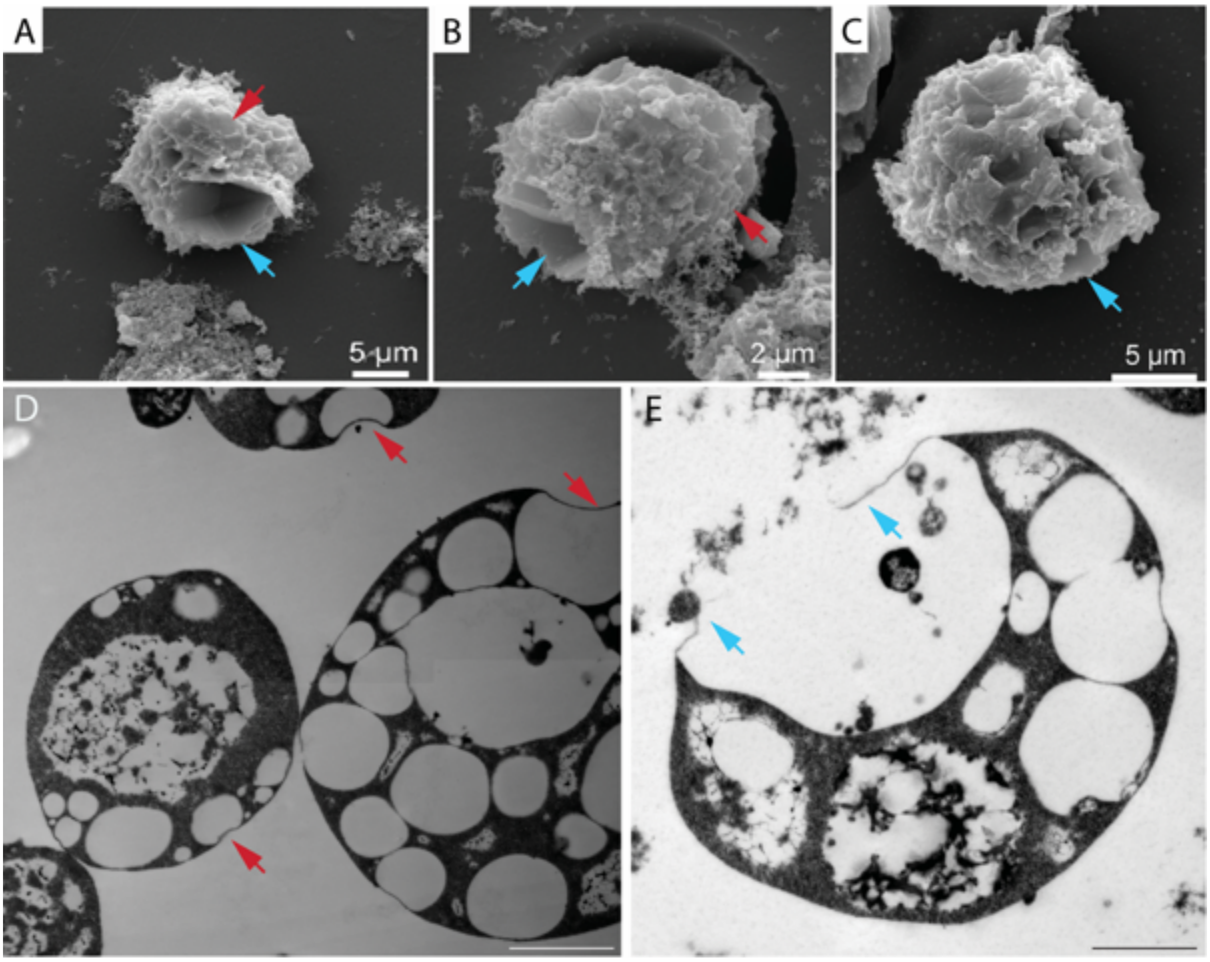


**Fig. S6. Morphological comparison between *EM-P* and SPF microfossils.**

Images A-C are SEM images of SPF spherical microfossils with spherical/polygonal surface depressions (originally published by Delarue *et al*., 2019)(35). Images D & E are TEM images of morphologically analogous *EM-P* cells. Red arrows in these images point to surface depressions formed by the partial collapse of the vacuole membrane. Blue arrows in these images point to surface depressions formed by the complete rupture of the vacuole membrane. Scale bar: 200nm (D & E).

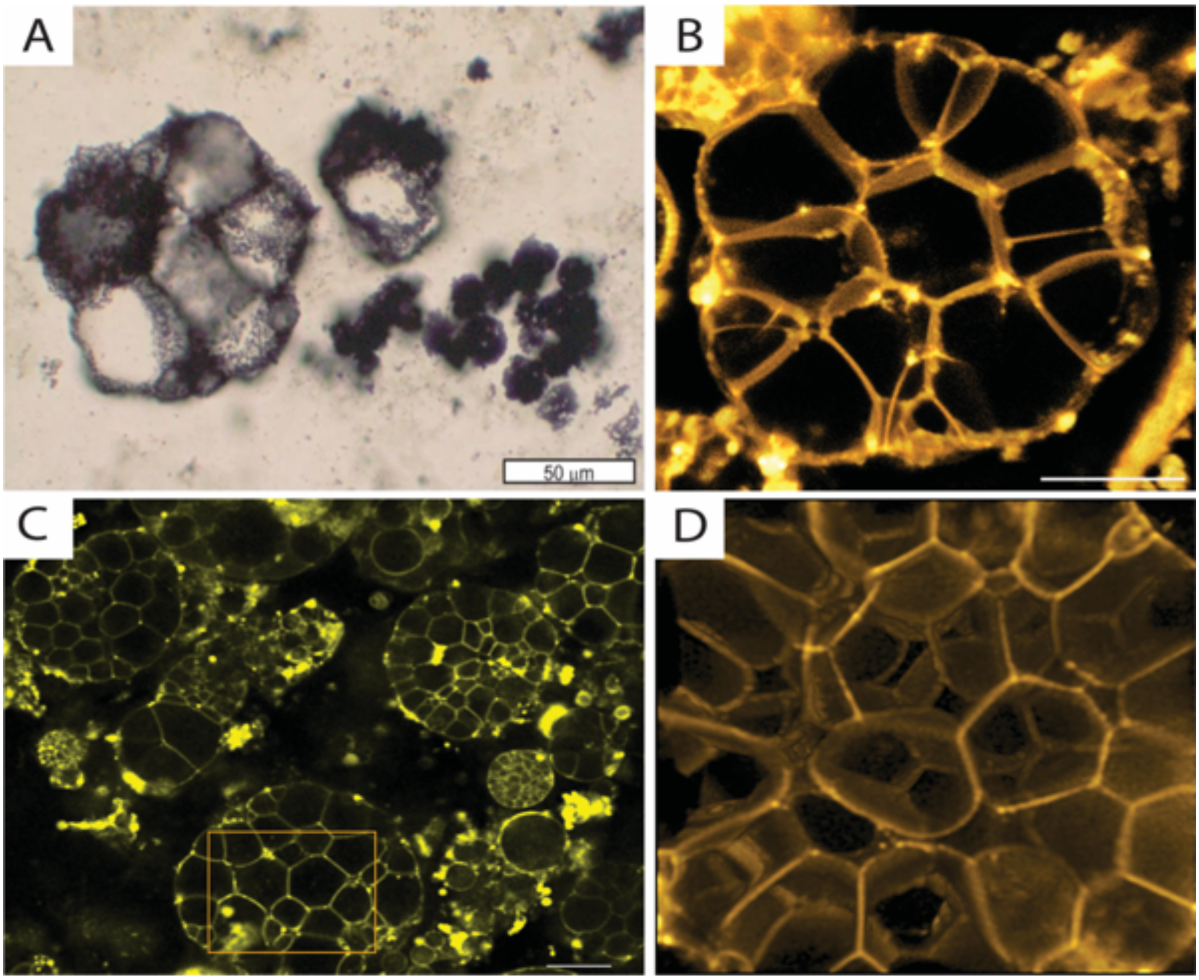


**Fig. S7. Morphological comparison between *EM-P* and the Farrel Quartzite microfossils.** Image A shows microfossils with hollow polygonal vacuoles from the Farrel Quartzite formation (originally published by Retallack *et al*., 2016)(36). Images B-D are images of morphological analogous *EM-P* cells with polygonal vacuoles. Cells in images B-D were stained with membrane stain, FM$^{TM}$5-95. Polygonal vacuoles in these images could have formed by the confinement of many vacuoles within a cell, as shown in Fig S4F. The cytoplasm in these cells was restricted to narrow spaces between the vacuoles, as shown in Fig. 1E & 1F. Scale bar: 10µm (B) and 20µm (C).

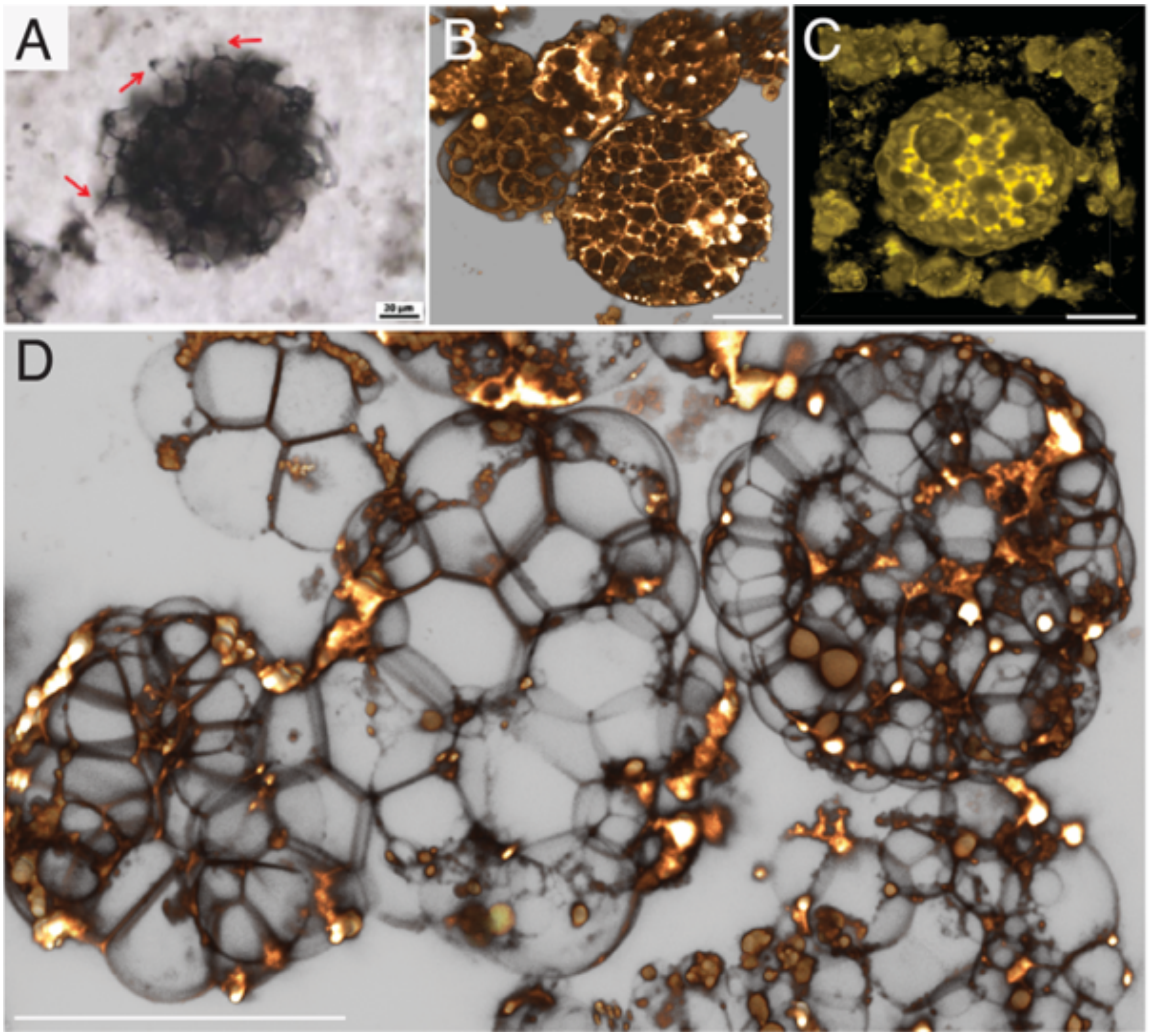


**Fig. S8. Morphological comparison between honeycomb structures reported from the Turee Creek Formations and *EM-P.*** Image A shows a spherical microfossil reported from the Turee Creek Formation (originally published by Barlow *et al*., 2018)(40). Images B-D are morphologically analogous to *EM-P* cells. Cells in these images were stained with membrane stain, FM$^{TM}$5-95. Scale bar: 20µm.

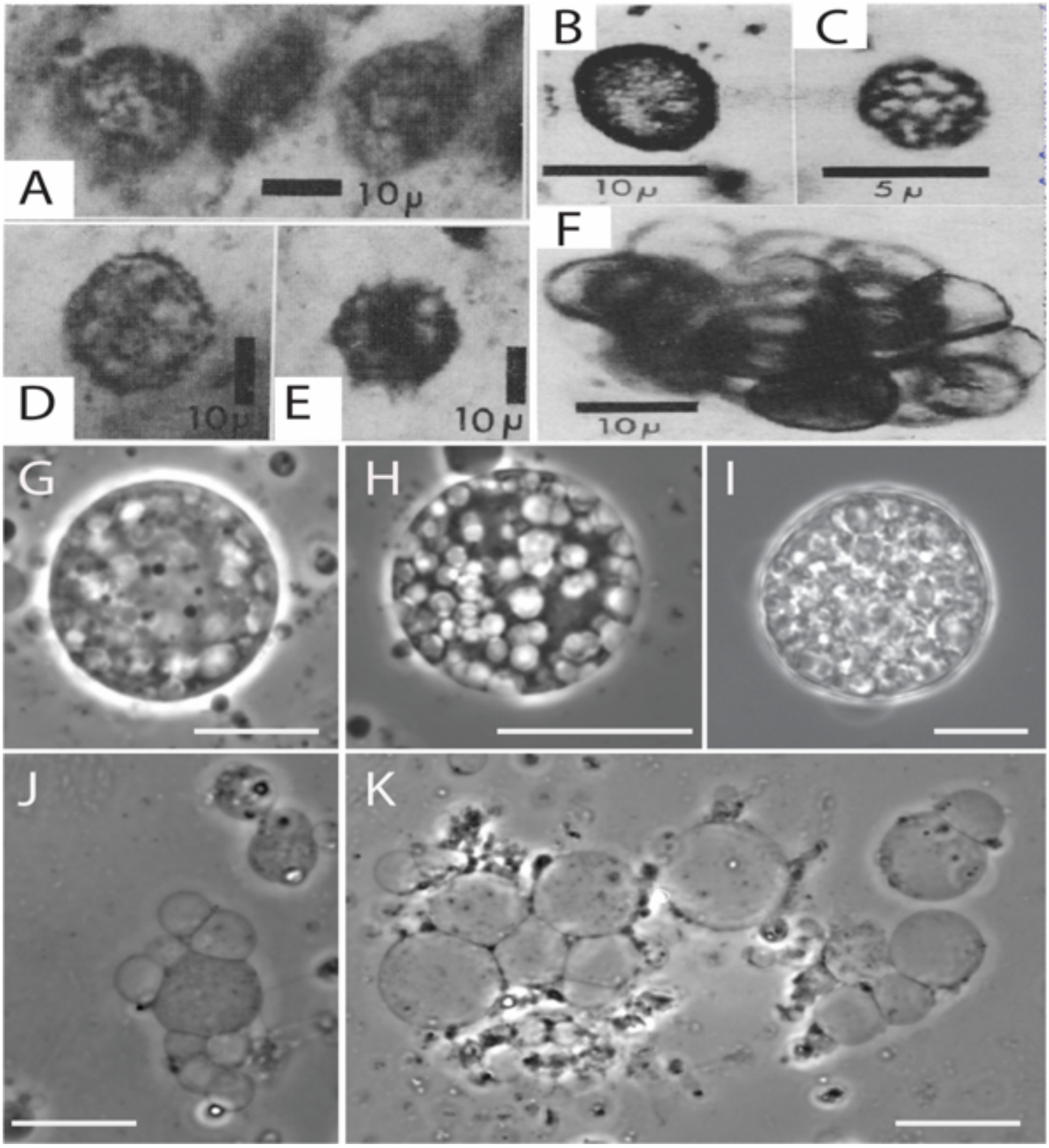


**Fig. S9. Morphological comparison between *EM-P* and the Fig Tree Formation microfossils.** Images A-F show spherical microfossils reported from Fig Tree Formation (originally published by Schopf *et al*., 1967)(38). Images G-I are morphologically analogous phase-contrast images of *EM-P*. The presence of regions with and without organic carbon can be seen in microfossil images A-E and *EM-P* cell shown in images G & H. Distinctive spherical vacuoles devoid of organic carbon can be seen in images B & C, G & H. Image F shows hollow structures morphologically similar to hollow ICVs released by the lysis of *EM-P* cells (J & K). Scale bar: 10µm (G-K).

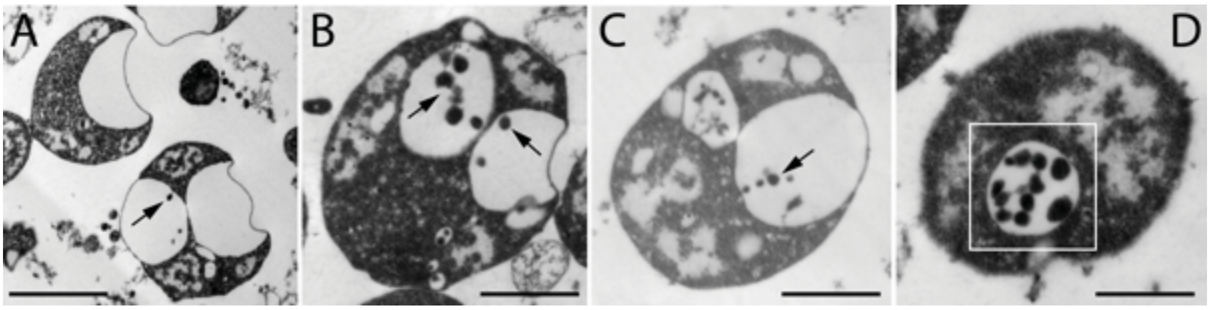


**Fig. S10. Sequential stages of intracellular daughter cell formation.** Images a-d are TEM images of *EM-P* cells showing the formation of daughter cells into hollow ICVs. Arrows in image A point to the first step in daughter cell formation by a process resembling budding. Images B-D show the gradual growth of these buds into a string of daughter cells. Scale bar: 500nm.

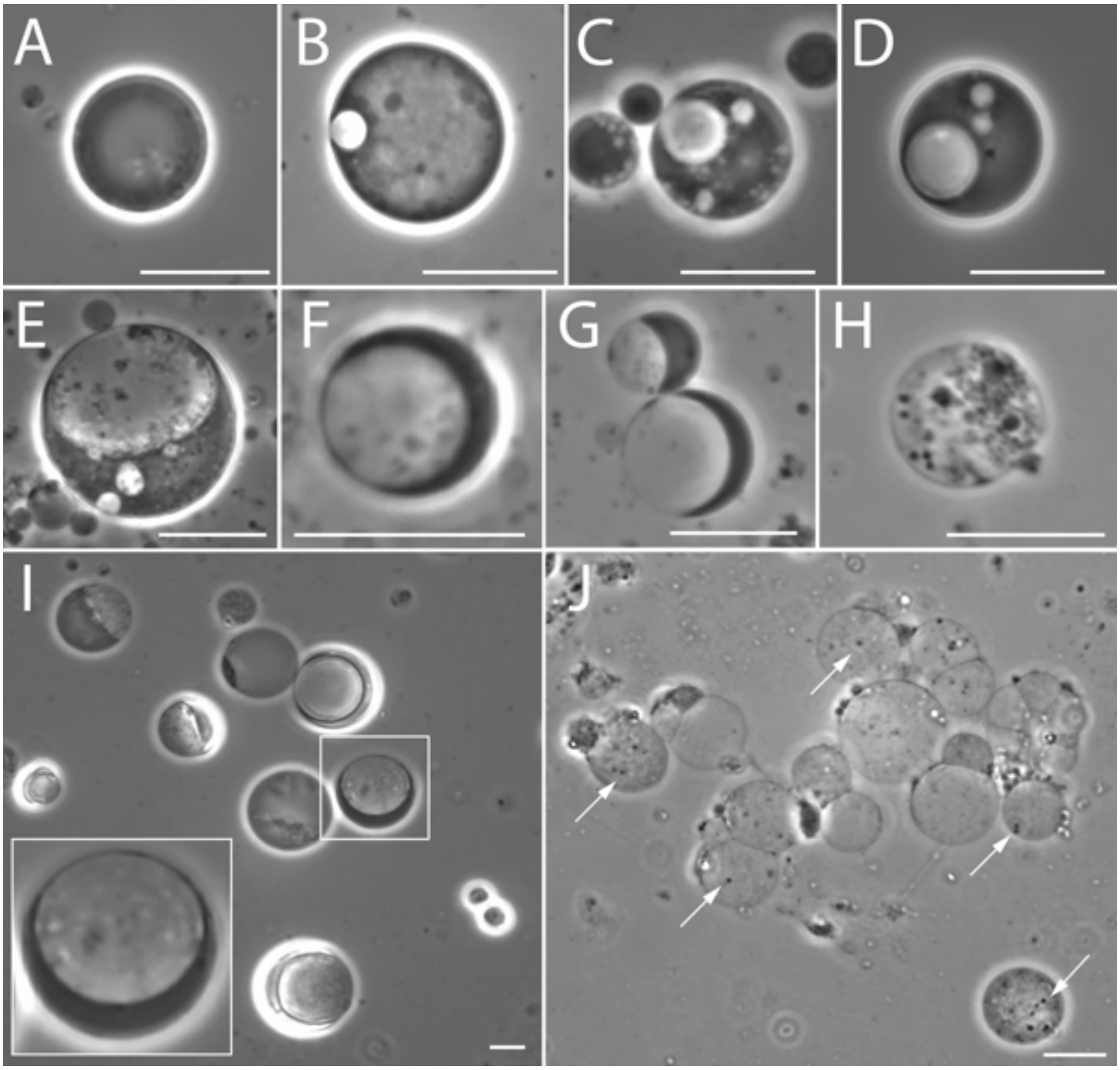


**Fig. S11. Lifecycle of *EM-P* cells reproducing via the formation of internal daughter cells.** Images A-H show phase-contrast images of *EM-P* reproducing by forming internal daughter cells. These images show a gradual increase in the volume of the ICV and a proportional decrease in the cytoplasmic volume of the cells. Images I & J show *EM-P* cells in their mid and late stationary growth phase. Insert in the image I show a magnified view of the highlighted cell. White arrows in image-J point to internal daughter cells. Scale bars: 10µm.

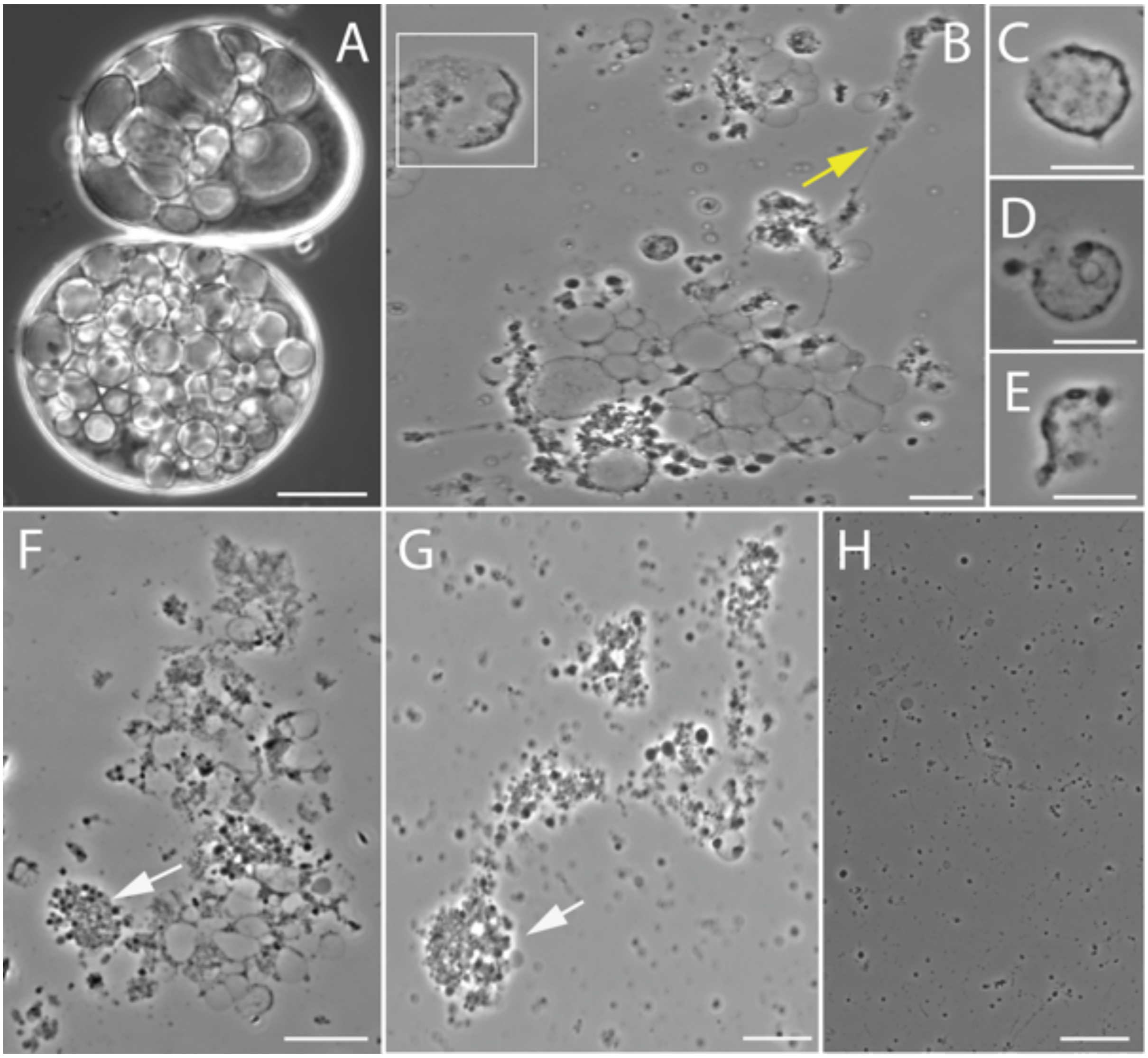


**Fig S12. Lifecycle of *EM-P* cells reproducing via the formation of internal daughter cells.** Images A-H show phase-contrast images of *EM-P* cells undergoing lysis and release of internal daughter cells. The highlighted region in image B and images C-E show a cell undergoing lysis and cells in different stages of lysis. Arrows in image B point to thin strands of membrane debris formed during cell lysis. Arrows in images F & G point to clusters of daughter cells released by the lysis of *EM-P* cells. These cell clusters gradually dispersed, leading to the formation of individual daughter cells (H). TEM images of such clusters of interconnected daughter cells and individual daughter cells with membrane overhangs are shown in Fig. S18F. Scale bar: 10µm.

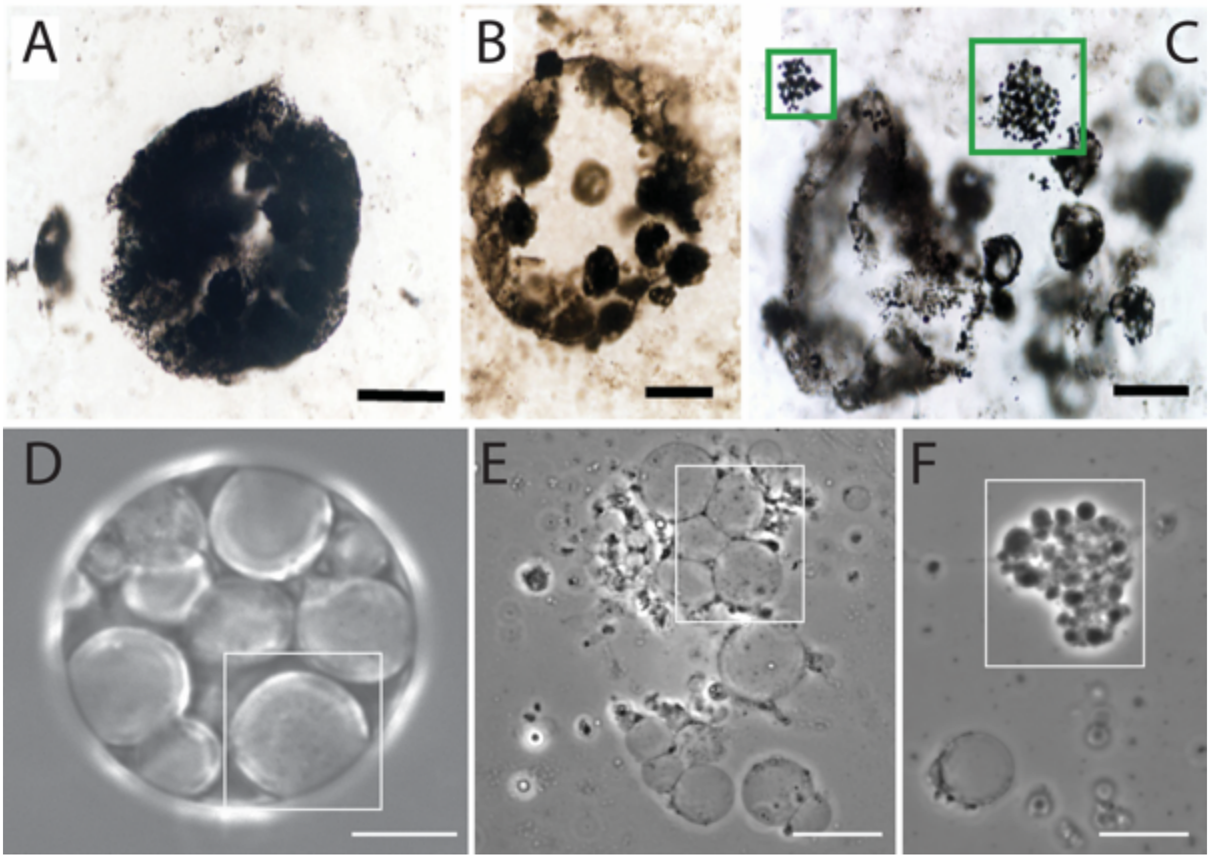


**Fig. S13. Morphological comparison between *EM-P* and the Mt. Goldsworthy Formation microfossils.** Images A-C are the microfossils reported from the Mt. Goldsworthy locality in the Pilbara formations (originally published by Sugitani *et al*., 2009)(41). Images D-F are morphologically analogous to *EM-P* cells. The highlighted regions in images D-F show in sequence the ICVs with tiny daughter cells, the lysis and release of these daughter cells into the surrounding media, the lysis of the vesicle membrane, and the release of a cluster of intracellular daughter cells. Similar lysis of cells, intracellular vesicles, and the clusters of daughter cells (highlighted region in C) can be seen in images A-C. Scale bar: 10µm.

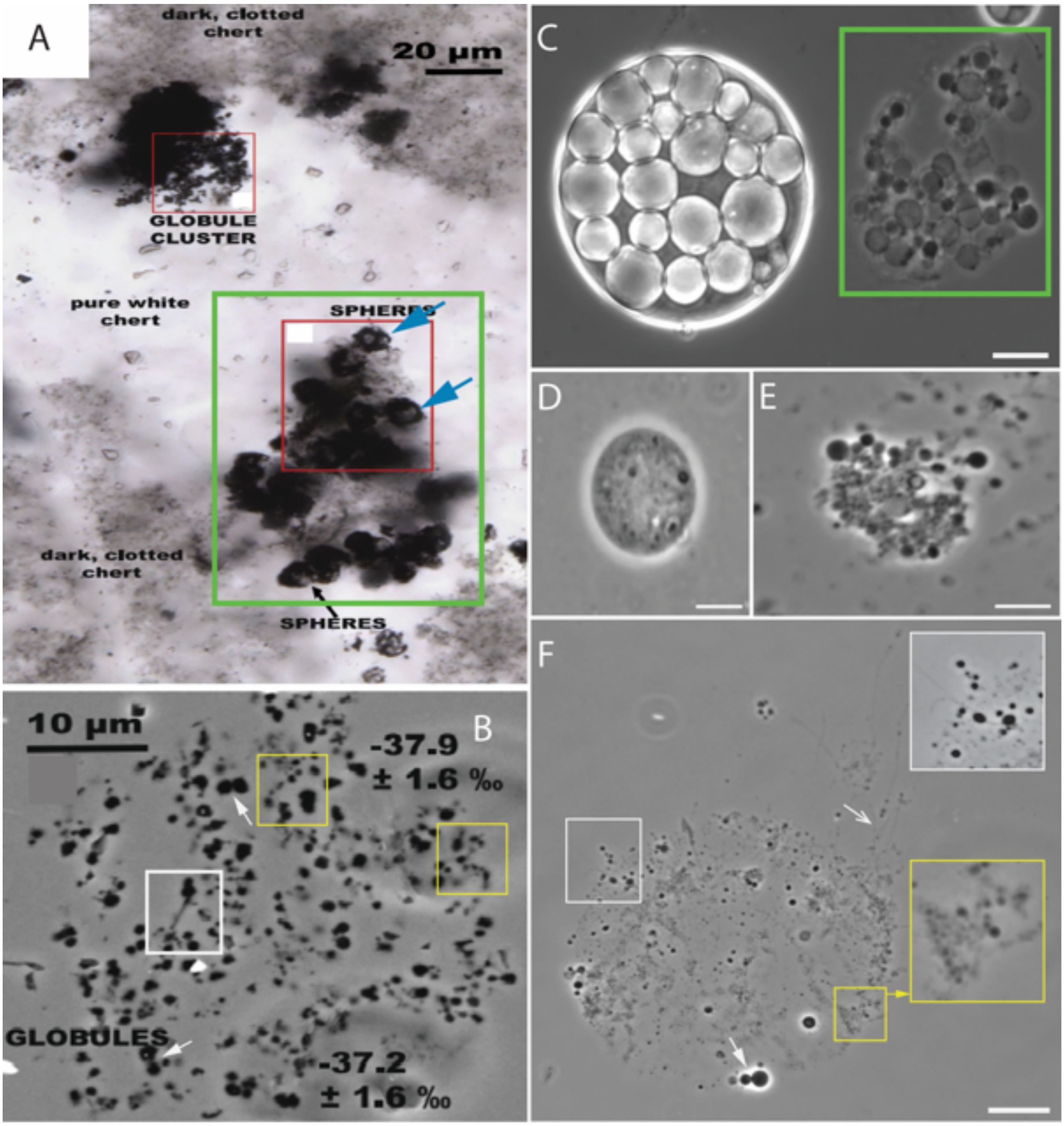


**Fig. S14. Morphological comparison between *EM-P* and the Mt. Goldsworthy Formation microfossils.** Images A & B are Mt. Goldsworthy Formation microfossils either with hollow intracellular vesicles (A, blue arrows) or spherical clumps of organic inclusions (B) (originally published by Lepot *et al*., 2013)(13). Images C-F show morphologically analogous *EM-P* cells. Image-c shows an intact and lysed *EM-P* cell. Image D shows a close-up view of the individual vesicle with daughter cells. Image E&F shows clumps of daughter cells formed after the lysis of vesicles. *EM-P* daughter cells exhibit all the morphological features of Mt. Goldsworthy microfossils, like the cluster of spherical daughter cells, daughter cells undergoing binary fission, and filamentous extensions. These cells also resemble the microfossils reported from SPF in their morphology (Fig. S19 & S20). Scale bar: 10μm (B-F).

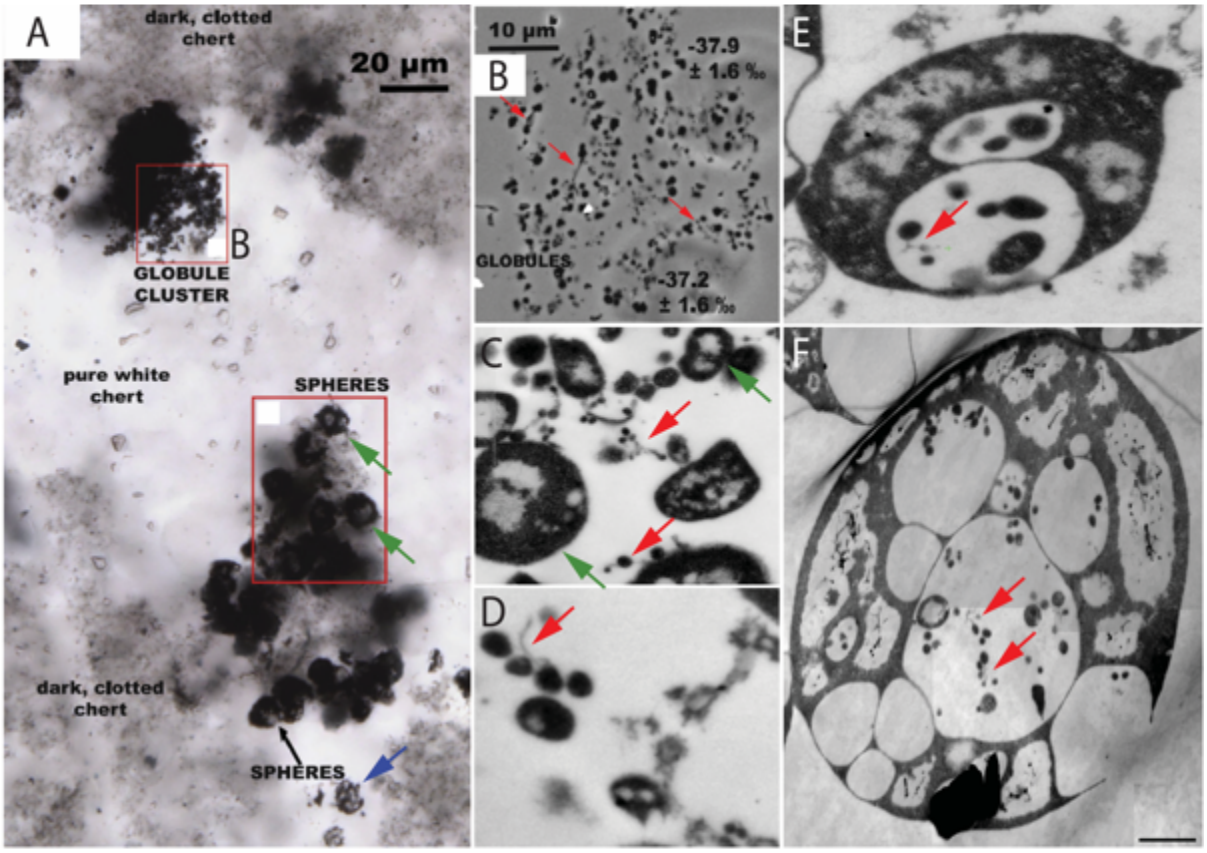


**Fig. S15. Morphological comparison between *EM-P* and the Mt. Goldsworthy Formation microfossils.** Images A & B show the microfossils reported from Mt. Goldsworthy's locality within SPF (originally published by Lepot *et al*., 2013)(13). Images C-F show the TEM images of *EM-P* cells. Red arrows in these images point to the tiny globules reported from the Mt. Goldsworthy Formation and *EM-P* daughter cells. *EM-P* daughter cells and the globules reported from the Mt. Goldsworthy Formations exhibit similar morphological features like membrane overhangs. The green arrows in images A & C point to the *EM-P* cells with uneven distribution of intracellular organic carbon. The blue arrow in image A points to a spherical cell with hollow intracellular spaces. Similar *EM-P* cells were shown in Fig. 1 & S7-S9. Scale bar: 500 nm (C-E) and 1μm (F).

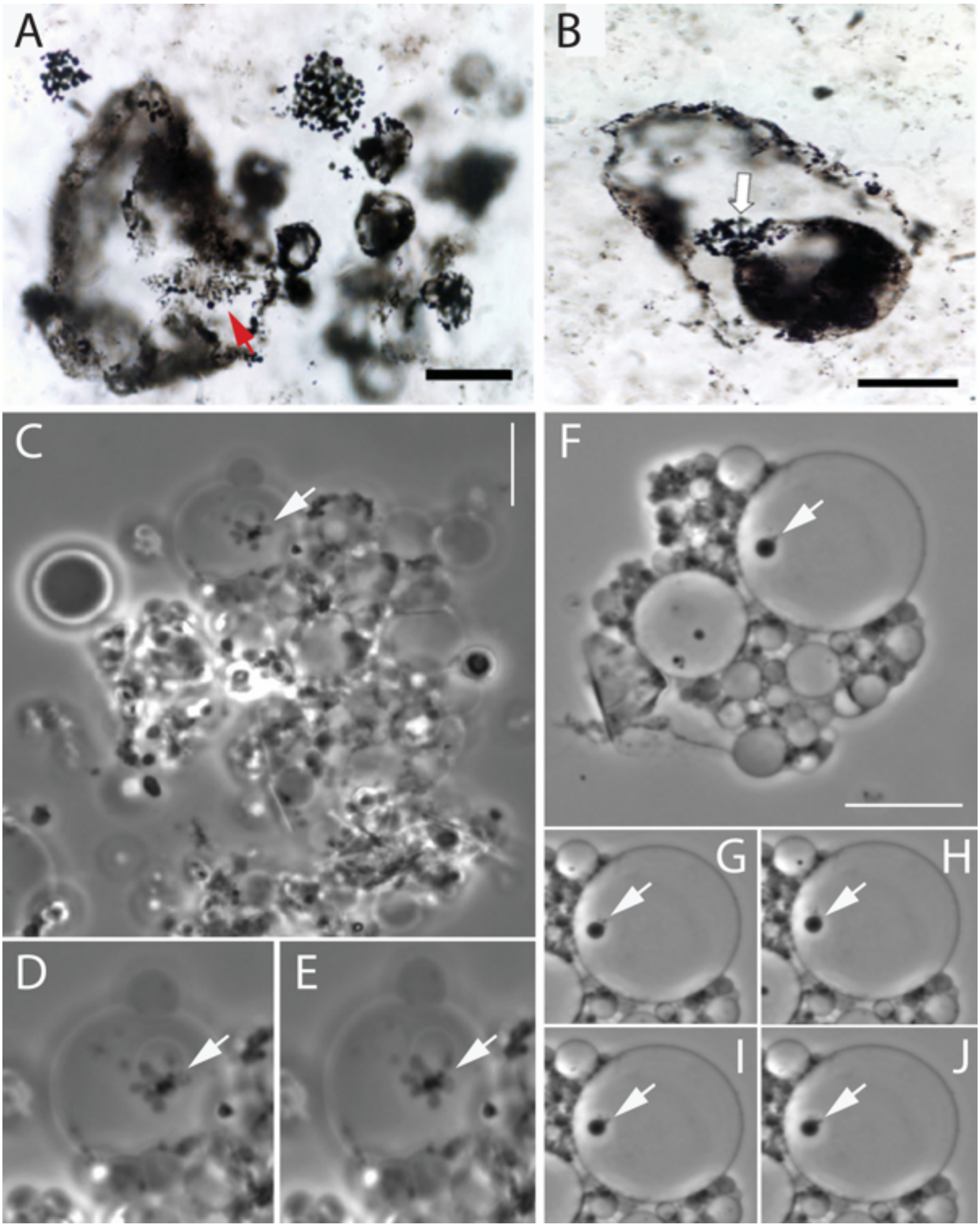


**Fig. S16. Morphological comparison between *EM-P* and the Mt. Goldsworthy Formation microfossils.** Images A & B are the microfossils reported from the Mt. Goldsworthy Formation (Sigutani et al., 2007)(37). These images show hollow spherical cells associated with clusters of spherical structures. Images E-J are the phase-contrast microscope images of *EM-P*, morphologically similar to the Mt. Goldsworthy Formation microfossils. Images D & E are the time series images of the magnified region of image C. Images G-J are the time series images of the magnified region of image F. Time series images show the movement of spherical structures attached to the inner spheroid. Scale bars: 20µm (A & B) and 10µm (C & F).

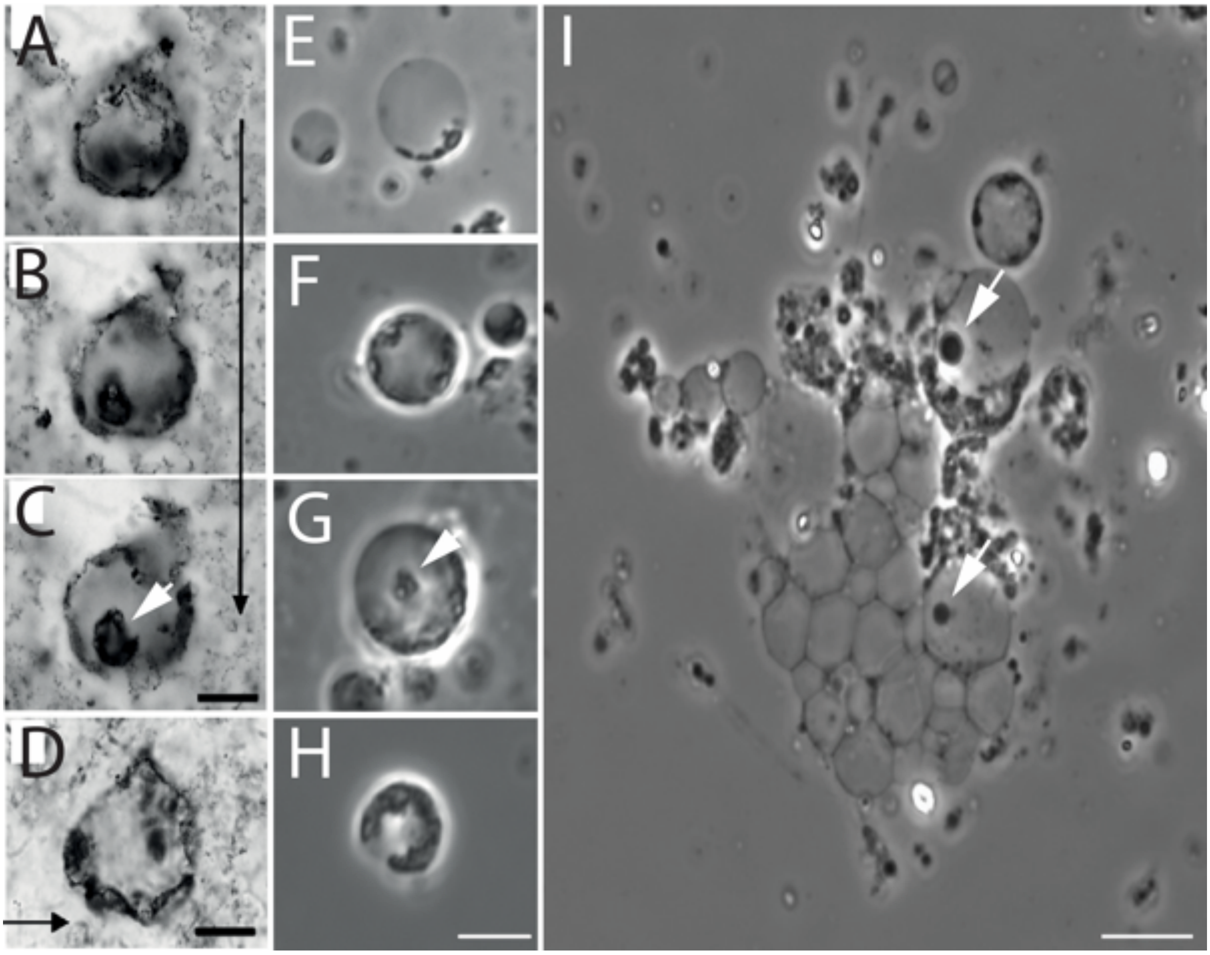


**Fig. S17. Morphological comparison between *EM-P* and the Mt. Goldsworthy Formation microfossils.** Images A-D are the microfossils reported from the Mt. Goldsworthy Formation (Sigutani et al., 2007)(37). The black arrow in these images points to the deepening focal depths. The white arrow in image C points to the internal carbon-rich spheroidal structure. Images E-I are the phase-contrast microscope images of *EM-P*, morphologically similar to Mt. Goldsworthy formation microfossils. Arrows in these images point to a similar internal spheroidal structure observed in the Mt. Goldsworthy Formation microfossils. Scale bar: 20µm (A-D) and 10µm (E-I).

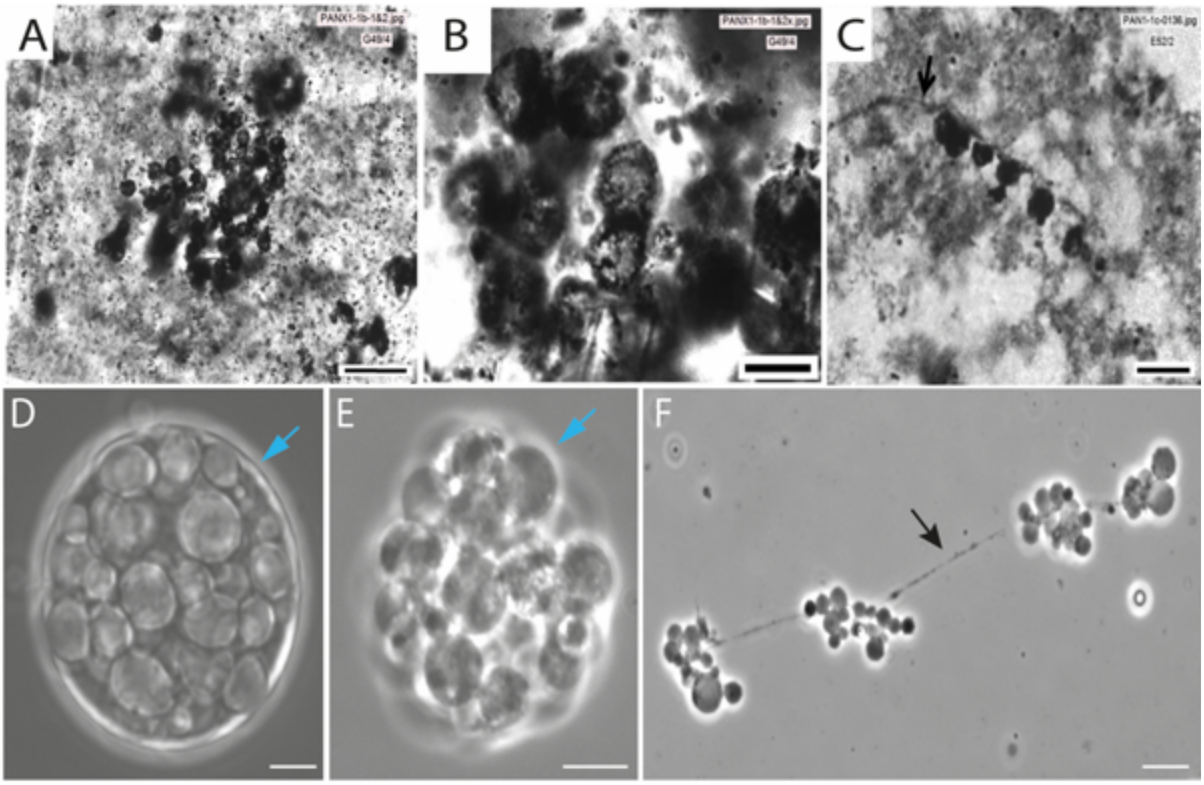


**Fig. S18. Morphological comparison between the SPF microfossils and *EM-P*.**
Images A-D are hollow spherical SPF microfossils with internal inclusions (originally published by Sugitani *et al*., 2013)(39). Images D-F are images of *EM-P* cells undergoing lysis (E) and dispersion (F) of intracellular vacuoles. Cyan arrows in images D and E point to the presence and absence of the outer cell membrane. Black arrows in images C & F point to filamentous structures interlinking spherical structures. Scale bar: 50µm (A), 20µm (B & C), 2µm (D & E), and 5µm (F).

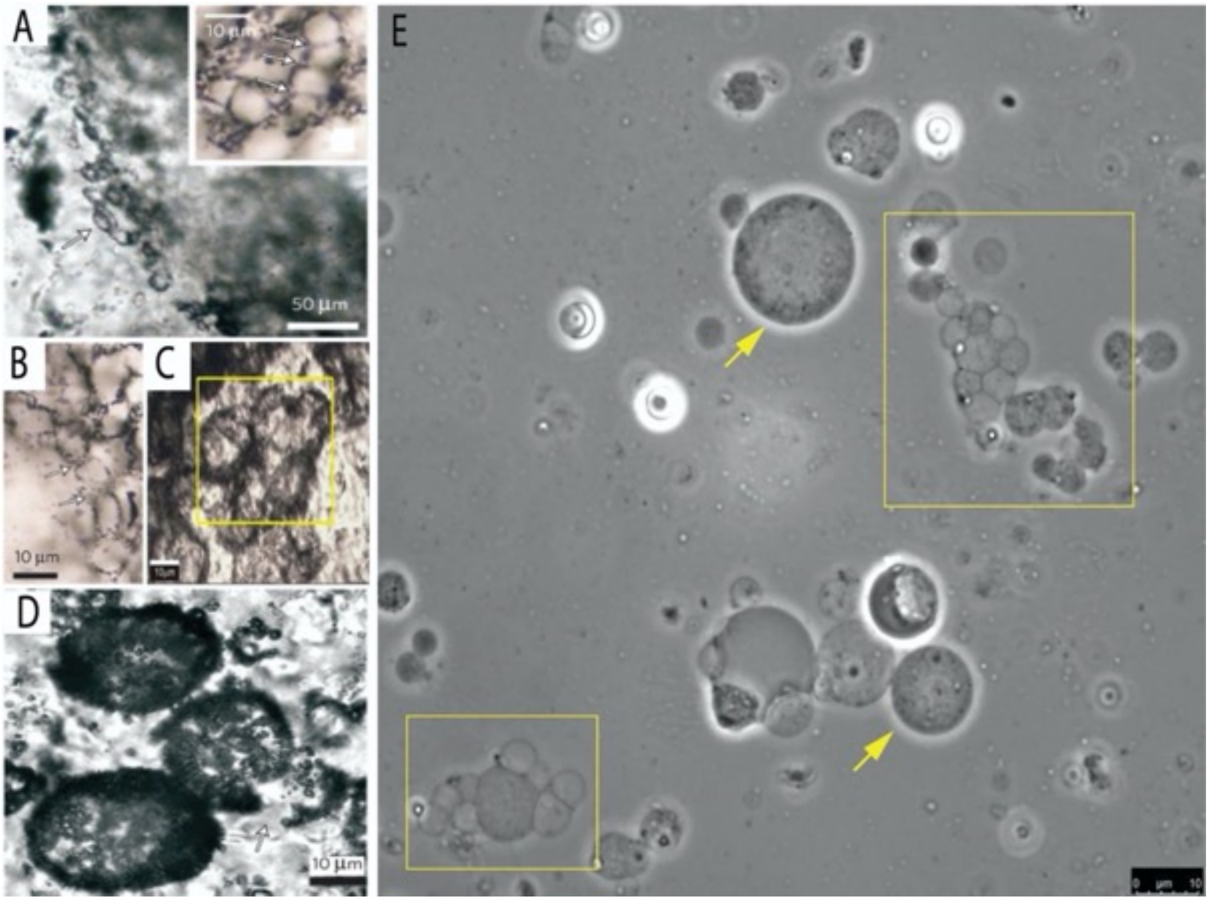


**Fig. S19. Morphological comparison between *EM-P* and SPF microfossils.** Images A-D depict SPF microfossils with intracellular organic inclusions (originally published by Wacey *et al*., 2011). Image-E shows morphologically analogous *EM-P* cells. Images A-C show chains and clumps of cells. A similar cluster of cells can be seen in image E (boxed regions). Image D shows spherical microfossils with intracellular spherical inclusions. Similar *EM-P* cells with intracellular daughter cells can be seen in image E (yellow arrows). Lysis and release of daughter cells can be seen in Fig. S12. Also see Movie 4.

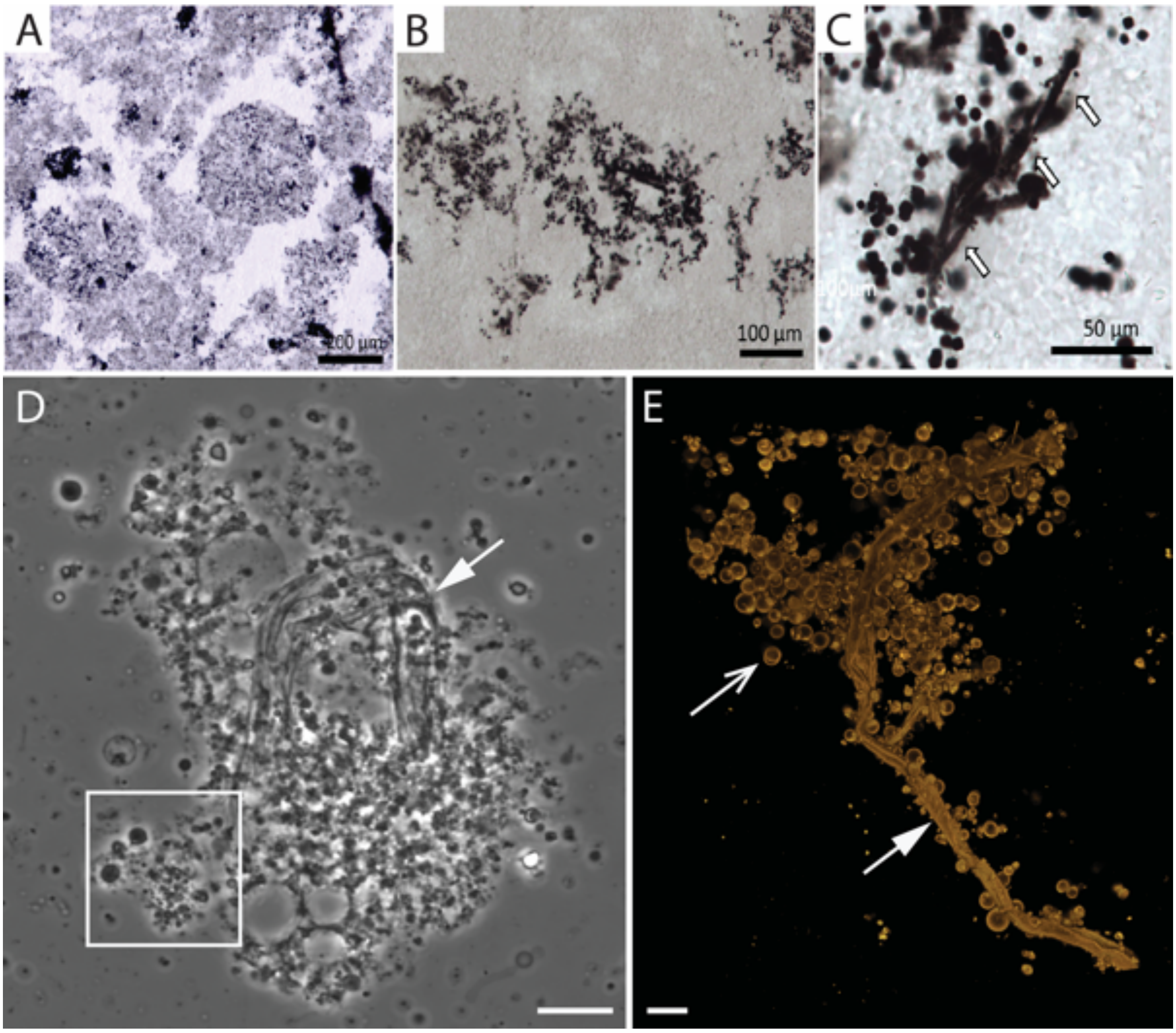


**Fig. S20. Morphological comparison between *EM-P* and the SPF microfossils.**
Images A-C show clusters of faint and dark spherical microfossils reported from SPF (originally published by Sugitani *et al*., 2015)(42). Images D & E show morphologically similar structures observed in *EM-P* incubations. Image A shows spherical clusters of spherical granules. Based on their morphological similarity, they likely represent the tiny intracellular daughter cells, as shown in D (highlighted region) and Fig. S19. The thread-like structures in c (arrows) likely are the membrane debris often found associated with the release of daughter cells, as shown in D & E (arrows). Scale bars in images D & E are 10µm.

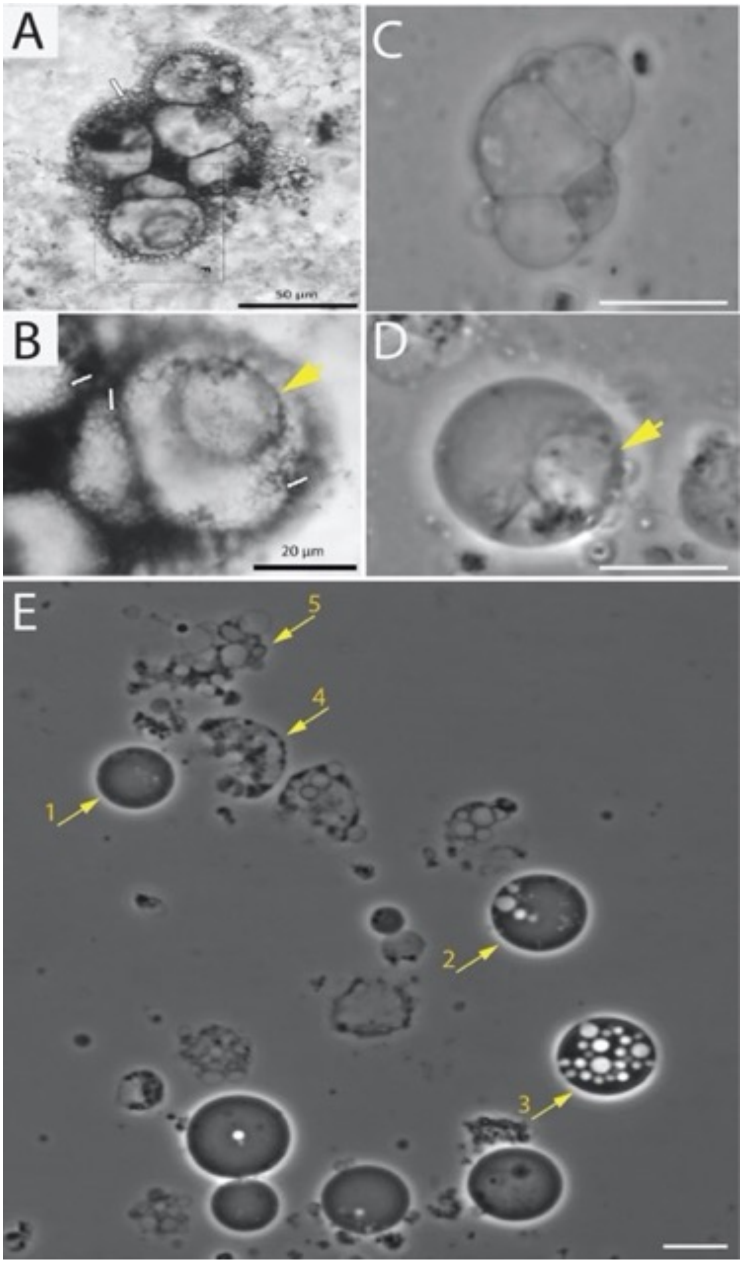


**Fig. S21. Morphological comparison between *EM-P* and the Waterfall microfossils**
Images A & B are microfossils reported from the Waterfall locality within the SPF (originally published by Sugitani *et al*., 2015)(42). Images C & D show morphologically analogous *EM-P* cells. Arrows in images C & D point to spherical intracellular vacuoles within the cell. Image E shows sequential stages involved in the formation of structures shown in images A & B. Sequential steps are indicated by numbers next to the arrows. Scale bars: 10µm (C-E).

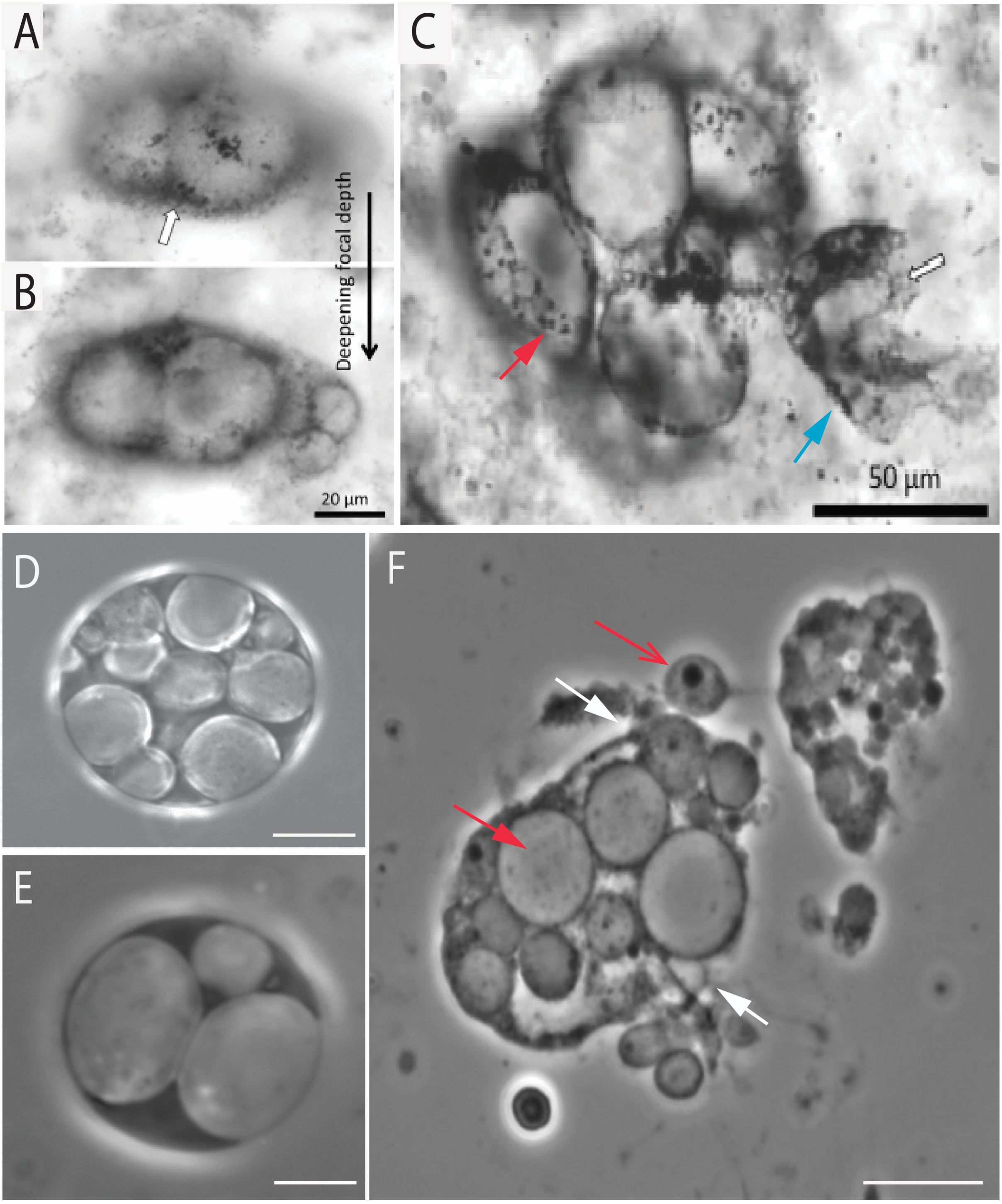


**Fig. S22. Morphological comparison between *EM-P* and Waterfall microfossils.** Images A, B, and C are microfossils reported from the waterfall locality within SPF (originally published by Sugitani *et al*., 2015)(42). Images D-F are phase-contrast images of morphologically analogous structures formed by *EM-P*. Red arrows in these images point to intracellular vacuoles with daughter cells of different sizes. Open red arrows point to a vacuole with a relatively large daughter cell. The white arrows in image F points to the ruptured region of the cell membrane and release of the ICVs. Scale bars: 10µm (D-F).

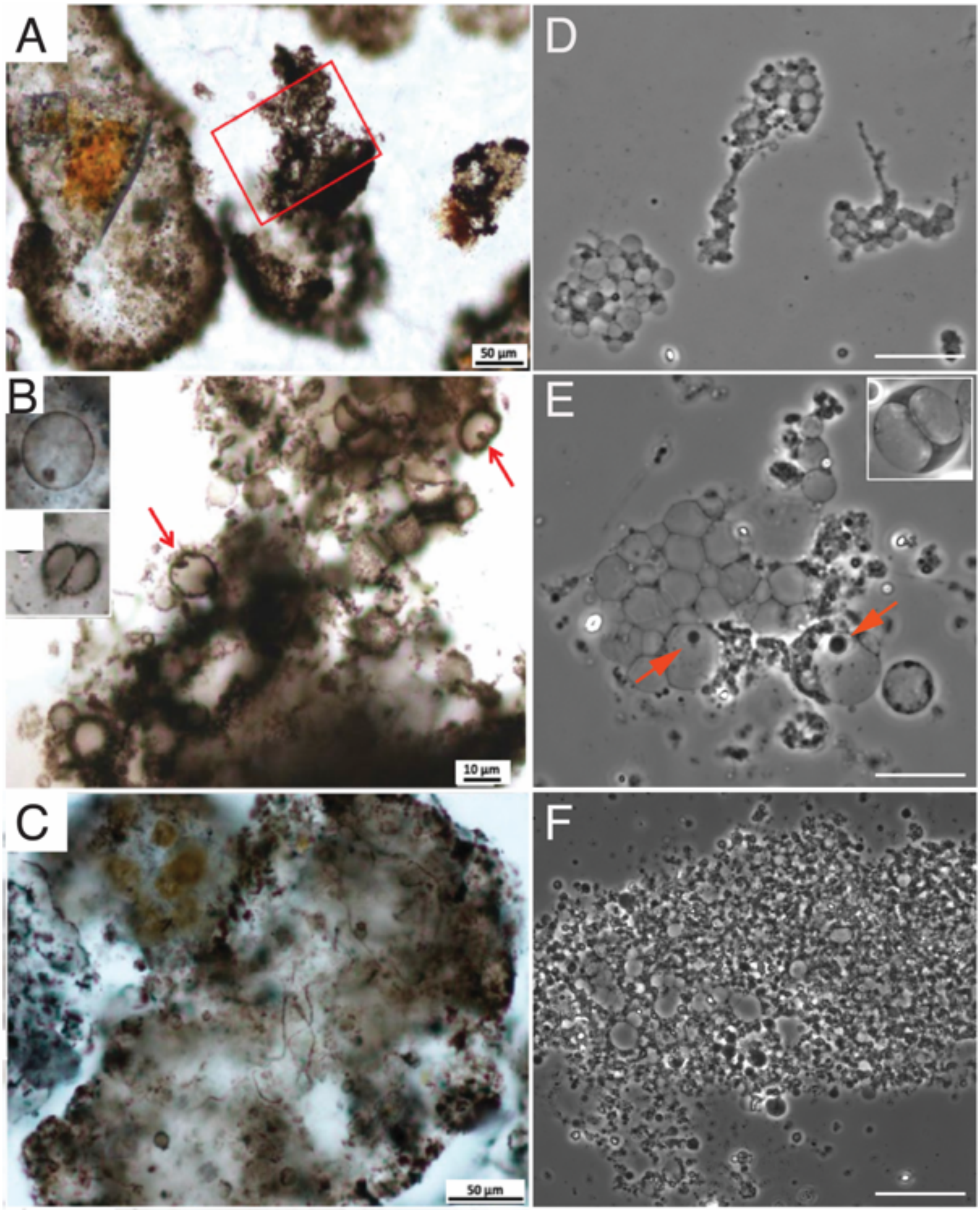


**Fig. S23. Morphological comparison between honeycomb structures reported from the Turee Creek Formations and *EM-P.*** Images A, B, and C are microfossils reported from the Turee Creek locality within the SPF (originally published by Barlow *et al*., 2018)(40). Images D-F are phase-contrast images of morphologically analogous structures formed by *EM-P*. The boxed region in image A resembles the lysed cells of *EM-P*, shown in image D. Arrows in image B & E point to the similar organic carbon inclusions (B) and morphologically analogous daughter cells (E) within a hollow spherical *EM-P* cell. Insert in images B & E show spherical cells with two intracellular vesicles or nearly equal volume. Scale bars: 10µm (D-F).

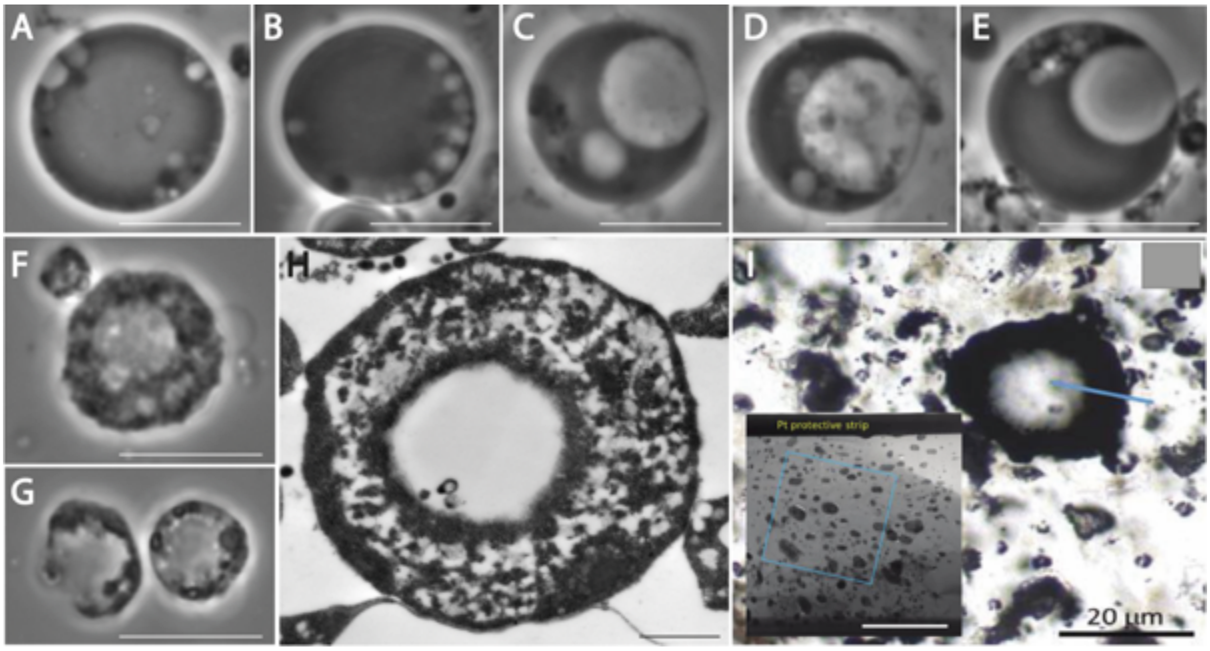


**Fig. S24. Sequence of morphological transformations leading to the formation of *EM-P* cell having an appearance of a thick cell wall.** Images A-G show the sequential morphological transformations of *EM-P* cells with one large central vacuole and multiple smaller vacuoles. Over the course of its growth, smaller vacuoles were squeezed between the cell membrane and the vacuole membrane (F & H), leading to the appearance of a porous cell wall. Images F-G and H are phase-contrast and TEM images of such *EM-P* cells. The image-I shows a spherical microfossil reported from Dresser formation (originally published by Wacey *et al.,* 2018)(10). Insert in the image-I shows the TEM image of the region indicated by the blue line. The porous nature of this region is similar to the *EM-P* cell shown in image H. Scale bars: 10µm (A-G) and 200nm (H).

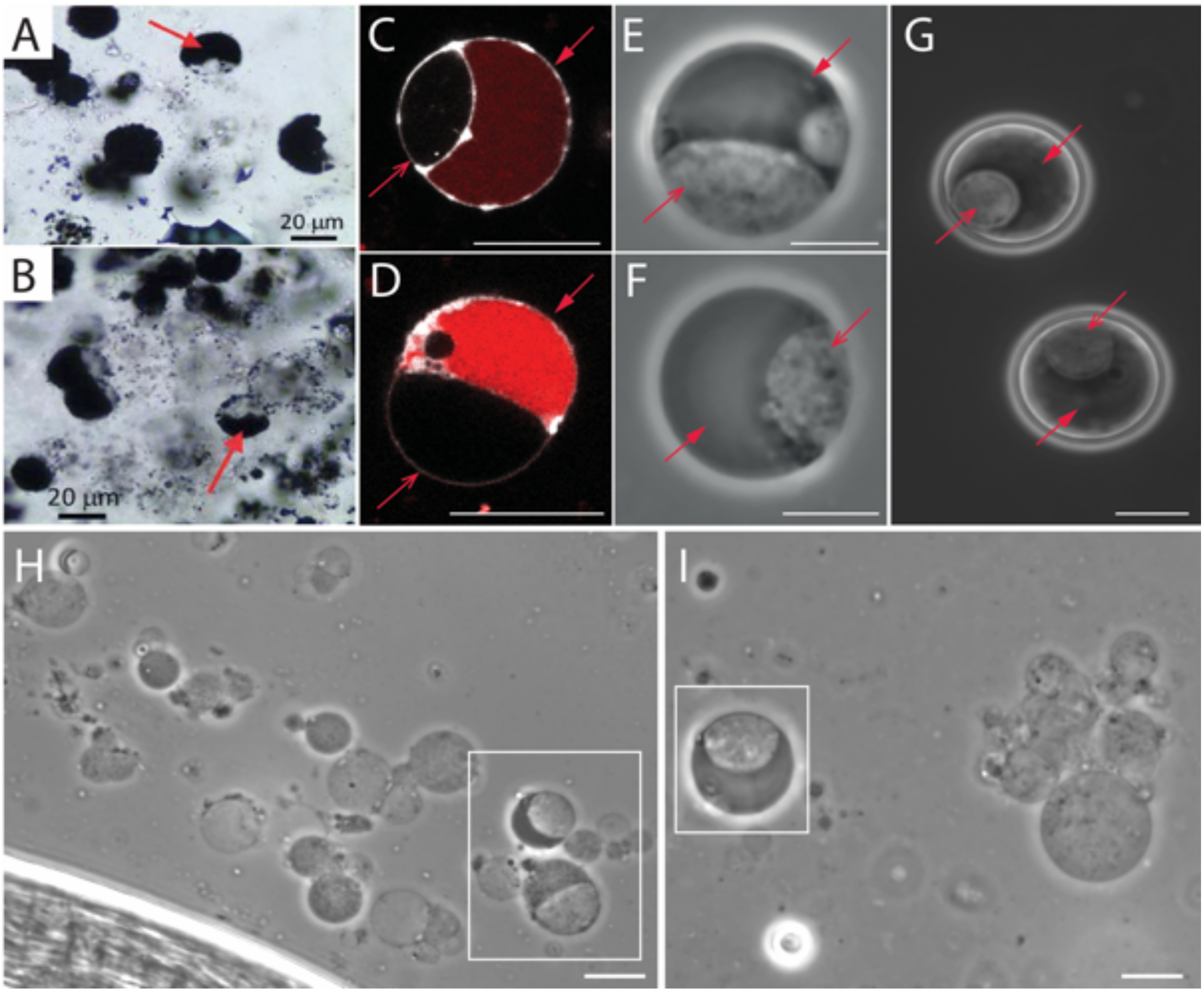


**Fig. S25. Morphological comparison of organic structures reported from the Dresser Formation with *EM-P.*** Images A & B are spherical microfossils reported from the Dresser formation (originally published by Wacey *et al.,* 2018)(10). Images C & D are STED microscope images of morphologically analogous *EM-P* cells. Cells in these images were stained with FM™5-95 (membrane, white) and PicoGreen™ (DNA, red). Images G-I are phase-contrast images of morphologically analogous *EM-P* cells. Closed arrows in these images point to the regions of the cell with the cytoplasm (organic carbon), and open arrows point to the hollow space created by the ICVs. Scale bars 10µm (C-I).

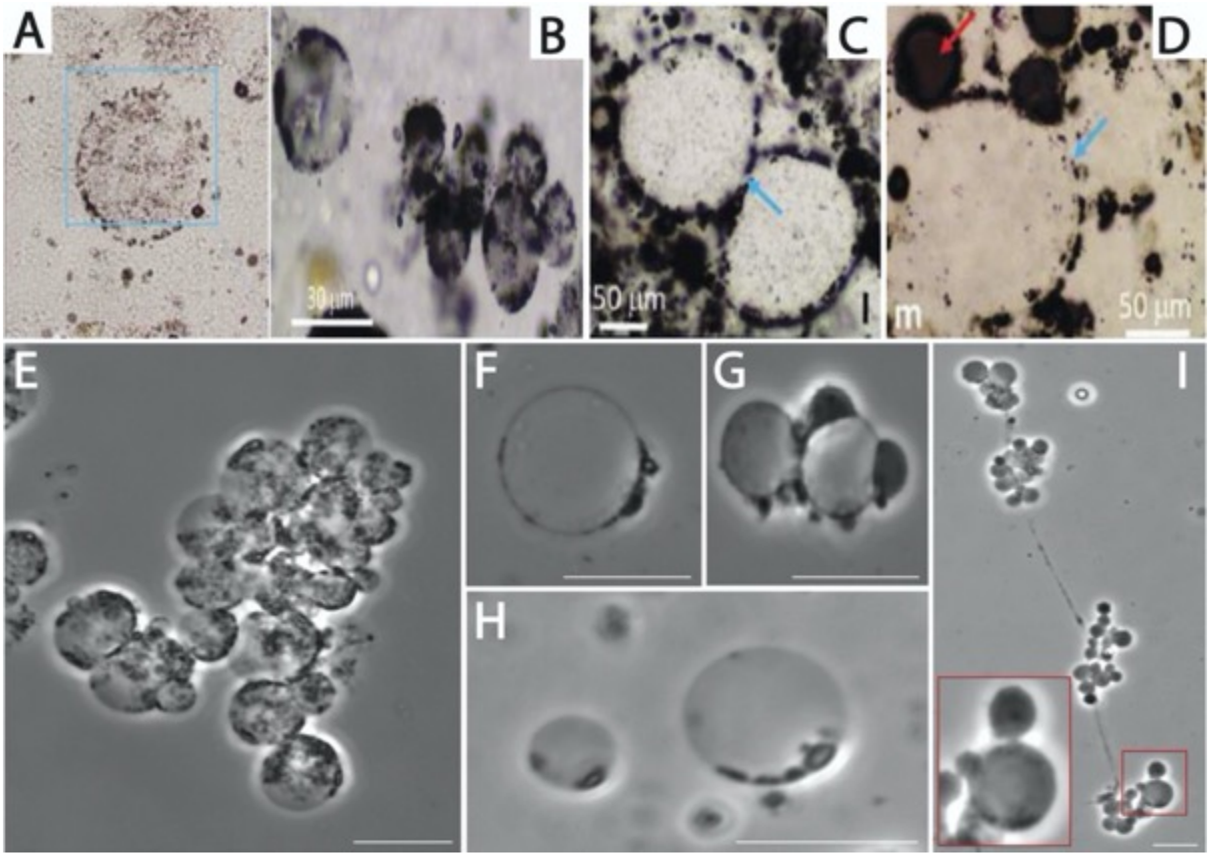


**Fig. S26. Morphological comparison of Dresser formation spherical microfossils with *EM-P.*** Images A-D were aggregations of spherical microfossils reported from Dresser formation (originally published by Wacey *et al.,* 2018)(10). E-I are images of morphologically analogous structures observed from *EM-P*. Blue arrows in images C & D point to cells with discontinuous cell walls. Similar cell boundary features can be seen in *EM-P* (E-H). Sequential stages involved in forming such *EM-P* cells are shown in Fig. S27. The red arrow in image D points to structures with organic carbon attached to the hollow spherical cells. Image-g showed morphologically analogous *EM-P* cells. Scale bar: 10µm (E-I).

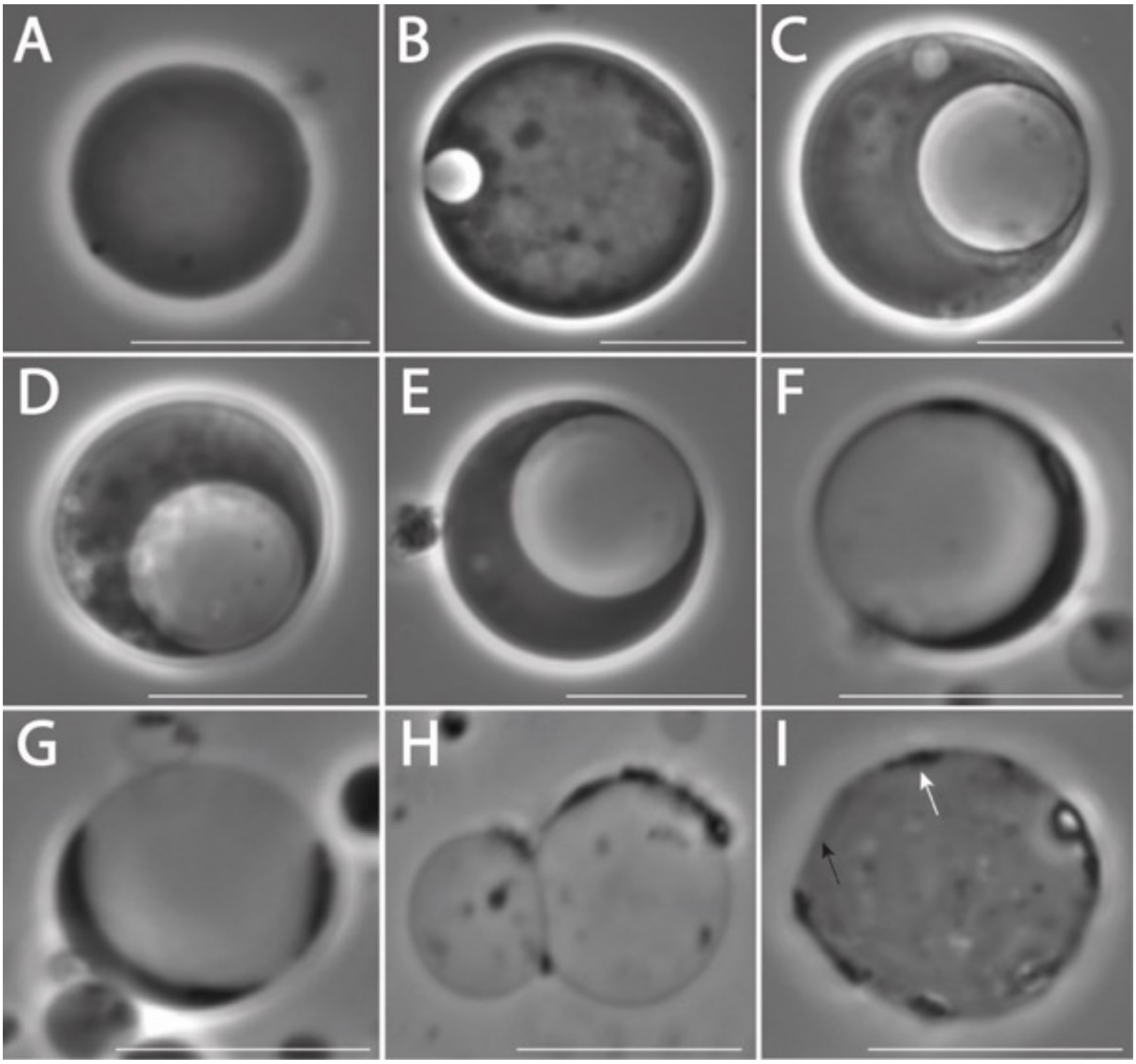


**Fig. S27. Sequential morphological transformation of *EM-P* cells with cytoplasm to hollow vacuoles.** Images A-I phase-contrast images of *EM-P*. In sequence, they show formation (B) and an increase in the size of the intracellular vacuole (B-G). Over the course of its growth, cytoplasm from the parent cell is transferred into the daughter cells in the vacuole (barely visible tiny spherical structures in the vacuole), which leads to the gradual depletion of cytoplasmic volume and a gradual increase in vesicle volume. In their late growth stages, the presence of cytoplasm is restricted to the periphery of the cell as discontinuous patches (H & I). Scale bars: 10µm.

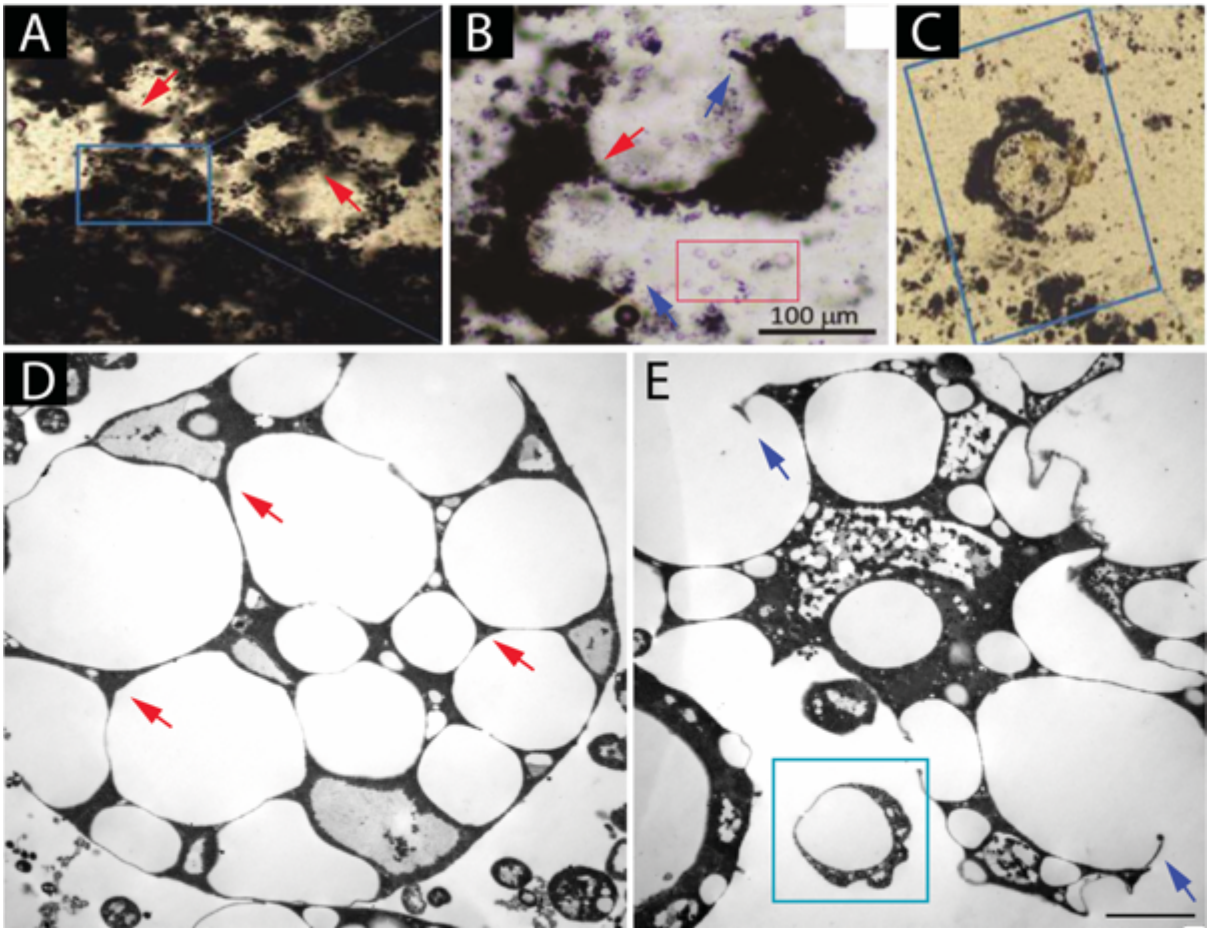


**Fig. S28. Morphological comparison of the Dresser Formation spherical microfossils with *EM-P*.** Images A-C were aggregations of spherical microfossils reported from the Dresser Formation (originally published by Wacey *et al.,* 2018)(10). Images D &E are TEM images of morphologically analogous *EM-P* cells. Image A shows hollow spherical aggregations that were devoid of organic carbon. Similar *EM-P* cells were shown in image D. Red arrows in this image point to Y-shaped junctions with organic carbon. Such *EM-P* cells, over the course of their growth, underwent membrane rupture to release daughter cells (image E, purple arrows). The process of lysis and dispersion of these cells is shown in Image E. The morphology of these *EM-P* cells was very similar to the organic structures reported from the Dresser Formation (Image B). Like *EM-P* cells, daughter cells can also be seen nearby (red box). Image C shows an isolated spherical structure, with internal spherical inclusions with the presumed thick and uneven cell wall. Morphologically analogous *EM-P* cells can be seen in Image E (Cyan box). Scale bar: 250nm (E).

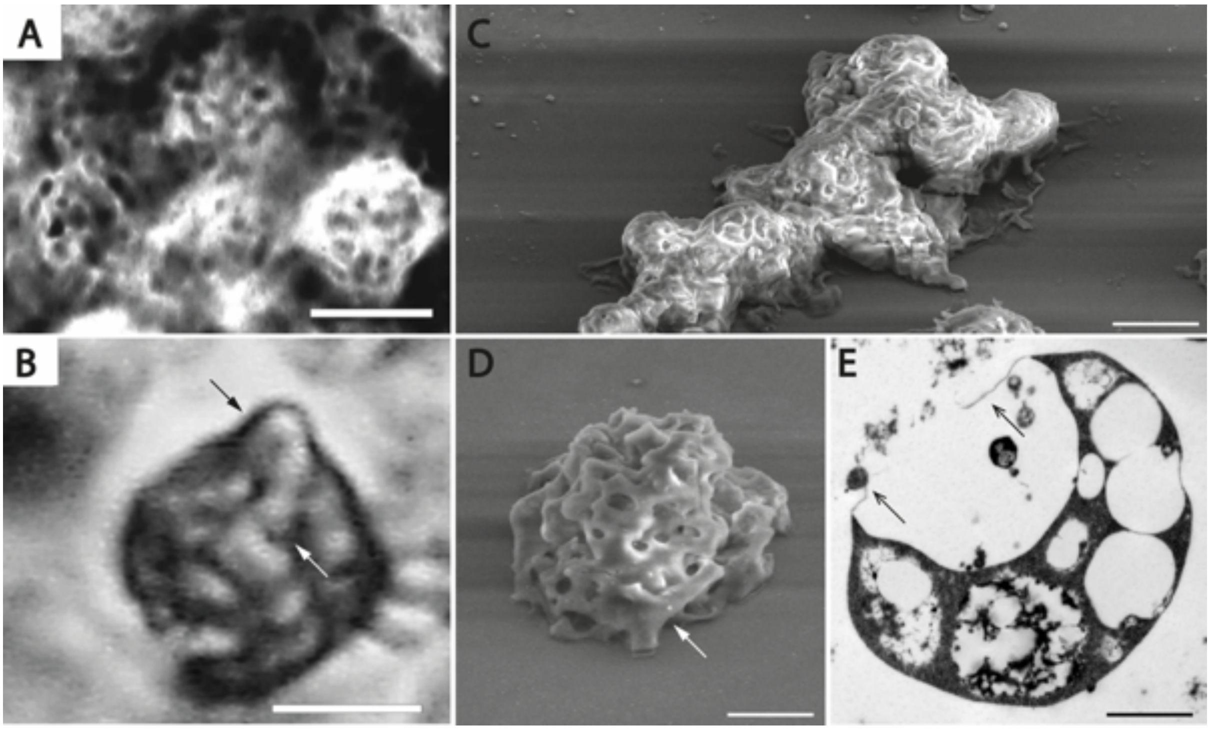


**Fig. S29. Morphological comparison between *EM-P* and the Kromberg Formation microfossils.** Images A & B are microfossils reported from the Kromberg Formation (originally published by Kaźmierczak *et al.,* 2019)(44). Images C, D & E are SEM and TEM images of morphologically analogous *EM-P* cells. Such *EM-P* cells were formed by deflation caused by lysis and release of daughter cells. SEM images show similar surface depressions in both the Kromberg microfossils and *EM-P* cells (arrows in A & C). Different stages of surface depression formation are shown in Fig. S1G-K. Arrows in images B & D point to similar surface textures of microfossils and *EM-P* cells. Scale bars: 10µm (A & B), 0.5µm (C & D), and 250nm (E).

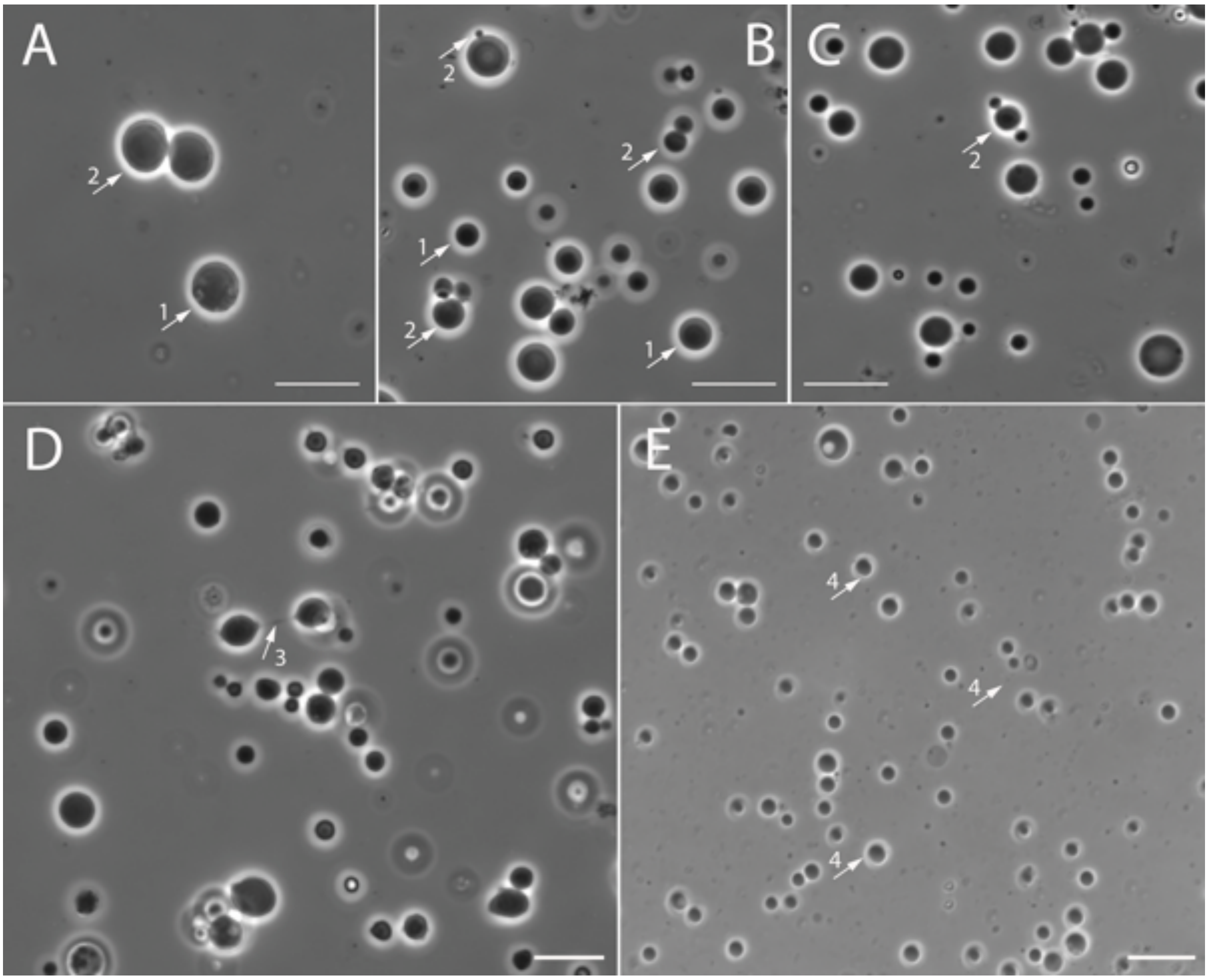


**Fig. S30. Reproduction by budding and binary fission.** Images A-E are the phase-contrast images of *EM-P* reproducing by budding or binary fission. The arrows in the images point to different stages of *EM-P* reproduction. The arrows in images B & C point to the cells undergoing budding. Scale bars: 10 µm.

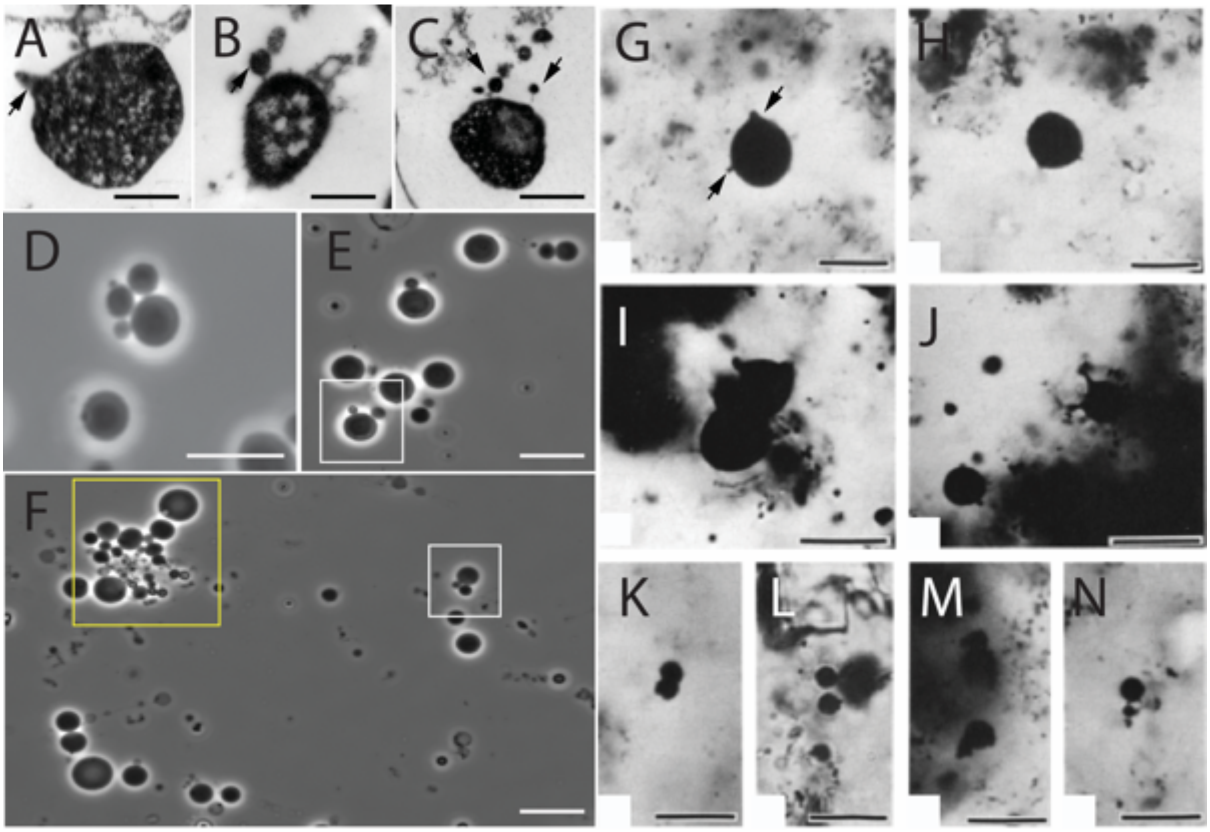


**Fig. S31. Morphological comparison between the North-Pole locality microfossils and *EM-P*.** A-C are TEM images of *EM-P* showing sequential stages of bud formation. Images D-E are phase-contrast images of *EM-P* cells that appear to be reproducing by budding or binary fission, depending on the size of the daughter cells. Images G-N show spherical microfossils reported from the North Pole locality (originally published by Buick *et al.*, 1990)(45). These microfossils resemble *EM-P* cells reproducing by budding. Arrows in images A & G point to pustular protuberances in *EM-P* and the North-Pole locality microfossils. Images E, F, I & K show *EM-P* cells and microfossils reproducing by binary fission. Image D and highlighted regions in images E & F (in white) show *EM-P* cells reproducing by the formation of multiple buds (also Fig 1B). Highlighted region in image F (in yellow) shows cells forming Y-junctions, similar to the microfossils shown in N. Scale bars: 500nm (A & B), 10µm (D-F), and 100µm (G-N).

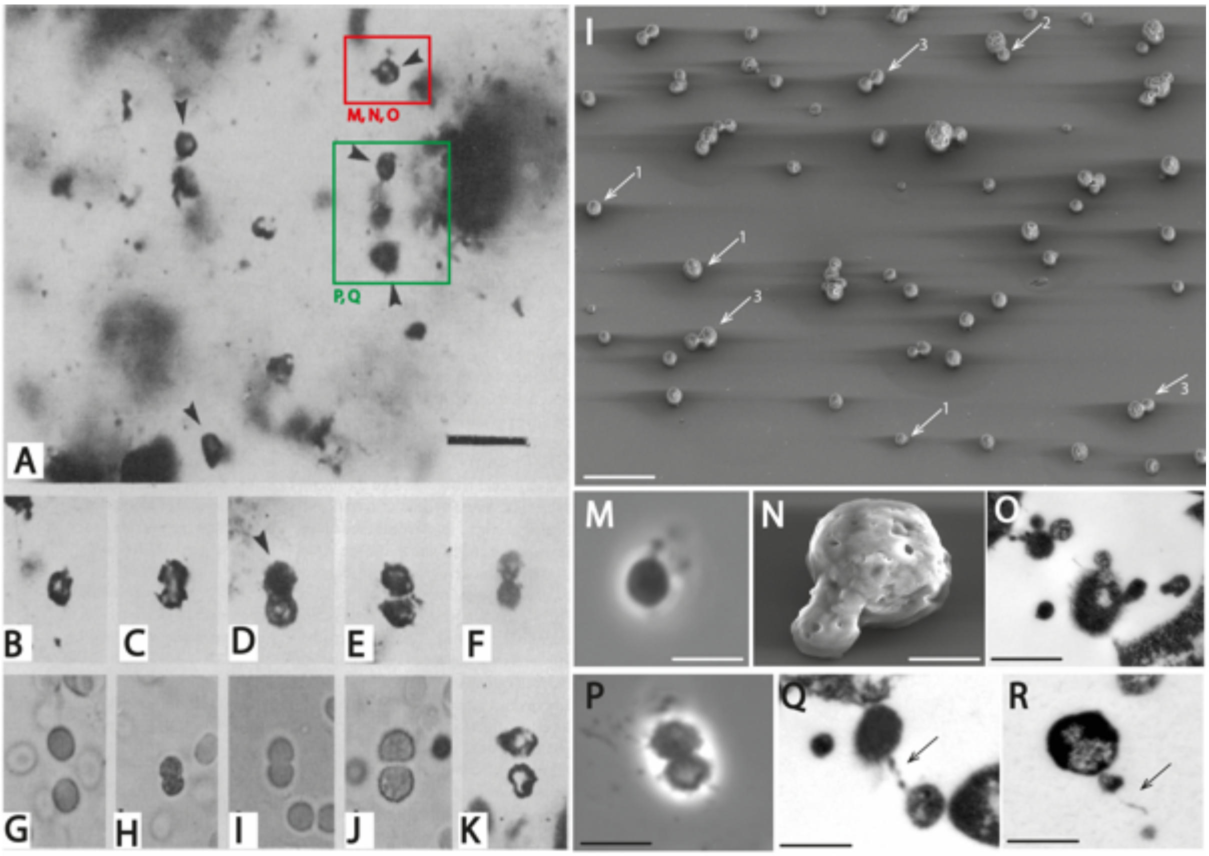


**Fig. S32. Comparison of *EM-P* with the Swartkoppie Formation microfossils.** Images A-K are the images of the Swartkoppie microfossils (originally published by Knoll *et al.,* 1977)(12). Images I-R are images of *EM-P* exhibiting morphological similarities with the Swartkoppie microfossils. Image A shows cells with membrane overhangs (red box), cells in dyads (green box), and individual spherical cells (arrows). Morphologically analogous *EM-P* cells can be seen in the images mentioned below in the red and green boxes. Images M, N, and O are phase-contrast, SEM, and TEM images of cells with membrane extensions, like the microfossil cell shown within the red box. Images P-R show a sequence of stages involved in *EM-P*'s cell division. Images B-F and G-K show the Swartkoppie microfossils reproducing by binary division. Morphologically similar *EM-P* cells were shown in image L (white arrows). Numbers next to the arrows indicate different stages of cell division. Scale Bar: 10 μm (A), 0.5μm (L & N), 0.25μm (O, Q & R) and 1μm (M & P).

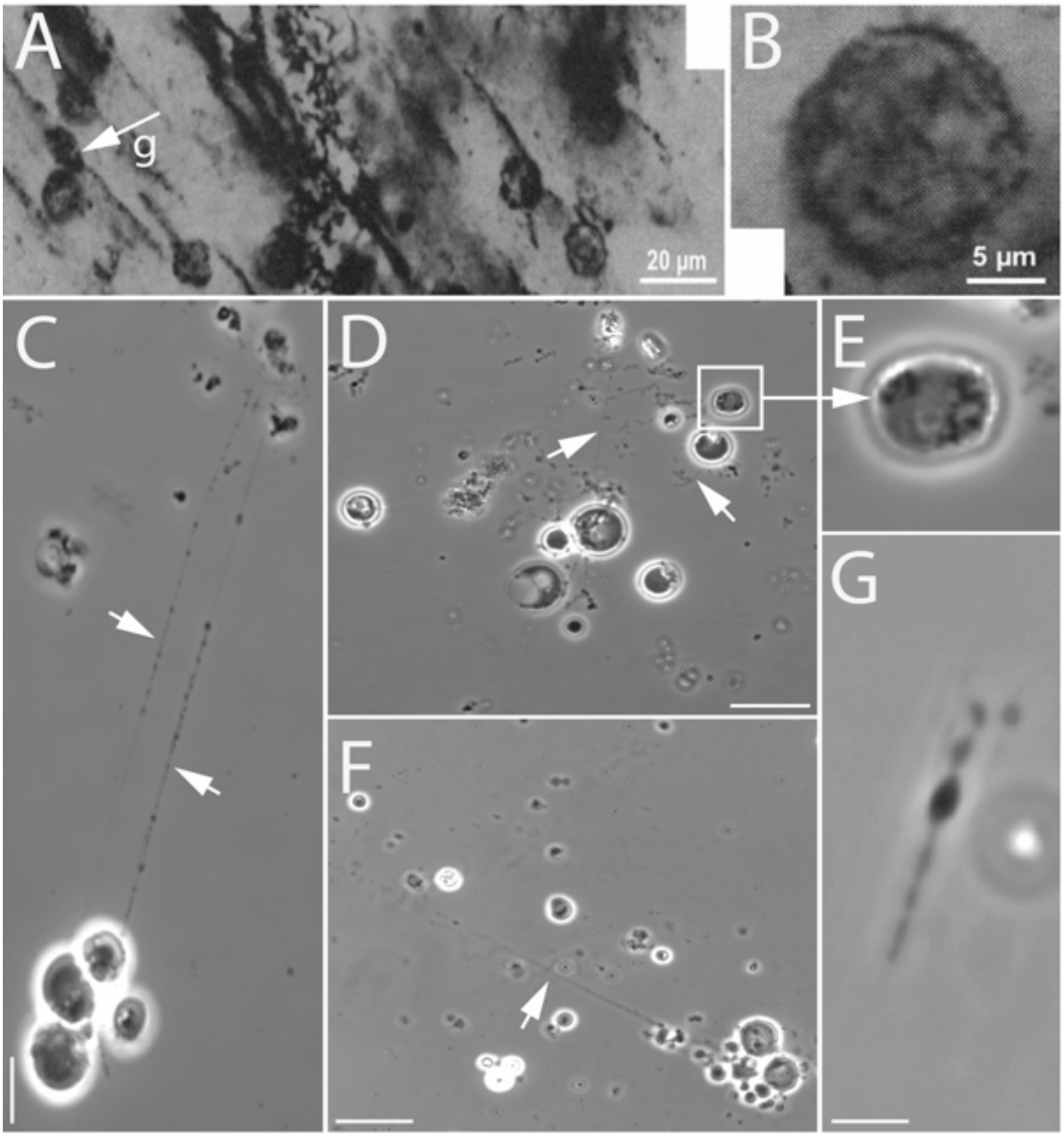


**Fig. S33. Comparison of *EM-P* with the Sheba Formation microfossils.** Images A & B show spherical microfossils reported from the Sheba Formation (originally published by Hickman-lewis *et al.,* 2018)(33). Image A shows the aggregation of spherical microfossils within organic carbon clasts. Image B shows a magnified image of spherical cells with unevenly distributed cytoplasm. Images C & G are morphologically analogous *EM-P* cells. Like the Sheba Formation, microfossils' spherical *EM-P* cells formed filamentous overhangs (arrows in images C-F, Movie S16). *EM-P* cells were also noticed to have uneven cytoplasm (E) (Fig. 2). Arrow in image-a points to spindle-like cells with filamentous extensions; morphologically similar *EM-P* cell is shown in image G. Scale bars: 10μm (C).

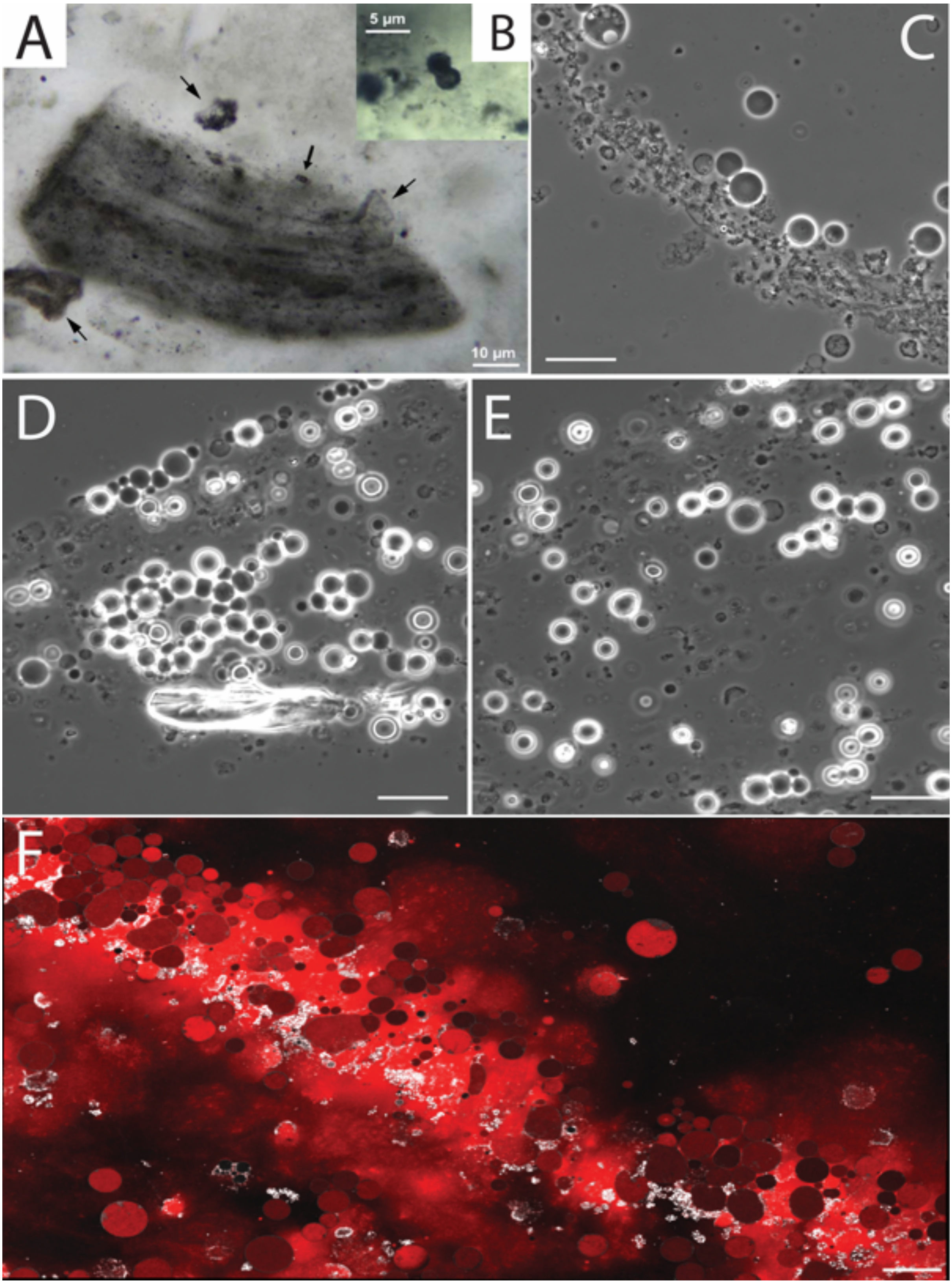


**Fig. S34. Comparison of *EM-P* with the Sheba Formation microfossils.** Image A & B show spherical microfossils reported from the Sheba Formation (originally published by Homann *et al.,* 2019)(46). Image B shows microfossils reproducing by what appears to be binary fission. Images C-F are the phase-contrast and STED microscope images of morphologically similar *EM-P* cells. Images C-E show spherical *EM-P* cells that are surrounded by membrane debris. Image F shows such debris composed of extracellular DNA (red). Scale bar: 10μm.

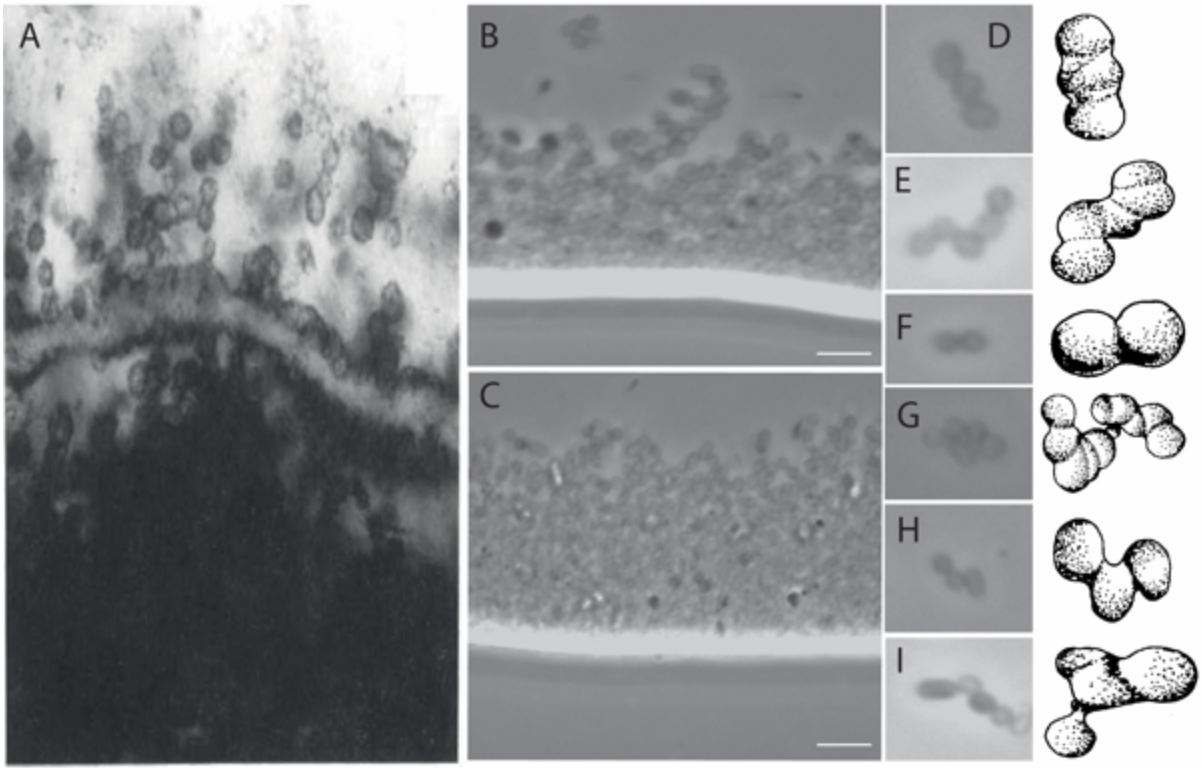


**Fig. S35. Comparison of the Onverwacht group microfossils with *EM-P*.**
Image A shows the Onverwacht microfossils (originally published by Walsh et al., 1992). Images C & D show sub-micrometer size cells *EM-P*, exhibiting morphological similarities with the Onverwacht microfossils. Images D-I show a cluster of *EM-P* cells organized in different configurations and interpretive drawings of morphologically similar Onverwacht microfossils (originally published by Walsh et al., 1992)(6). Scale bars: B & C (5μm).

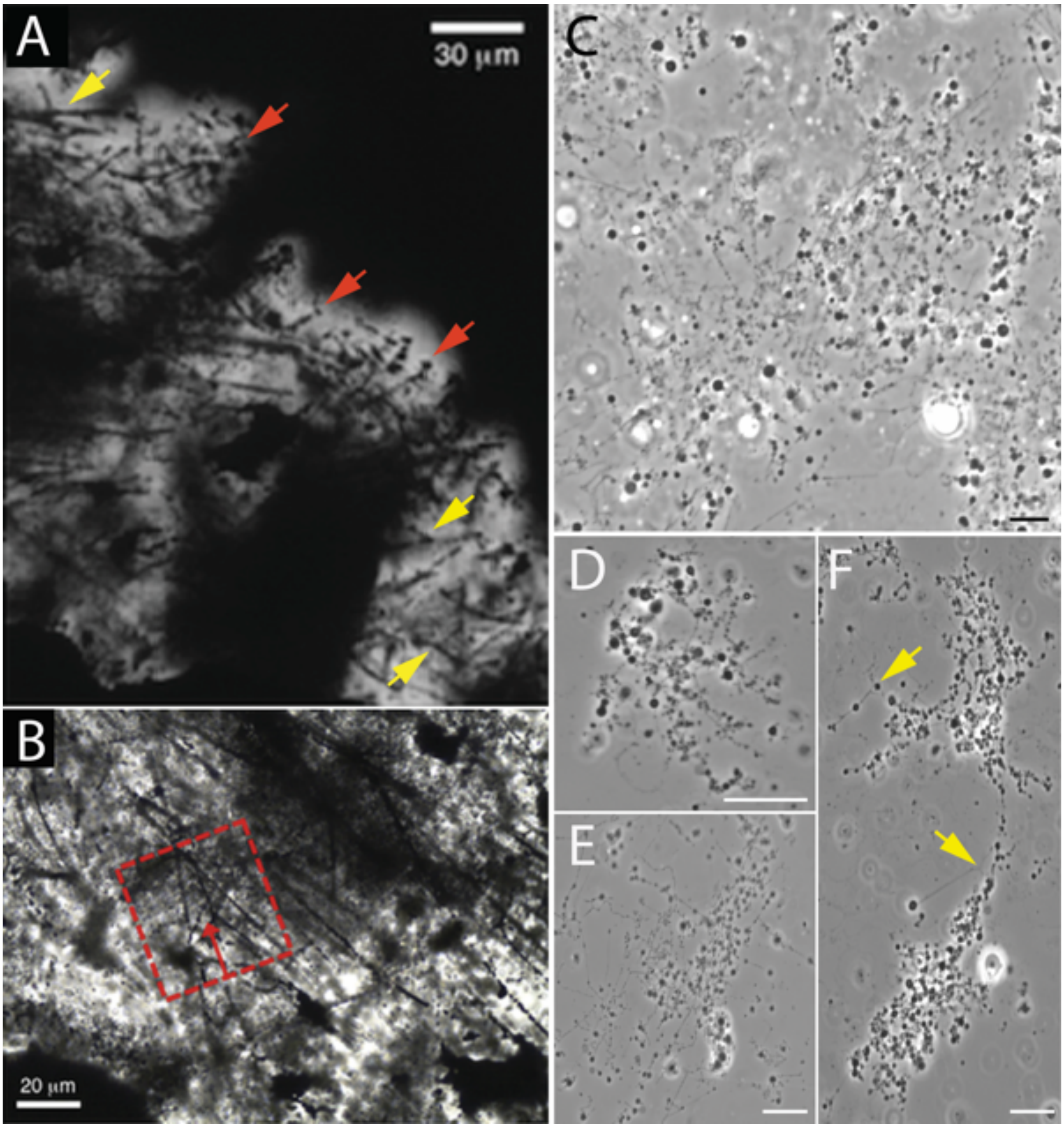


**Fig. S36. Morphological comparison between *EM-P* and the Sulphur Spring Formation microfossils.** Images A & B are microfossils reported from the Sulphur Spring Formations (originally published by Wacey *et al.,* 2014)(17). Red arrows in these images point to filamentous structures with spherical inclusions. Images C-F are morphologically analogous to *EM-P* cells. Similar to images A & B, *EM-P* exhibited strings of spherical daughter cells. Arrows in these images point to the branching of the filaments. Scale bars: 10µm (C-F).

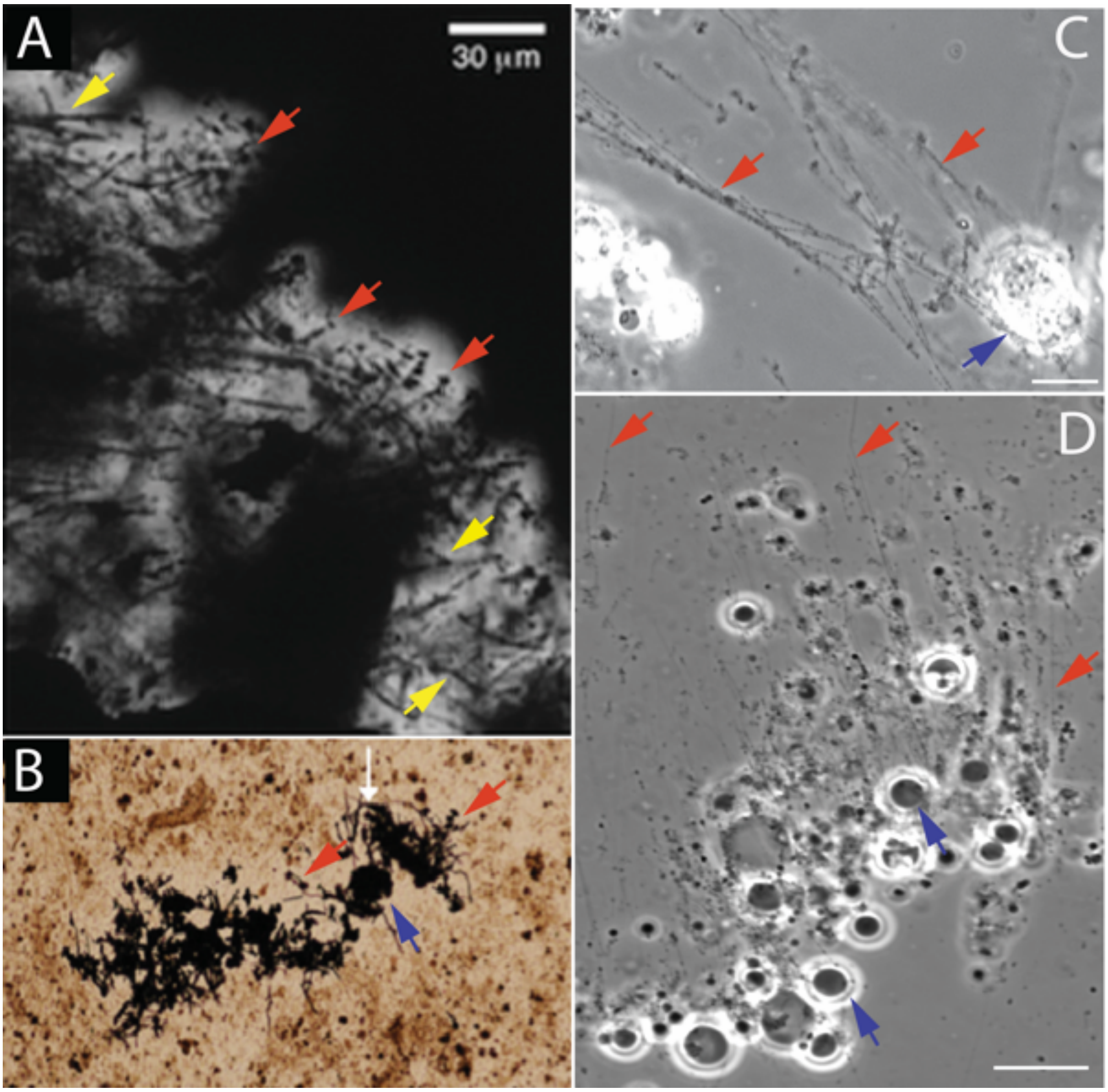


**Fig. S37. Morphological comparison between *EM-P* and the Sulphur Spring Formation. microfossils.** Images A & B are microfossils reported from Sulphur Spring Formations (originally published by Wacey *et al.,* 2014)(17). Red arrows in these images point to filamentous structures with spherical inclusions. The purple arrow in image B points to clusters of spherical organic structures from which filamentous structures appear to have originated. Yellow arrows point to the branching within the filamentous structures. Images C & D show morphologically similar *EM-P* cells. Images show spherical *EM-P* cells with filamentous extensions. Most filamentous extensions contain spherical daughter cells. Scale bars: 10µm (B & C).

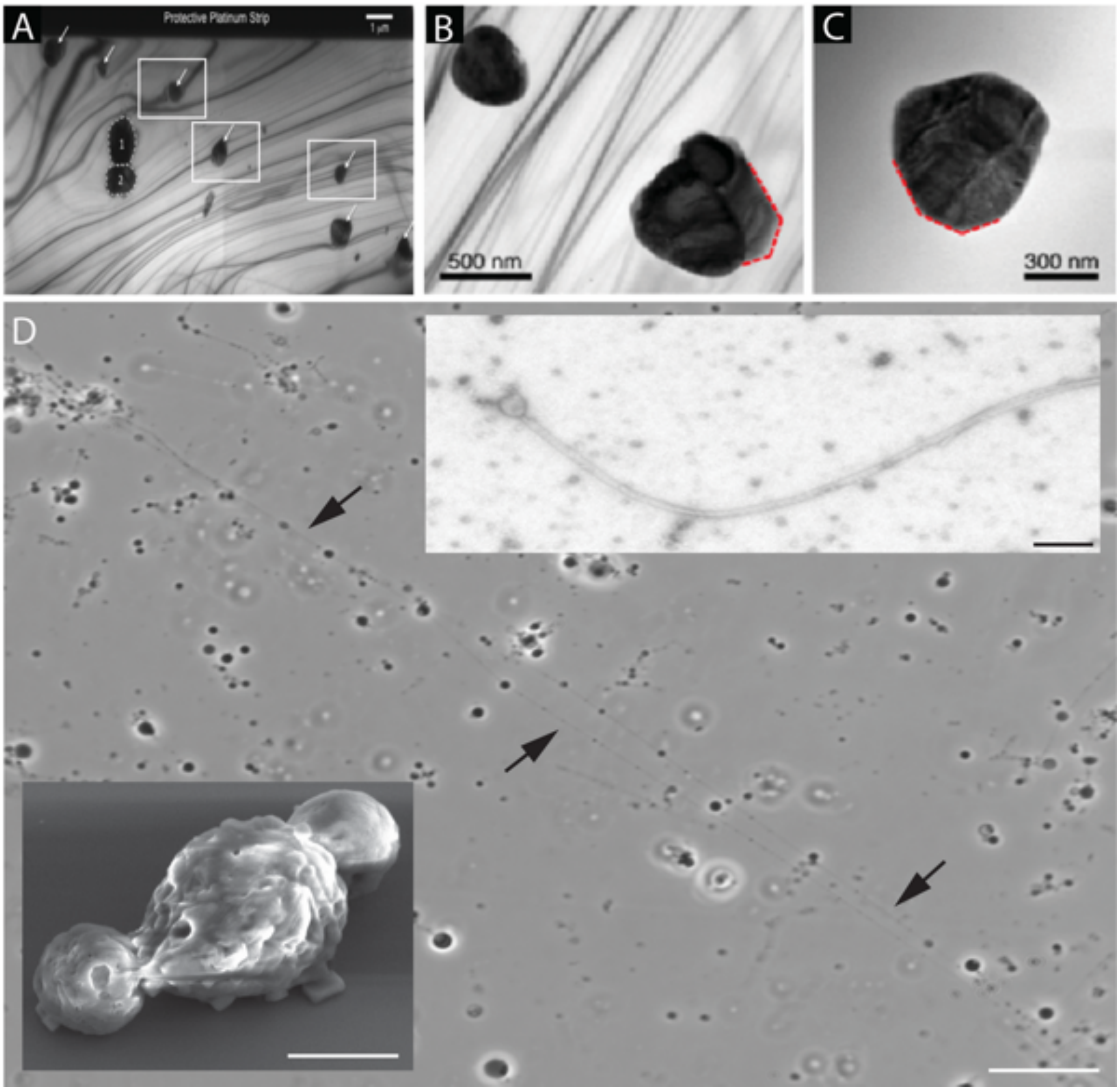


**Fig. S38. Morphological comparison between *EM-P* and the Sulphur Spring Formation. microfossils.** Images A-C are microfossils reported from the Sulphur Spring Formations (originally published by Wacey *et al.,* 2014)(17). The boxed regions in image A show spherical pyrite-encrusted microfossils within hollow filaments. Images B & C show close-ups of spherical structures with wrinkly surfaces. Image D shows hollow filaments with spherical inclusions morphologically analogous to the Sulphur Spring Formation microfossils (Scale bar: 10µm). Arrows in the image point to spherical daughter cells within the filaments. Insert at the bottom of the image shows an SEM image of such a cell with a wrinkled surface (Scale bar: 1µm). The Insert image in the top right corner of image D shows a TEM image of such hollow filaments (Scale bar: 500nm).

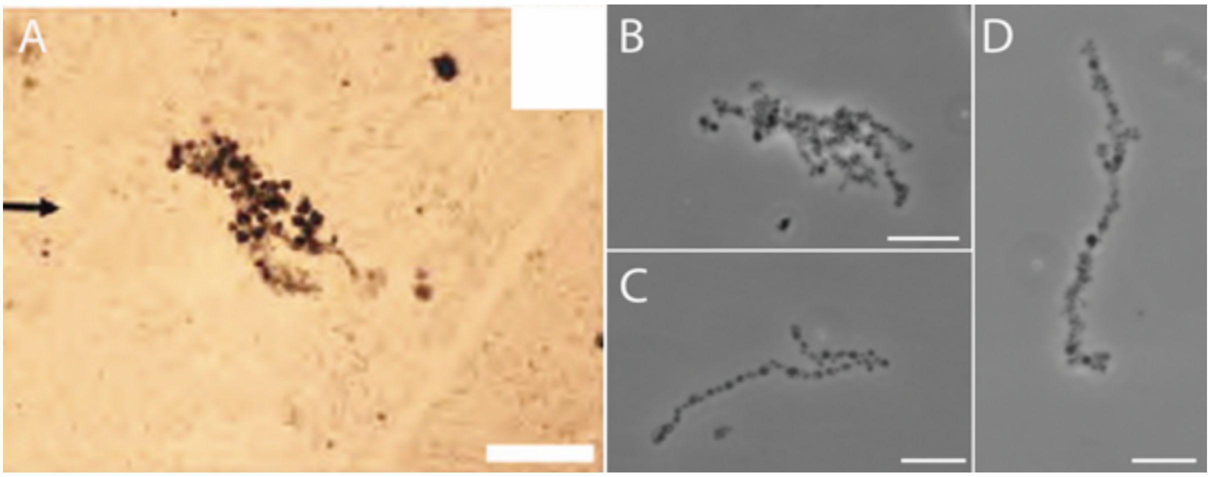


**Fig. S39. Morphological comparison between the Mt. Grant microfossils with *EM-P*.** Images A & B show microfossils reported from the Mt. Grant Formation (originally published by Sugitani *et al*., 2007)(37). Images B-D are morphologically analogous to *EM-P* daughter cells. Morphologically analogous *EM-P* cells to shown in image A are also shown in motion in Movie S14. Scale bars: 100µm (A) & 10µm (B-D).

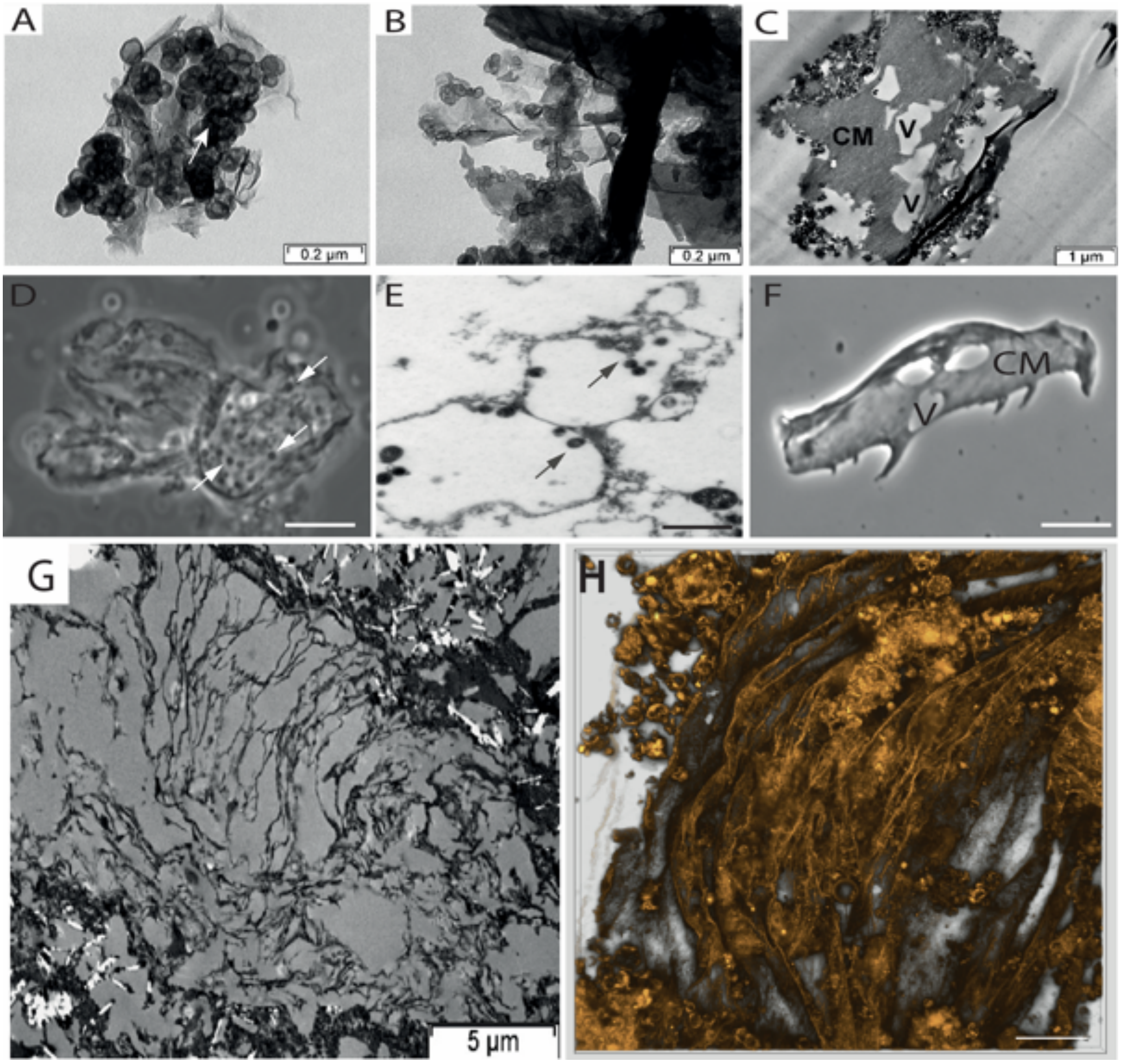


**Fig. S40. Morphological comparison between the Sulphur Spring Formation microfossils and *EM-P.*** Images A-C & G are organic structures reported from the Sulphur Spring Formation (originally published by Duck *et al*., 2007)(79). Images D-F are Phase-contrast (D & E) and TEM (E) images of morphologically analogous structures formed by *EM-P.* Images D & E show daughter cells daughter cells attached to membrane debris. CM and V in C & F stands for Cell Membrane and Vacuole respectively. The movement of daughter cells attached to membrane debris can be seen in Movie 17. Image H shows the 3D reconstituted confocal image of folded membrane debris similar to the ones reported from the Sulphur Spring Formation. Scale bars: 10µm (C & E) and 200nm (D).

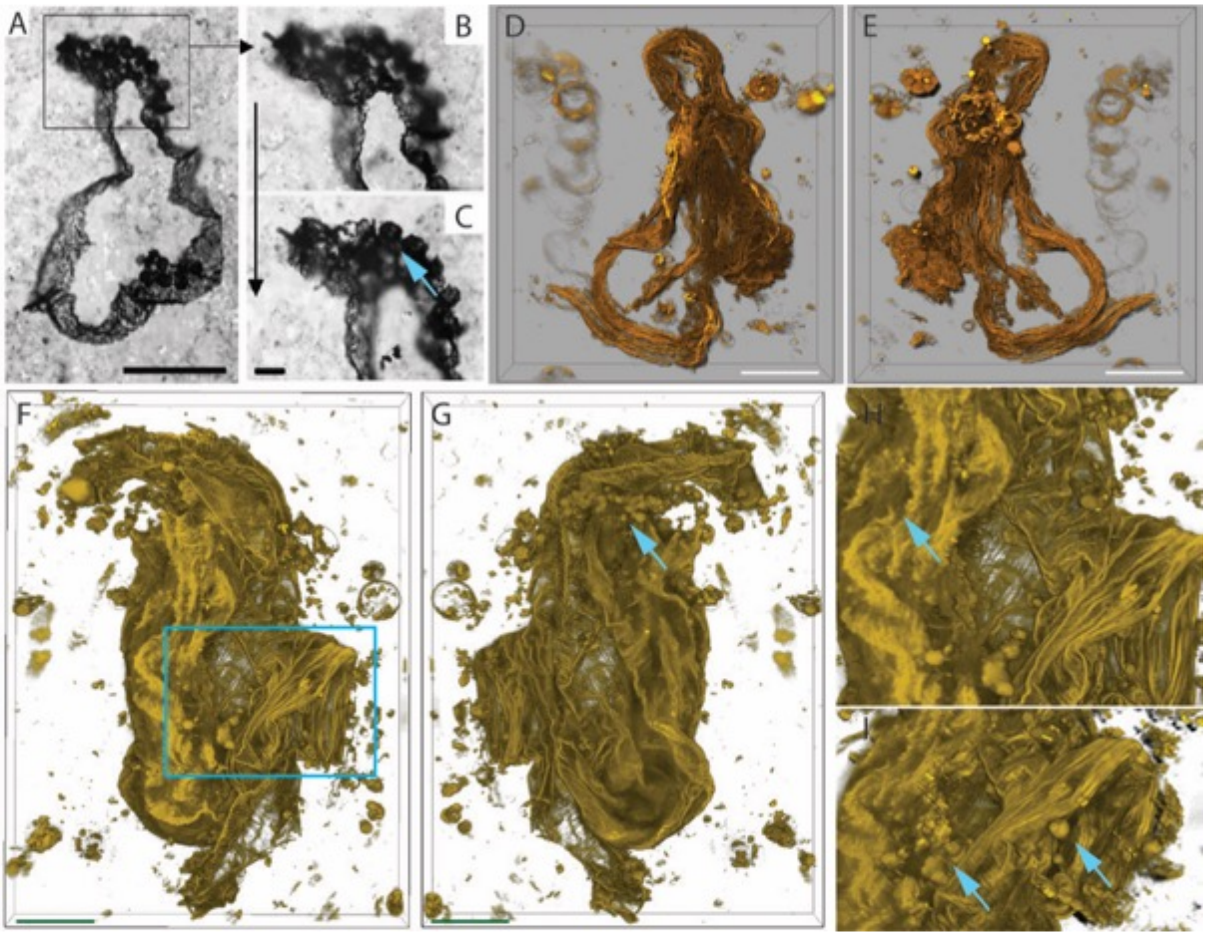


**Fig. S41. Morphological comparison between *EM-P* and the Mt. Goldsworthy microfossils.** Images A, B & C are organic structures reported from the Mt. Goldsworthy Formation (originally published by Sugitani *et al.,* 2007)(37). Images D & E, F & G, and H & I (magnifier region of image-F, highlighted in cyan box) are anterior and posterior views of morphologically analogous membrane debris produced by *EM-P* cells. Both sets of images show a film-like membrane with clusters of hollow spherical attachments (Cyan arrows). Images H & I show flexible film-like membrane debris of *EM-P* with similar spherical attachments. Scale bars: 50µm (A) and 10µm (D-G).

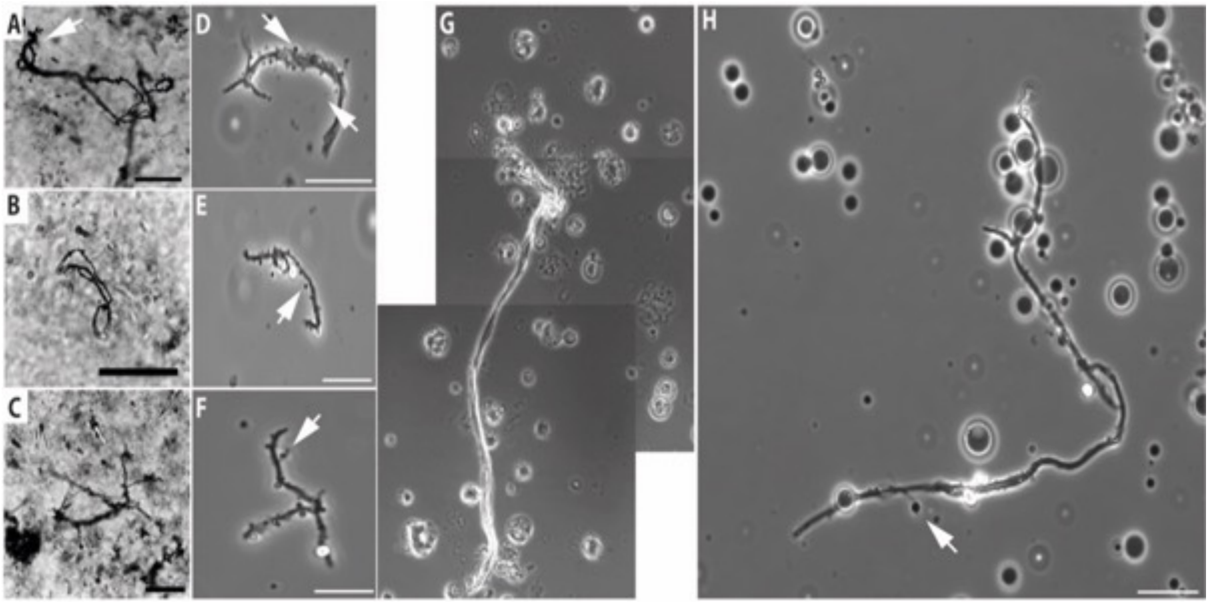


**Fig. S42. Morphological comparison between *EM-P* and the Mount Grant microfossils.** Images A, B & C are organic structures reported from the Mount Grant Formation (Sugitani *et al.,* 2007)(37). Scale bar: 20µm (A-C). Images D & H are morphologically analogous to *EM-P's* membrane debris. Arrows in the images point to spherical inclusions attached to thread-like membrane debris. Membrane debris in images G & H should have formed by lysis and collapse of either individual or cluster of large spherical cells of *EM-P* with hexagonal vesicles (Fig. S43). Scale bars: 10µm (D-H).

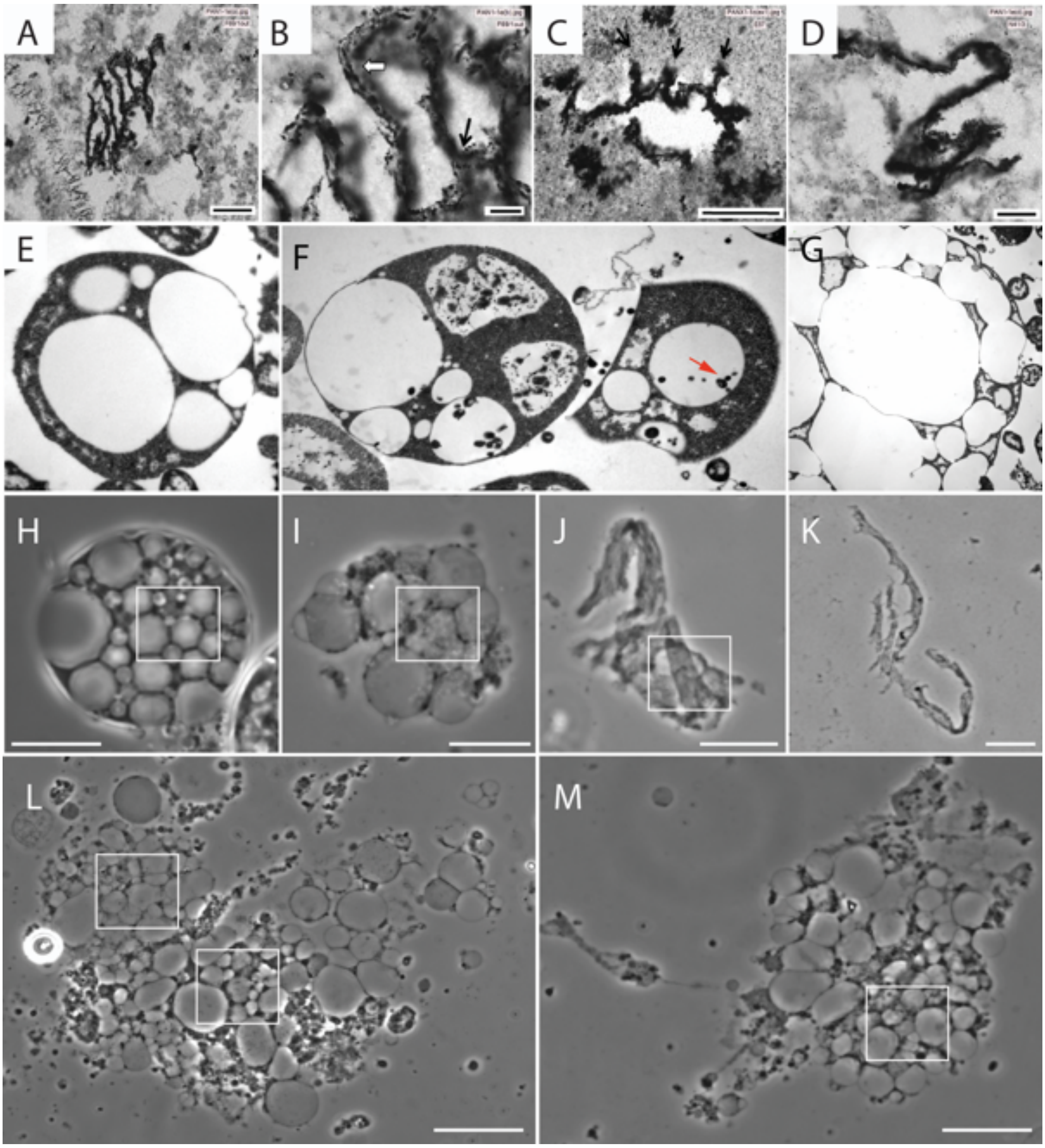


**Fig. S43. Morphological comparison between *EM-P* and the SPF organic structures.** Images A-D are organic structures reported from the Mount Grant Formation (Sugitani *et al.,* 2013)(39). Scale bar: 20µm (A-C). Images E-G are TEM images of morphologically analogous *EM-P* cells. Arrows in the images B & F point to spherical inclusions attached to thread-like membrane debris. Images H-M show the step-by-step transformation of *EM-P* cells into polygonal cell beds. Boxed regions within these images show the presence of a polygonal honeycomb-like structure within the cell debris. Scale bars: 1µm (E-G) & 10µm (H-M).

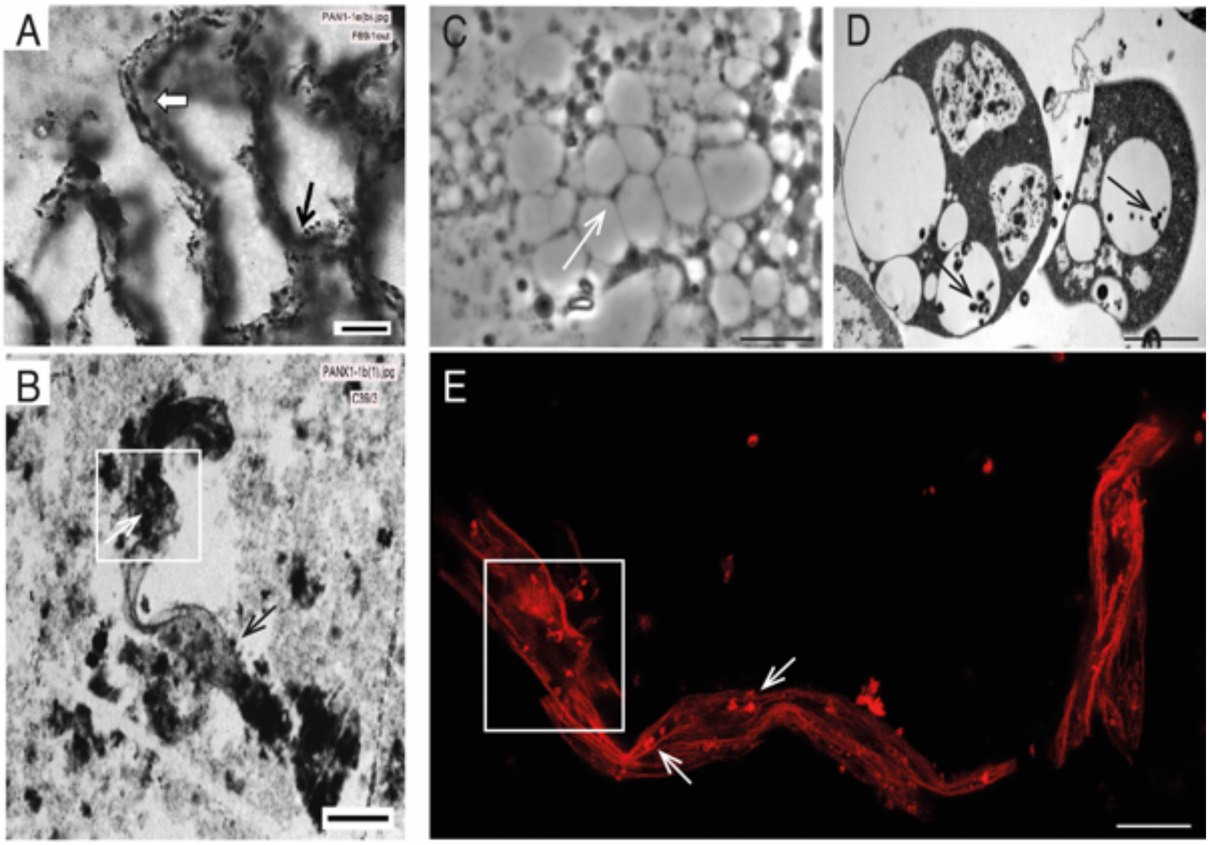


**Fig. S44. Morphological comparison of the SPF microfossils with *EM-P*.**

Images A & B are organic structures reported from the SPF (Sugitani *et al.,* 2013)(39). Scale bars in images A & B are 20µm & 50µm, respectively. Image C-D are morphologically analogous to *EM-P* cells or their membrane debris (E). Duel-walled honeycomb-like structures could be seen in both images A & C (white arrows). Black arrows in images A & D point to a string of spherical daughter cells attached to the walls of the honeycomb. Boxed regions in images B & E show similarities between film-like debris reported from SPF and membrane debris observed in *EM-P* incubations. These structures have spherical daughter cells with a membrane wrapping (arrows). Scale bar: 10µm (C&E) & 1µm (D).

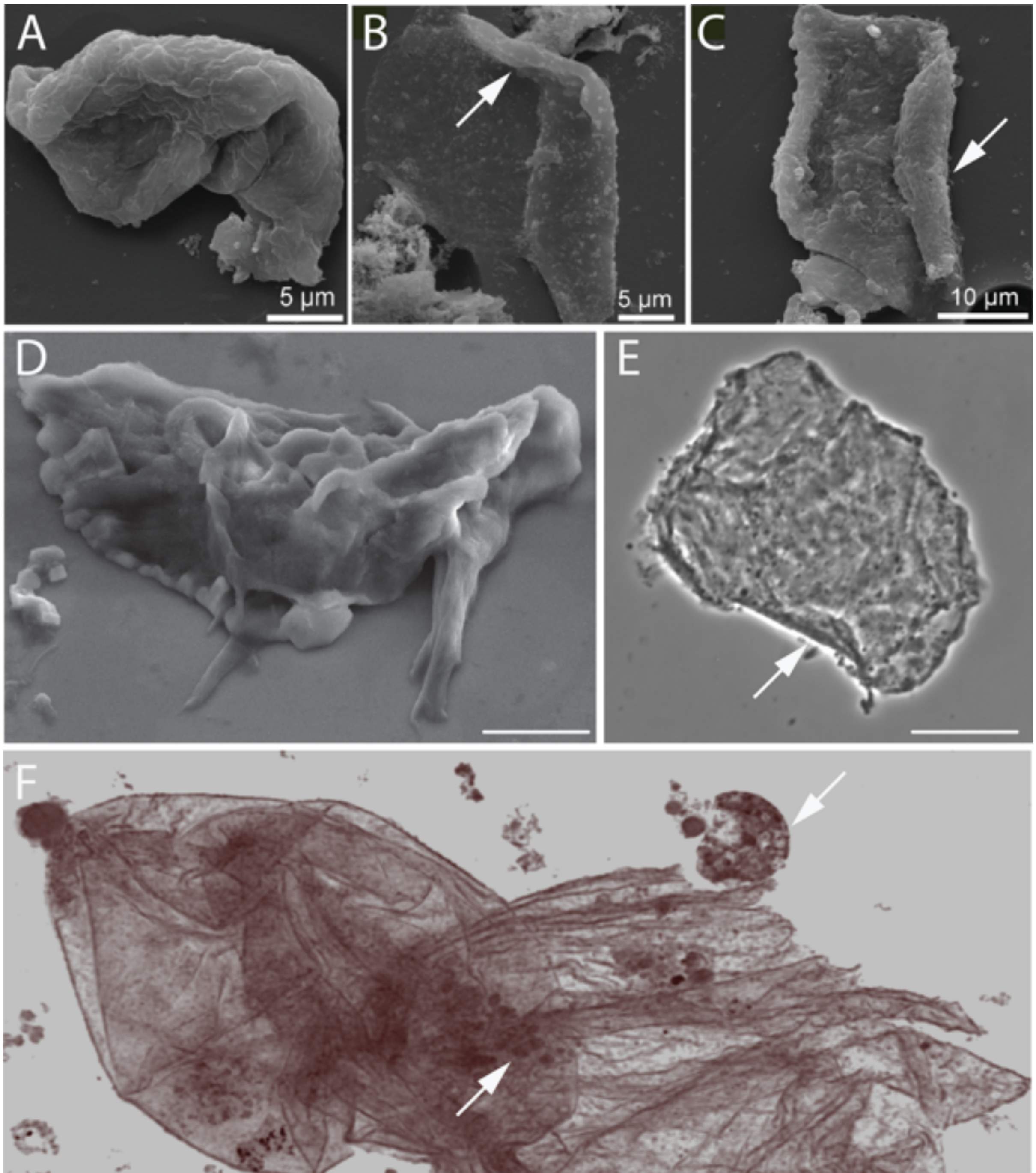


**Fig. S45. Morphological comparison between *EM-P* membrane debris and SPF membrane debris.** Images A-C are organic structures reported from SPF formation (originally published by Delarue *et al.,* 2019)(35). These images show folded membrane-like structures. Images D, E & F are SEM, phase-contrast, and 3D-rendered STED images of morphologically analogous *EM-P's* membrane debris. Arrows in images B, C & E point to the bending of the membrane at the edges in both SPF organic structures and *EM-P*. Arrows in image F point to spherical daughter cells of *EM-P* enclosed within the membrane debris. Scale bars: 2µm (D), 10µm (E) and 20µm (F).

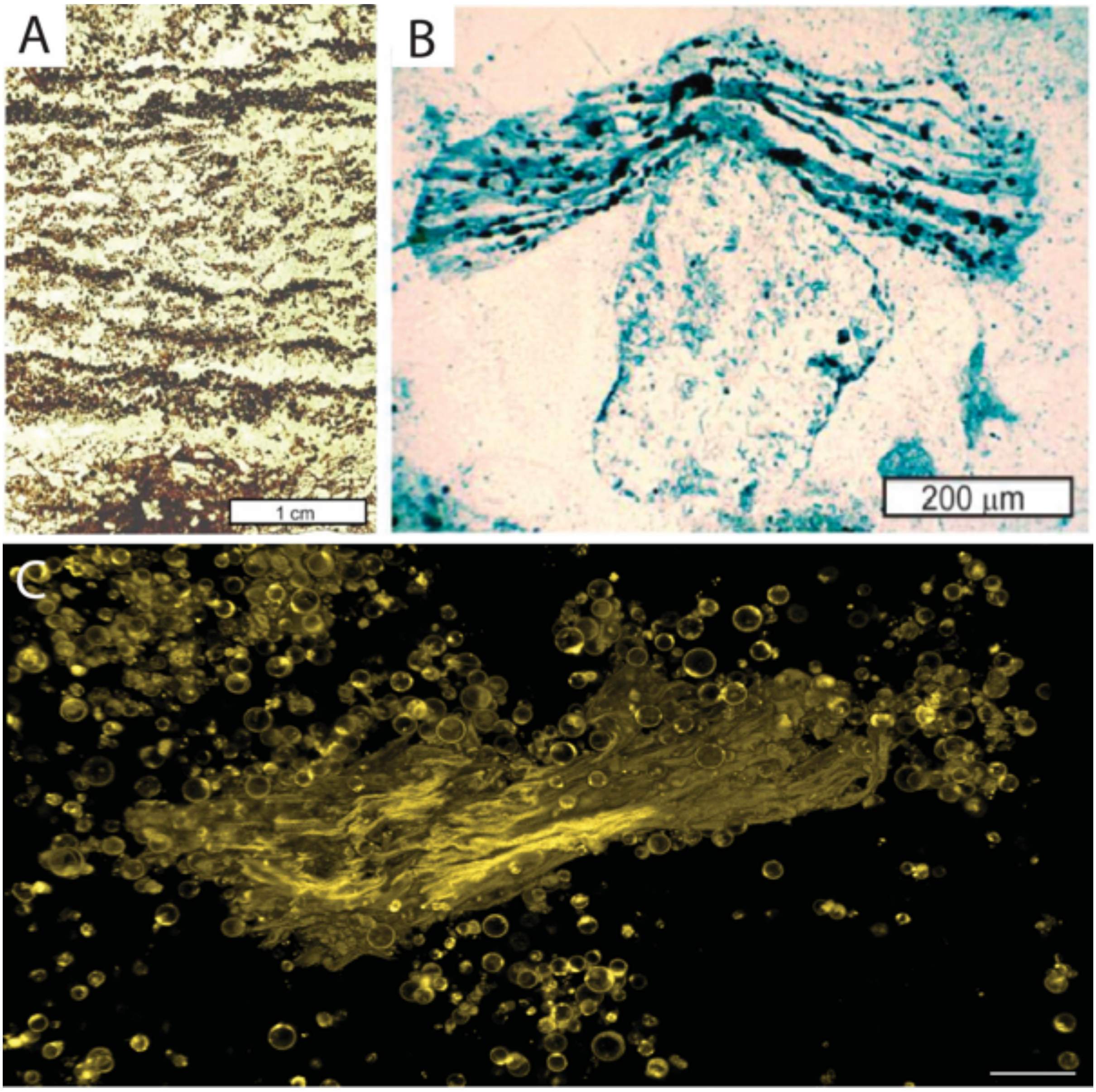


**Fig. S46. Morphological comparison between *EM-P* and the Farrel Quartzite film-like structures.** Images A & B are organic structures reported from the Farrel quartzite formation (Retallack *et al.,* 2016)(36). Both images show a folded film-like membrane with spherical inclusions. Image C shows a wrinkled membrane with spherical cells attached, similar in structures to the Farrel Quartzite microfossils. Scale bar: 20µm (C).

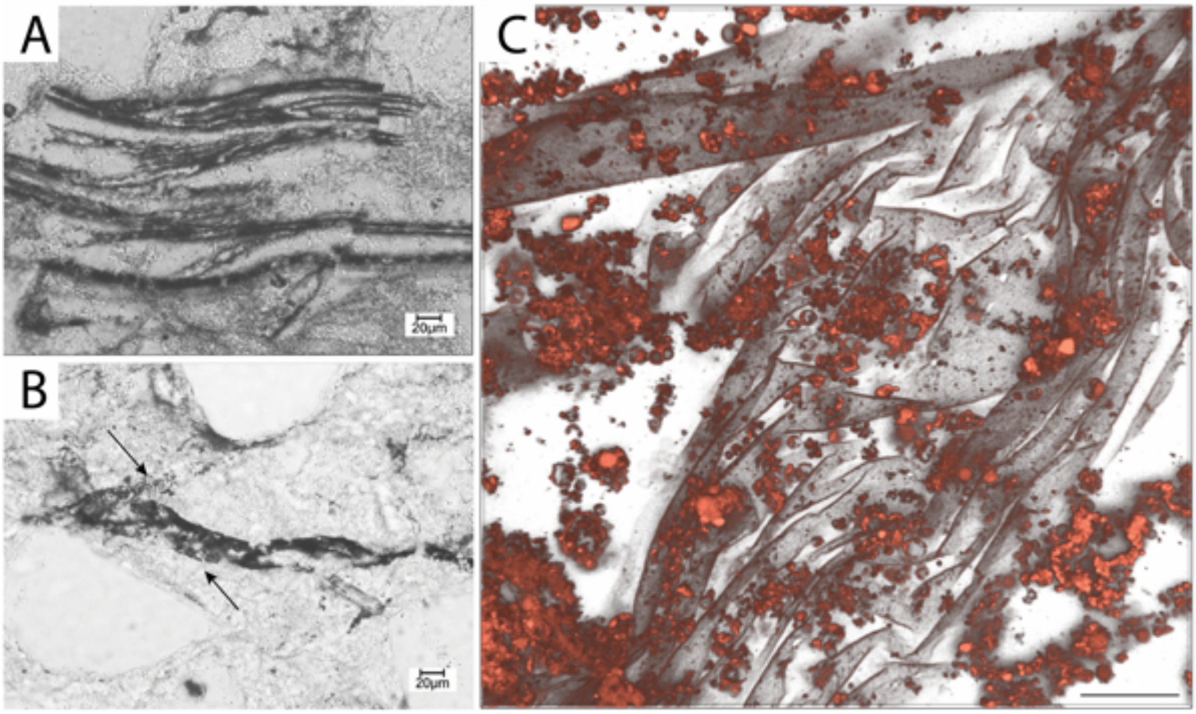


**Fig. S47. Morphological comparison between *EM-P* and the Moodies Group microfossils.** Images A & B are organic structures reported from the Moodies Group (Kohler *et al.,* 2019)(80). Both images show a folded film-like membrane with spherical inclusions (black arrows). Image C shows a wrinkled membrane with spherical cells attached to it, similar in structures to the Moodies microfossils. Scale bar: 10µm.

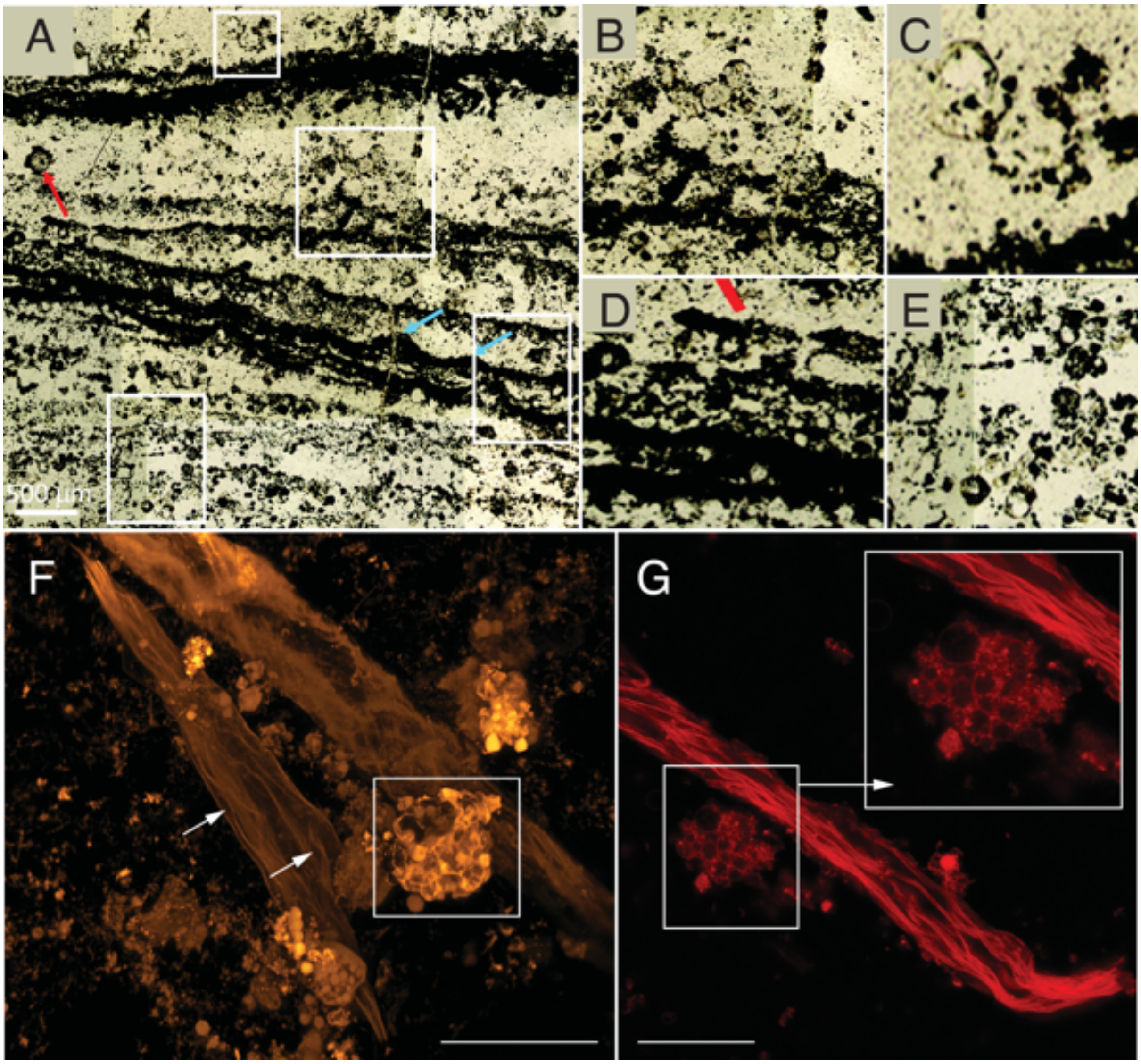


**Fig. S48. Morphological comparison of organic structures reported from the Dresser Formation with *EM-P*.** Images A-E show organic structures reported from the Dresser Formation (originally published by Wacey *et al.,* 2018)(10). It shows wavy filamentous structures (blue arrows) with individual (red arrow) or aggregations (box) of hollow spherical inclusions. Image B is the membrane debris formed by lysis of *EM-P*, which is analogous in its morphology to organic structures reported from the Dresser Formation. Cells and membrane debris in image F-H were stained with FM$^{TM}$5-95 (red) membrane stain. The boxed regions within A, F, & G show similar clusters of hollow spherical vacuoles. Scale bar: 10μm (F & G).

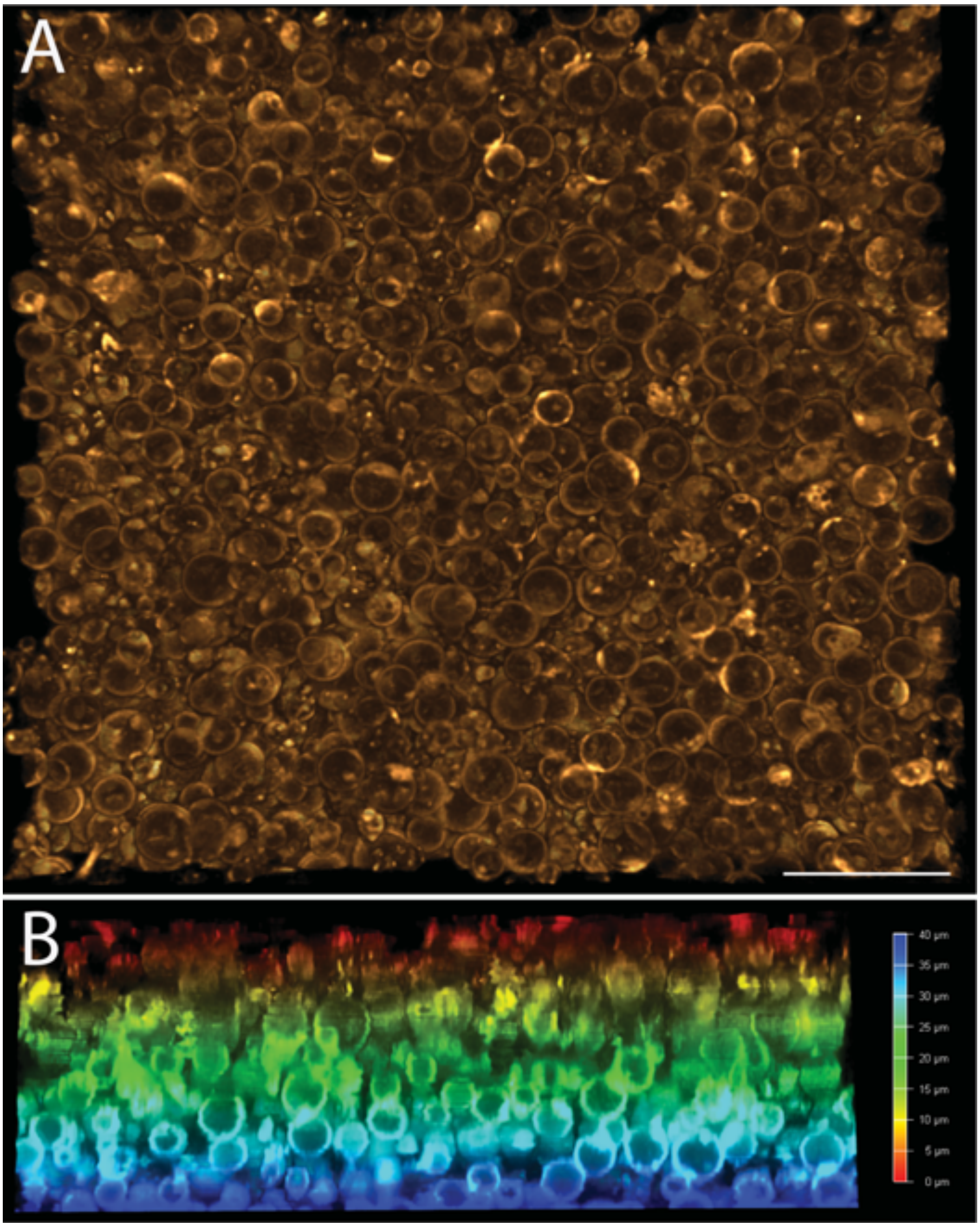


**Fig. S49. Multilayered *EM-P* cells.** Images A & B show a top and lateral view of *EM-P* cells growing in multiple layers at the bottom of the chamber slide. Cells in the above images were stained with membrane stain, FM$^{TM}$5-95. Scale bar: 20µm (A).

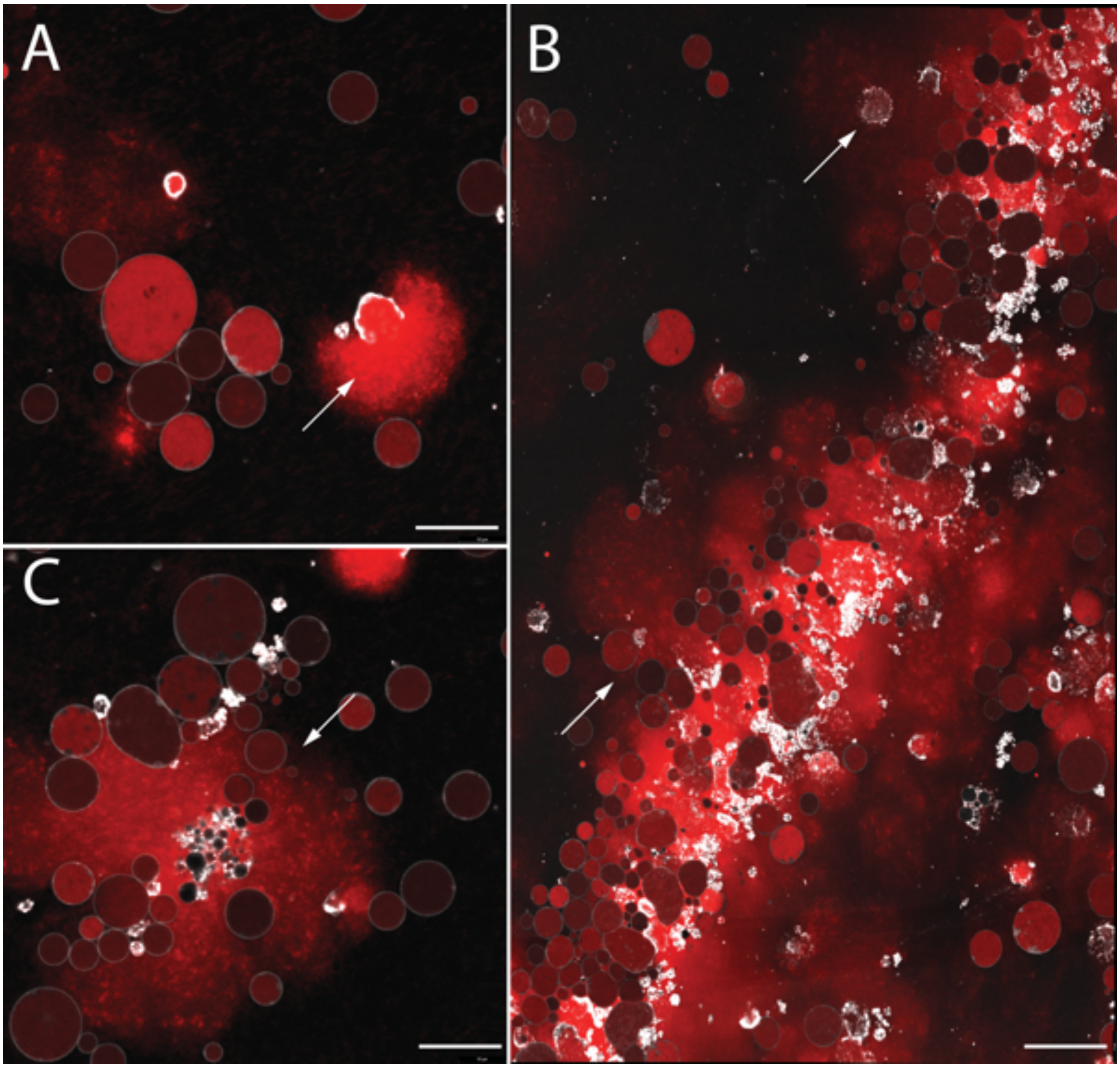


**Fig. S50. Cell lysis and release of DNA observed in *EM-P*.**

Images A & B show the lysis and release of cell constituents like DNA (red) into their surroundings. Image B shows the aggregation of cells within the biofilms with a considerable amount of extracellular DNA. Cells in these images were stained with FM$^{TM}$5-95 (membrane, white) and PicoGreen$^{TM}$ (DNA, red). Scale bars: 10µm.

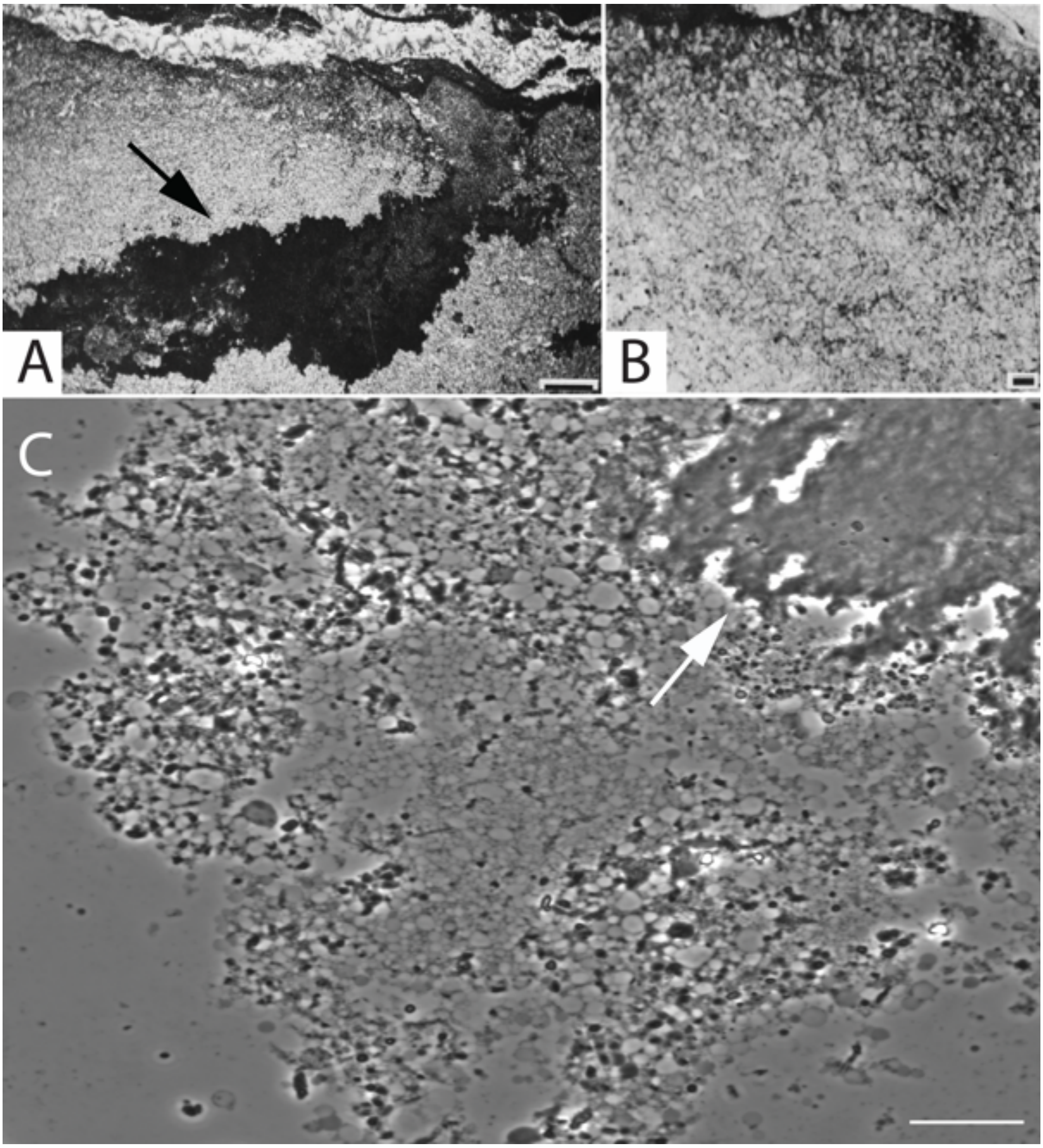


**Fig. S51. Morphological comparison of the North Pole locality microfossils and *EM-P*.** Images A & B are the organic structures reported from the North-pole locality (originally published by Buick *et al*., 1990)(45). Image D similar aggregation of spherical *EM-P* cells observed in our study. Arrows in images A & C point to the membrane debris between the cell aggregates. Further comparison of North Pole locality microfossils and *EM-P* cells is shown in Fig. S52 & S53. Scale bars: A (1mm), B (100µm), and C (20µm).

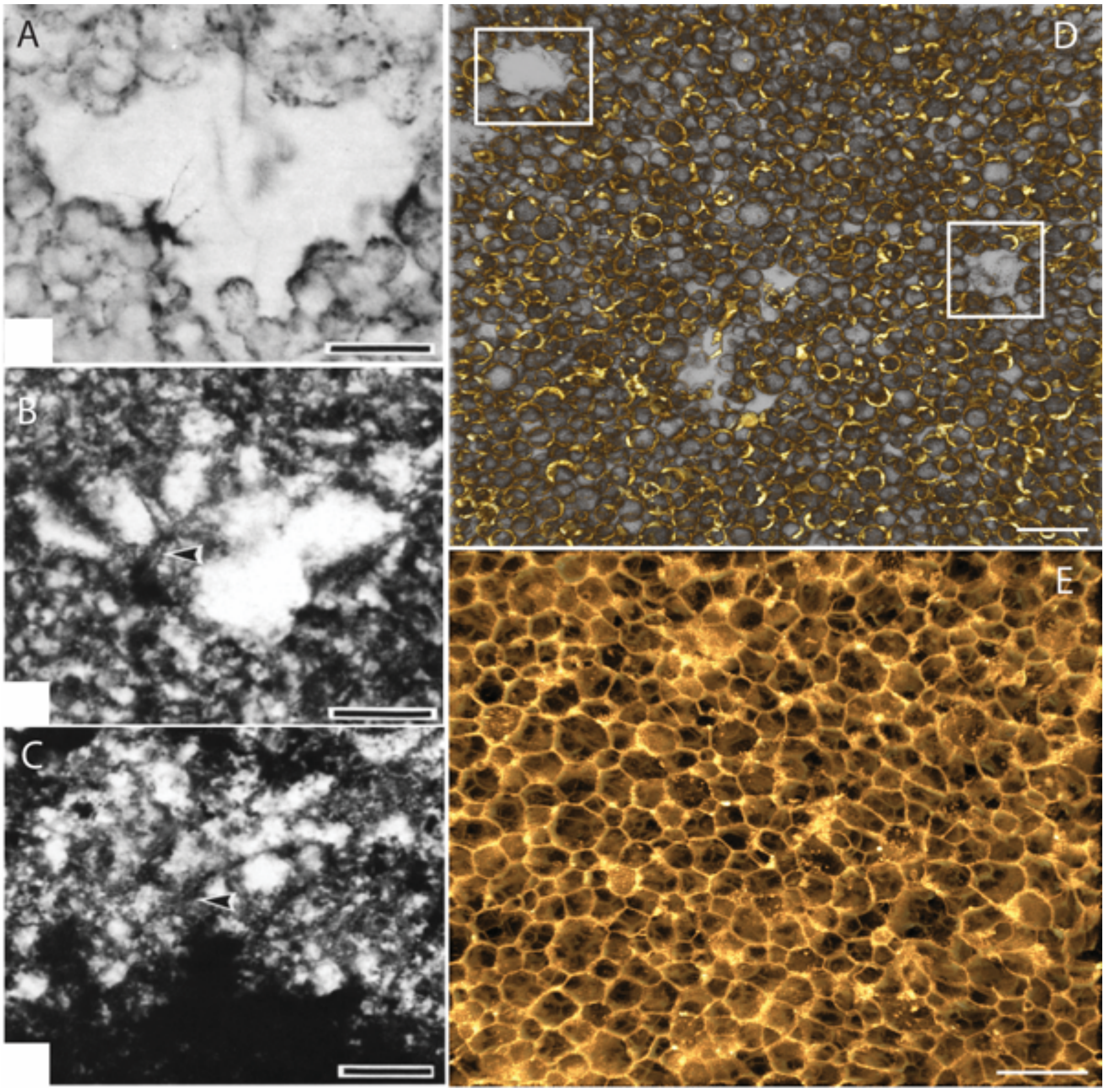


**Fig. S52. Morphological comparison of the North Pole locality microfossils and *EM-P*.** Images A-C are the organic structures reported from the North-pole locality (originally published by Buick *et al*., 1990)(45). They show organic mats composed of individual hollow spherical cells and large gaps within the mats (A & B). Images D & E are similar mat-like organic structures formed by *EM-P*. The highlighted regions in D show morphologically similar gaps observed in the North Pole locality and *EM-P* biofilms. Scale bars: D & E (10μm).

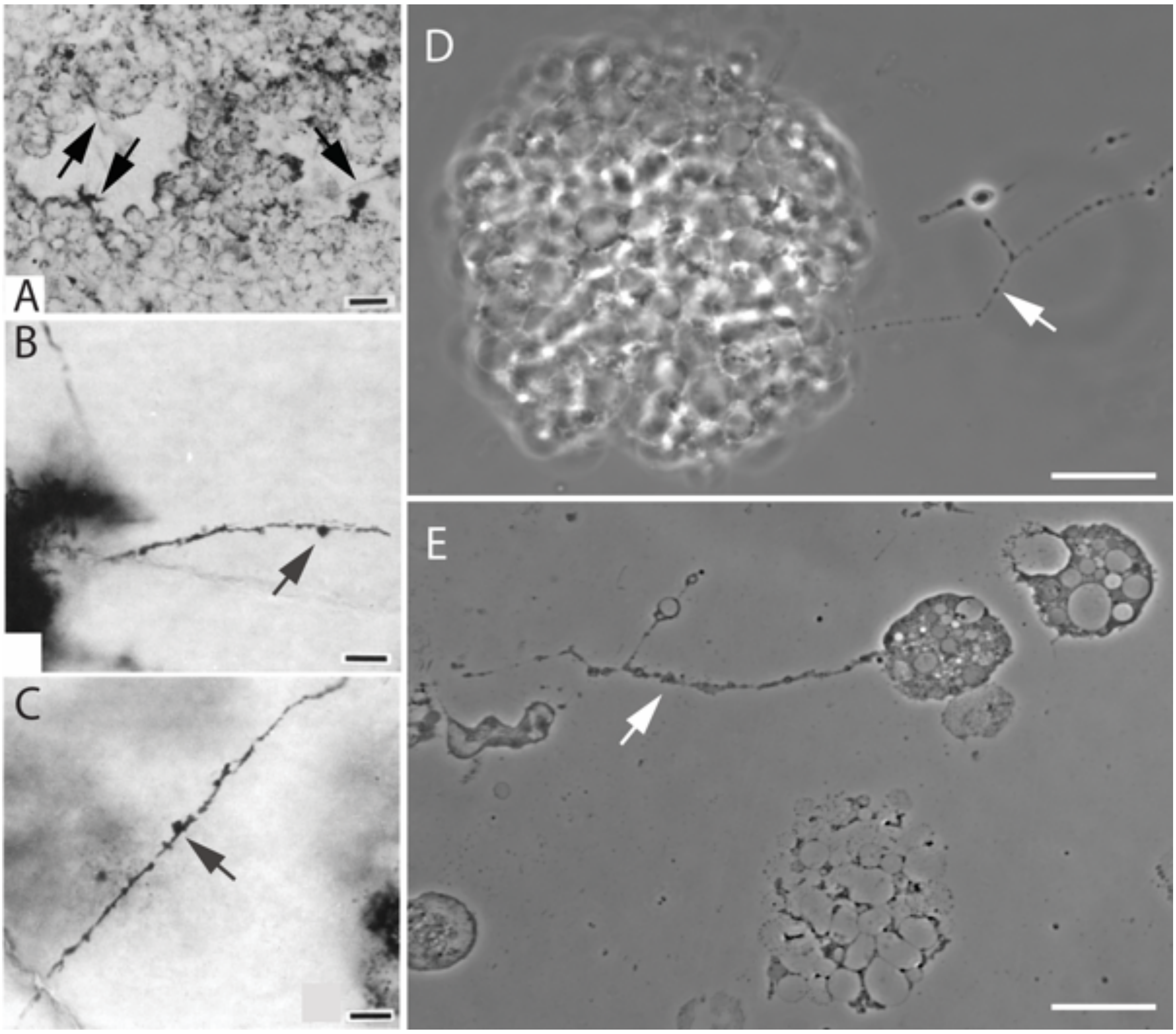


**Fig. S53. Morphological comparison of the North Pole locality microfossils and *EM-P*.** Images A-C are the organic structures reported from the North-pole locality (originally published by Buick *et al*., 1990)(45). Images D & E show phase-contrast images of *EM-P* cells. Image D shows an aggregation of hollow *EM-P* vesicles with filamentous structures. The method of their formation is shown in Movies 5-9. Scale bars: D & E (10μm).

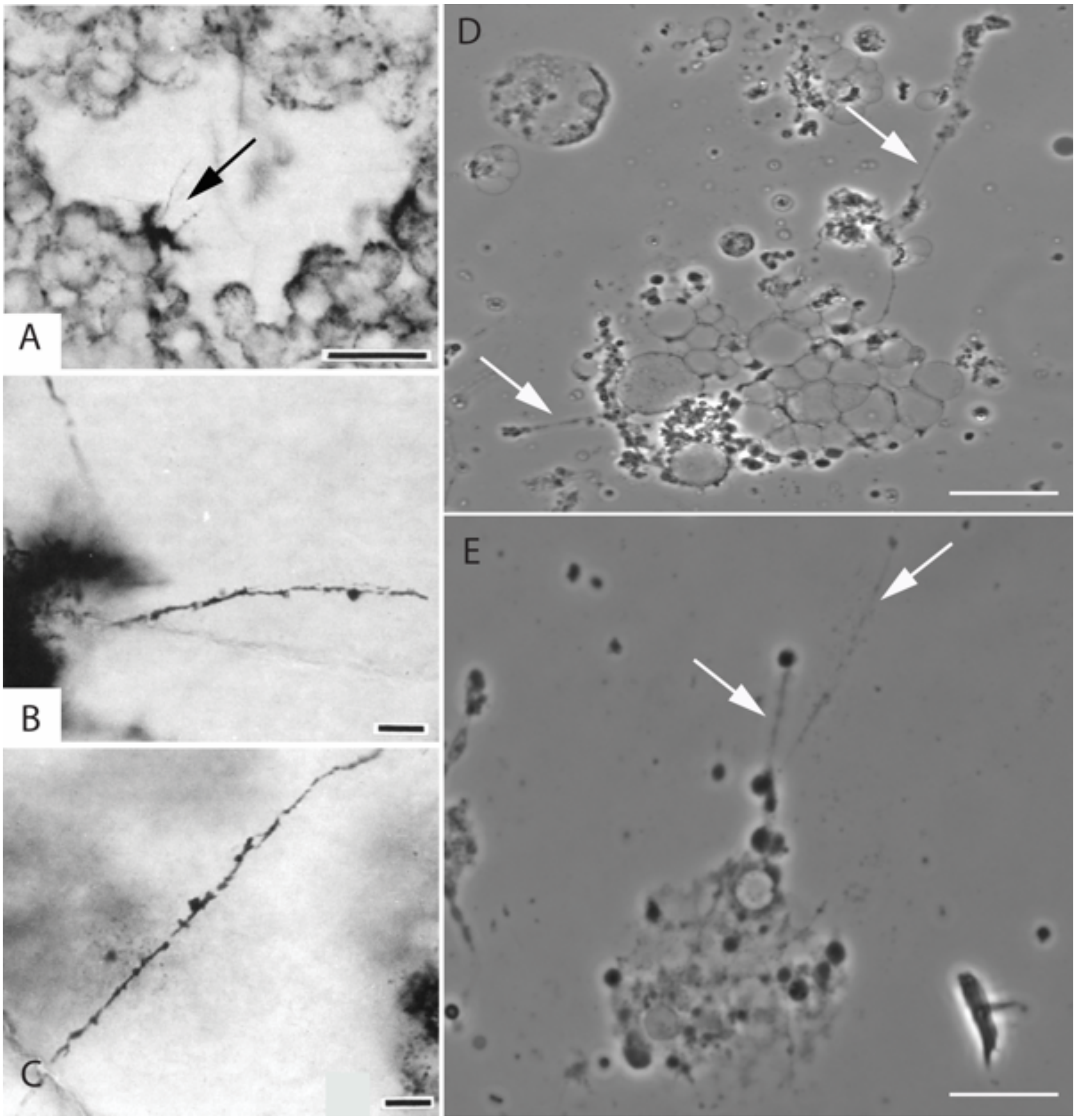


**Fig. S54. Morphological comparison of North Pole locality microfossils and *EM-P*.** Images A-C are the organic structures reported from the North-pole locality (originally published by Buick *et al*., 1990)(45). Images D & E show phase-contrast images of *EM-P* cells. Image D shows an aggregation of hollow *EM-P* vesicles with filamentous structures. The method of their formation is shown in Movies 5-9. Scale bars: D & E (10μm).

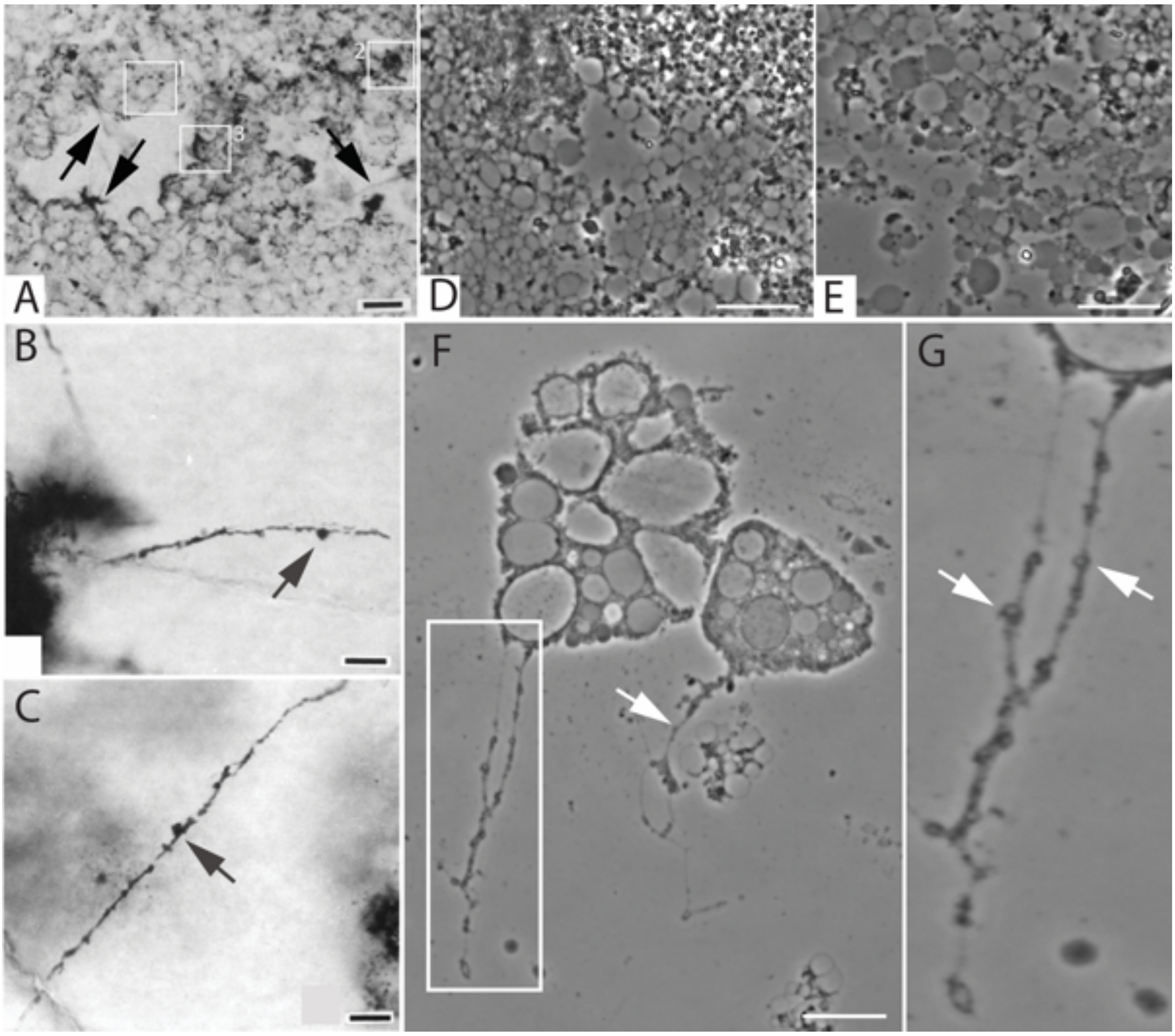


**Fig. S55. Morphological comparison of the North Pole locality microfossils and *EM-P*.** Images A-C are the organic structures reported from the North Pole locality (originally published by Buick *et al*., 1990)(45). Images D-G show phase-contrast images of *EM-P* cells. Image D shows an aggregation of hollow *EM-P* vesicles with filamentous structures. The method of their formation is shown in Movies 5-9. Arrows in images point to the similarities in the filamentous structure from both north-pole formation and *EM-P*. Scale bars: D & E (10µm).

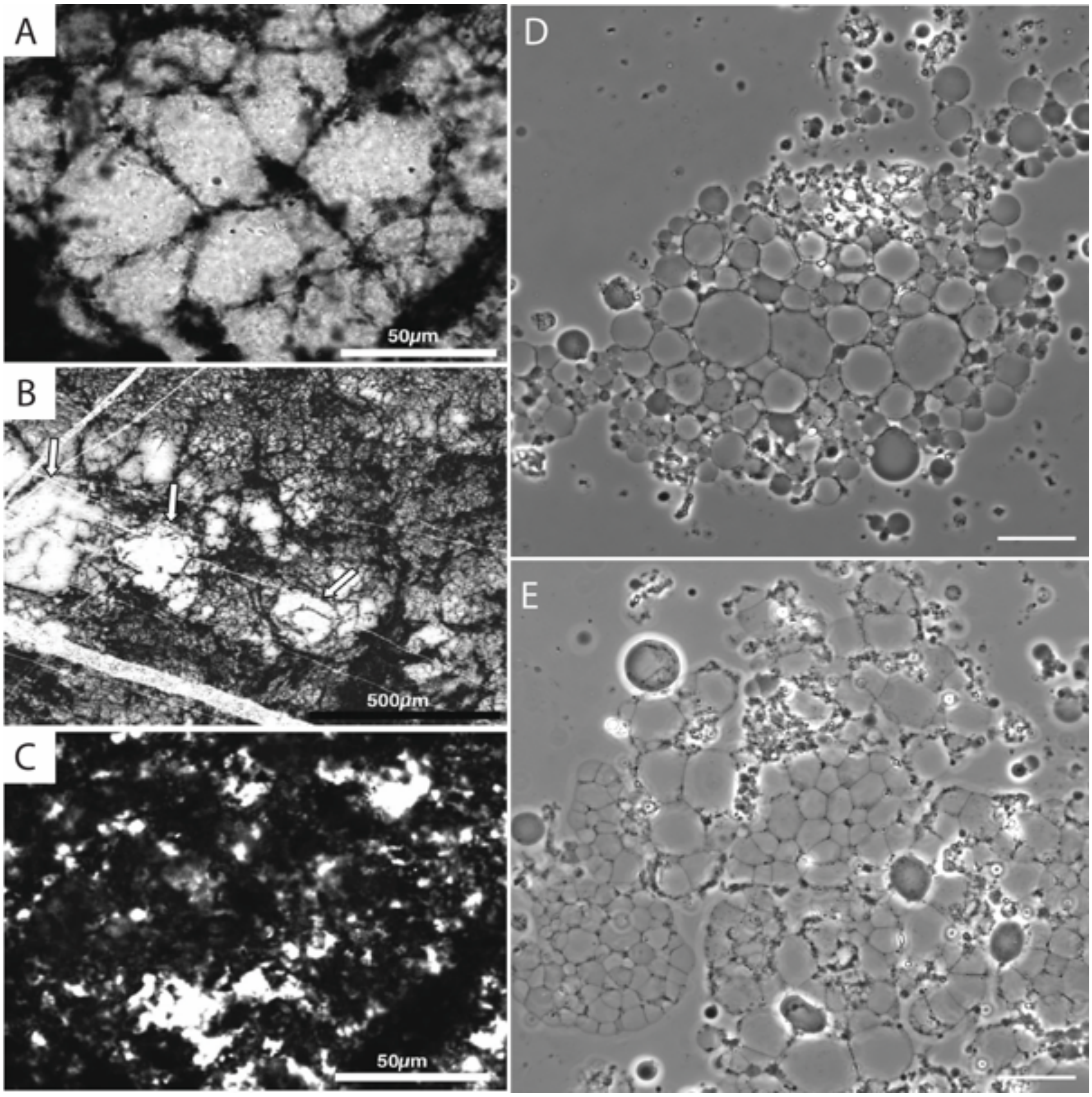


**Fig. S56: Morphological comparison of SPF microfossils with *EM-P*.** Images A-C are the organic structures reported from the SPF (originally published by Sugitani et al., 2010)(81). Images D & E show phase-contrast images of *EM-P* cells. Highlighted regions in images D & E show *EM-P* cells forming hexagonal structures, similar to SPF microfossils. Scale bar: D & E (20µm).

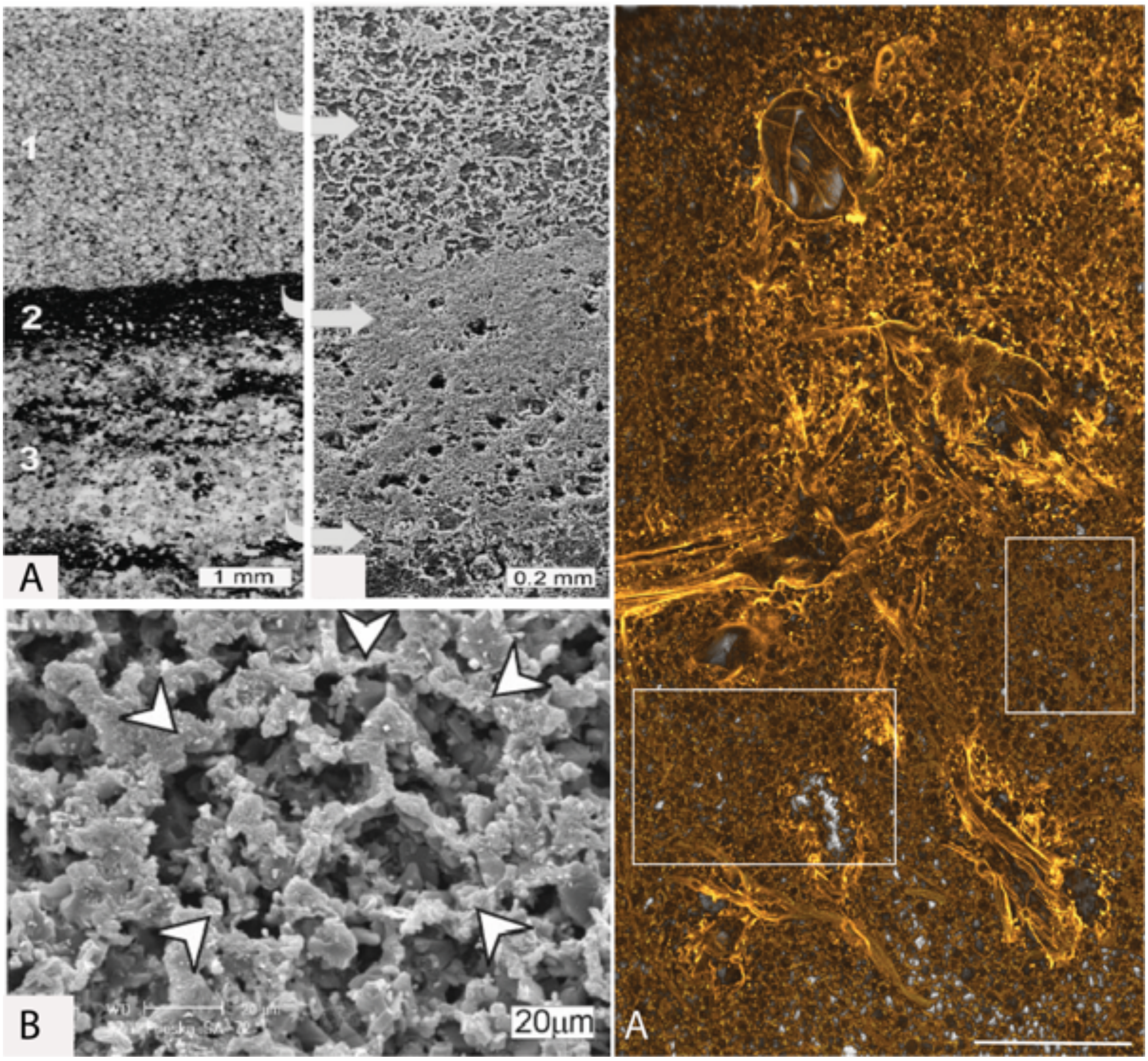


**Fig. S57. Morphological comparison between organic structures reported from the Nuga Formation and honeycomb structures of *EM-P*.** Images A & B are honeycomb-shaped organic structures reported from the Nauga Formation (originally published by Kazmierczak *et al*., 2009)(48). Arrows in A point to the magnified image. Arrows in B point to the honeycomb structures within the organic structures. Image D shows morphologically analogous structures formed by *EM-P*s. Cells in images D-F were stained with membrane stain, $FM^{TM}$5-95 (yellow). Boxed regions within Image C show polygonal structures similar in their morphology to organic structures shown in Image-b. The Scale bar: 20µm (C).

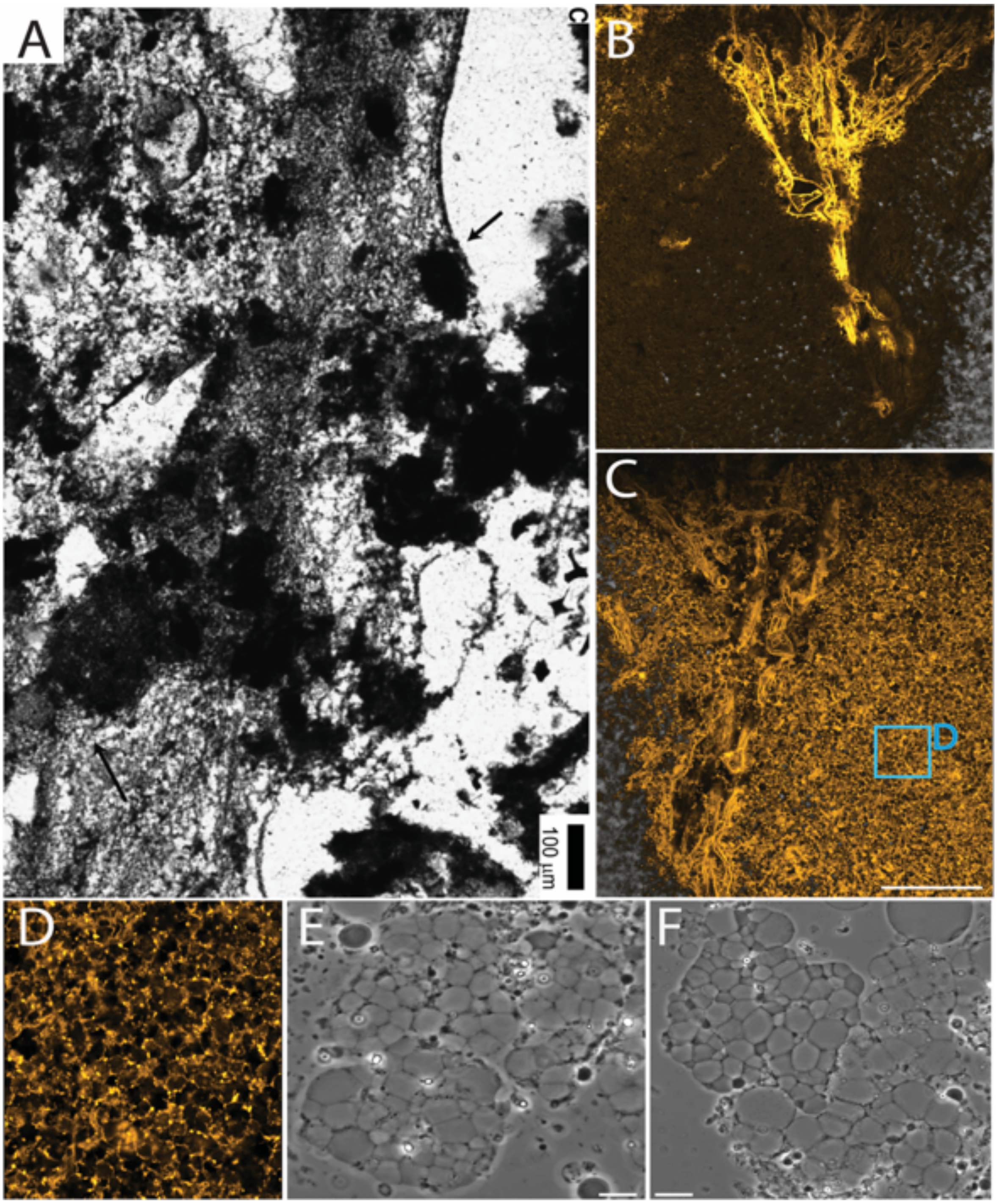


**Fig. S58. Morphological comparison between organic structures reported from the Buck Reef Chert (BRC) Formation and honeycomb structures of *EM-P*.** Image A is a honeycomb-shaped organic structure reported from the BRC (originally published by Tice *et al*., 2009)(59). Images B & C are anterior and posterior views of morphologically analogous *EM-P's* membrane debris. Cells in images B-D are stained with membrane stain, FM$^{TM}$5-95 (yellow). Image D is the magnified region of the biofilm showing honeycomb structures. Images E & F are phase-contrast images of large spherical *EM-P* cells with polygonal vacuoles undergoing clumping to form large honeycomb-shaped mats. Scale bars: B & C (100μm) and E & F (10μm).

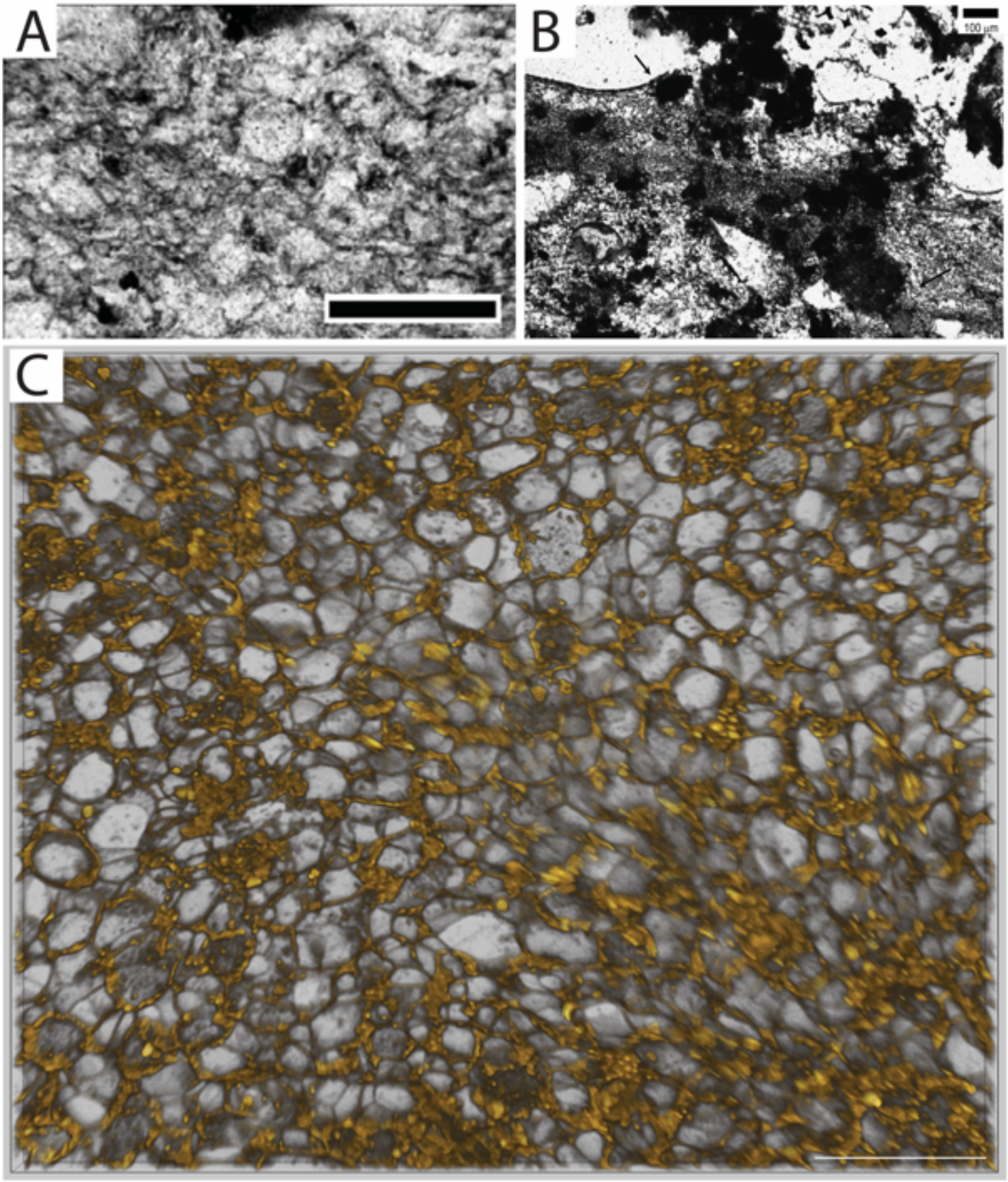


**Fig. S59. Morphological comparison between organic structures reported from the BRC Formation and honeycomb structures of *EM-P*.** Images A & B show the bifurcating or very fine strands of carbonaceous matter within the BRC formation (originally reported by Tice et al., 2006)(56). Image C is the morphologically similar structure formed by *EM-P*. The subsequent disintegration of these honeycomb structures into strands of filamentous membrane is shown in Fig. S60 B & C. Scale bars: C (20μm).

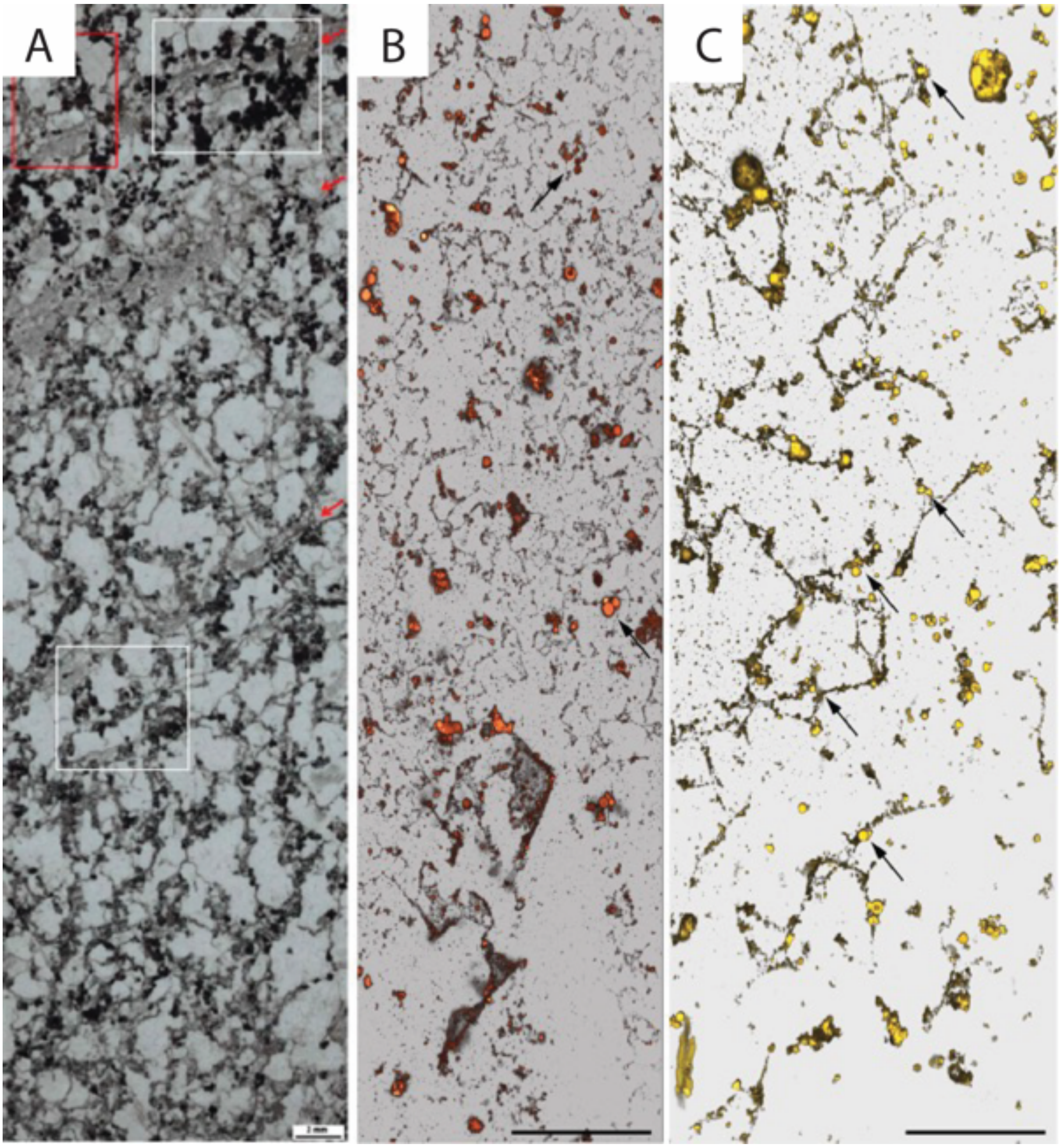


**Fig. S60. Morphological comparison of honeycomb structures reported from the Turee Creek Formations and *EM-P.*** Image A shows a tangled network of microfossil structures reported from the Turee Creek Formation (originally published by Barlow *et al*., 2018)(40). Images B & C are morphologically analogous structures formed by *EM-P*. Cells in these images are stained with membrane stain, $FM^{TM}$5-95, and imaged using a STED microscope. Boxed regions and arrows in image A highlight the spherical cells closely associated with honeycomb-shaped organic structures (see Fig. S56 & S59C). Arrows in B & C point to spherical daughter cells attached to membrane debris. Based on the morphological resemblance, we propose that the Turee Creek organic structures were leftover membrane debris of *EM-P*-like cells rather than an entangled network of filamentous microfossils. Scale bars: A (2mm), B & C (50μm).

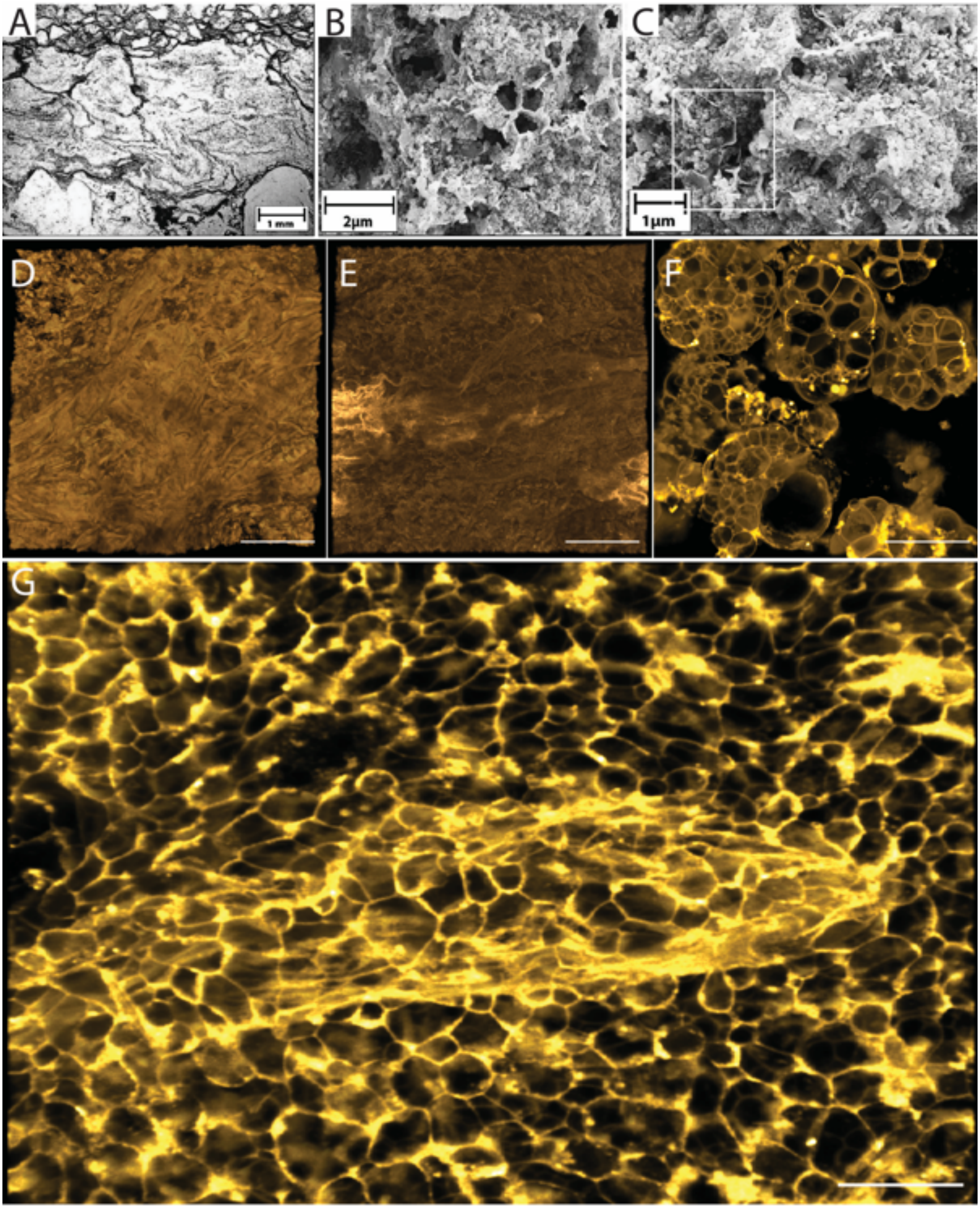


**Fig S61: Morphological comparison of organic structures reported from the Moodies Group with *EM-P*.** Images A-C are the microbial mats reported from the Moodies Group. Honeycomb-like polygonal structures (Gamper et al., 2012)(54). Images D-G are morphologically analogous structures observed in *EM-P* incubations. Images D & E show the encrusted surface and interior of the biofilm. The hexagonal structures underneath the surface can be seen in image E. Image F shows the individual *EM-P* cells that constitute the biofilm.

Image G shows the deeper layers of the biofilm with a distinctive honeycomb pattern. Scale bars: D-G (20μm).

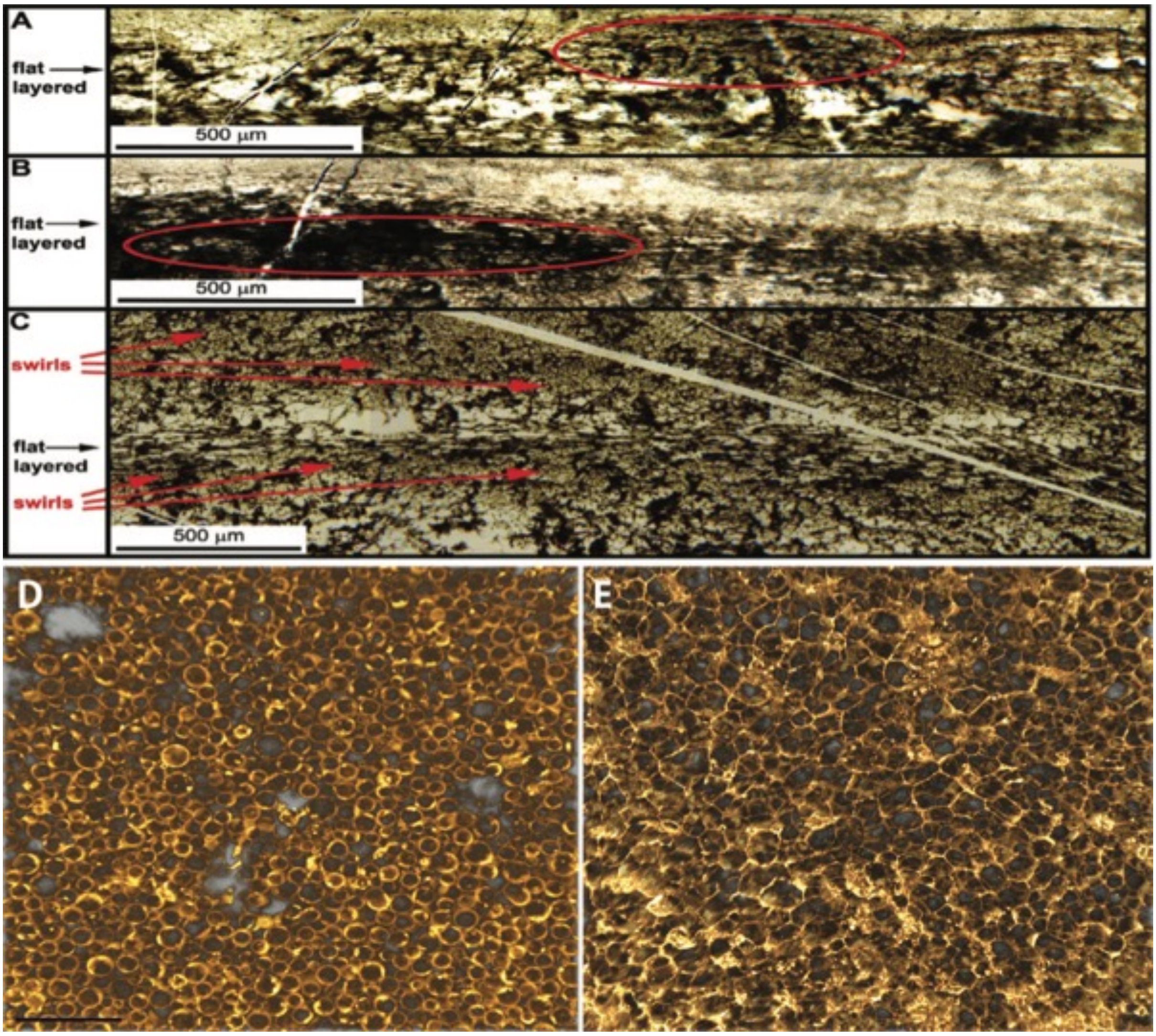


**Fig S62: Morphological comparison between honeycomb structures of *EM-P* organic structures reported from SPF.** Images A-C are honeycomb-shaped organic structures reported from SPF (originally published by Schopf *et al*., 2017)(55). The circled region in Image B points to flattened honeycomb structures. Images D & E show morphologically analogous honeycomb-like structures observed in *EM-Ps* incubations. Cells in images D-F were stained with membrane stain, FM$^{TM}$5-95. The scale bar: 20μm (D).

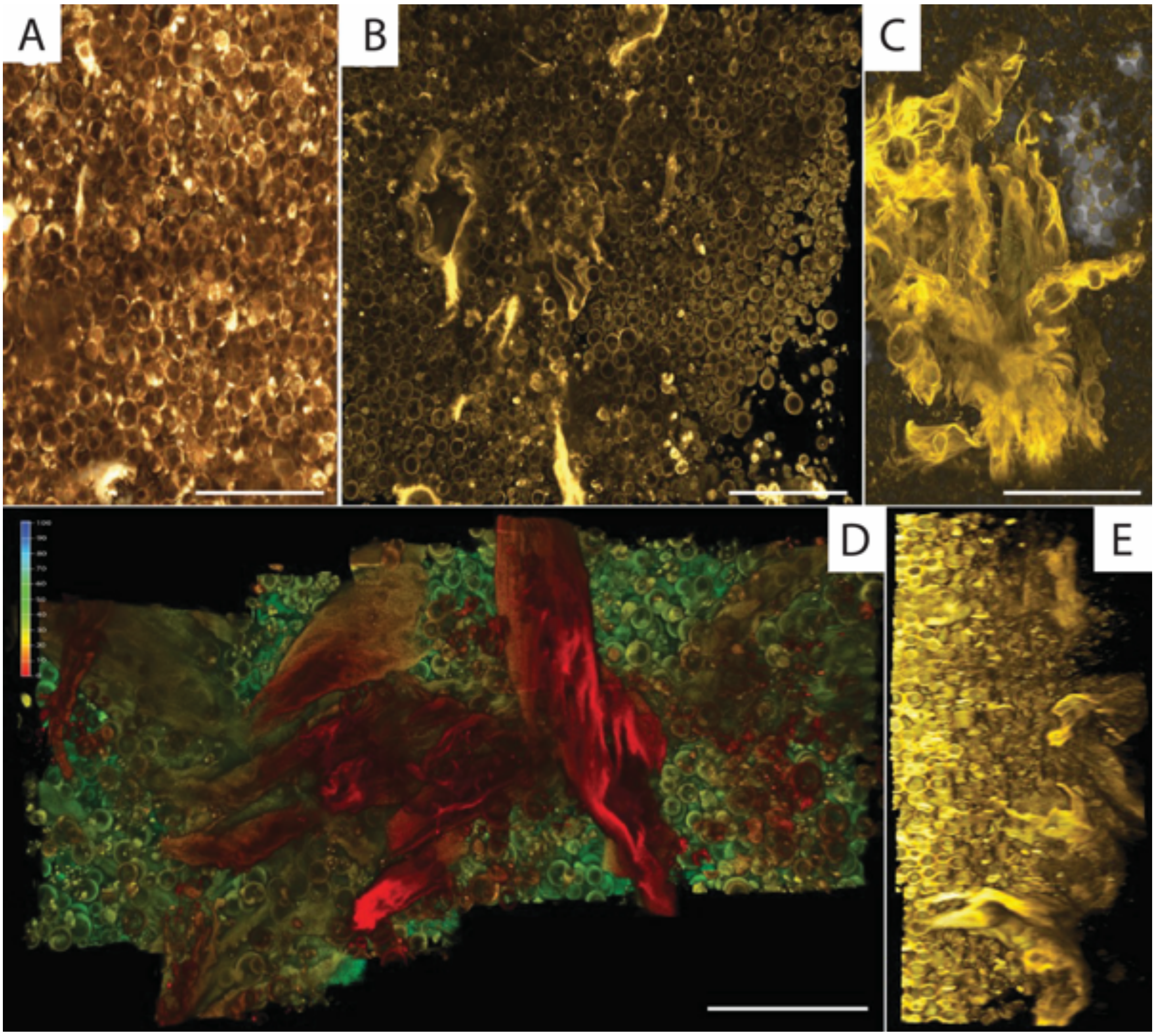


**Fig S63: Membrane debris formation.** Images A-E show different stages involved in forming fabric-like membrane debris formation. Image A shows a biofilm with stacks of spherical *EM-P* cells. Image B shows the membrane debris formed during the cell lysis. Image C shows the gradual accumulation and increase in the surface area of membrane debris. Image D shows the top view of the biofilm with sheets of membrane debris growing out of the biofilm surface. A later stage of the biofilm largely engulfed in the membrane debris is shown in Fig. S65. Image E is the lateral view of the *EM-P* biofilm shown in D. Cells and membrane debris in these images were stained with membrane stain, FM™5-95, and imaged using a STED microscope. Scale bars: 20µm.

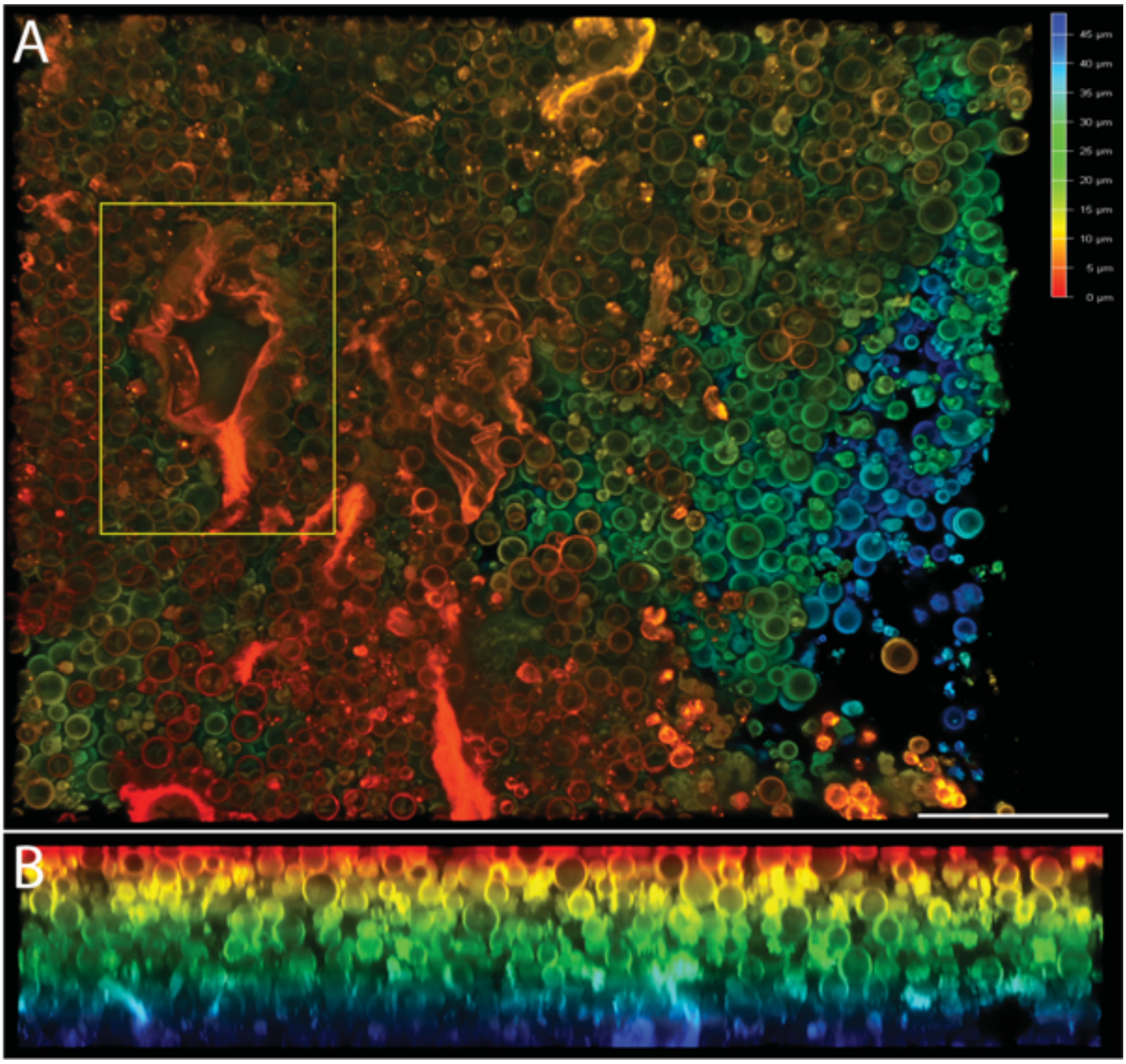


**Fig. S64. Cell lysis and formation of membrane debris within multilayered *EM-P* cells.** Images A & B show a top and lateral view of *EM-P* cells growing in multiple layers at the bottom of the chamber slide. Cells in the above images were stained with membrane stain, $FM^{TM}$5-95. The boxed region in image A shows the membrane debris formed from the lysis of cells. Scale bar: 20µm (A).

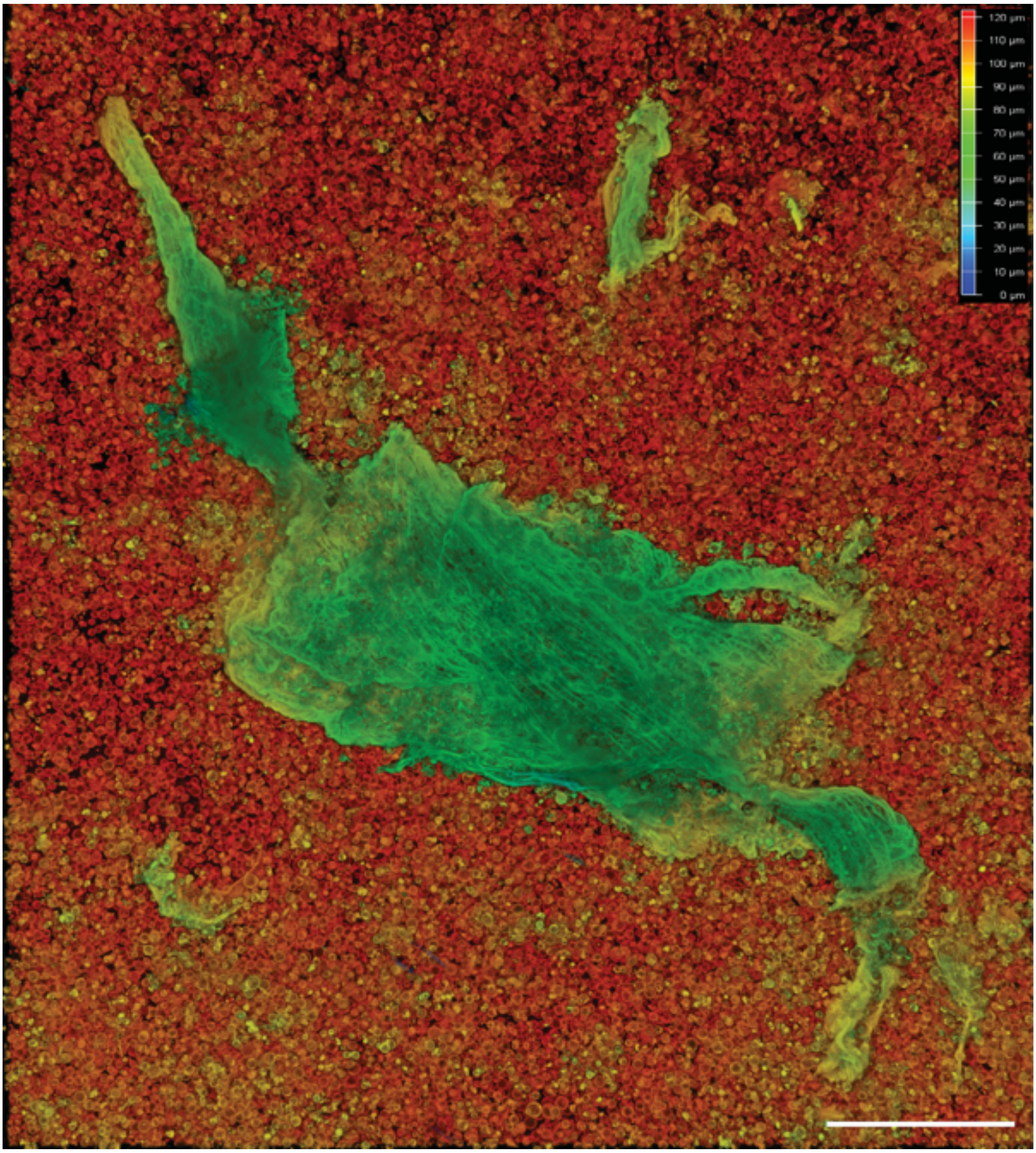

**Fig. S65. Membrane debris of *EM-P*.** The image shows membrane debris of *EM-P* forming a layer over a mat of multilayered cells. Based on the morphological similarities with laminated structures, we presume such structures were formed by a similar process. The lateral view of the image, along with its morphological comparison with α-type laminations reported from BRC, was shown in Fig. S66. Membranes were stained with Nile red and were imaged using a confocal microscope. Scale bar: 100μm.

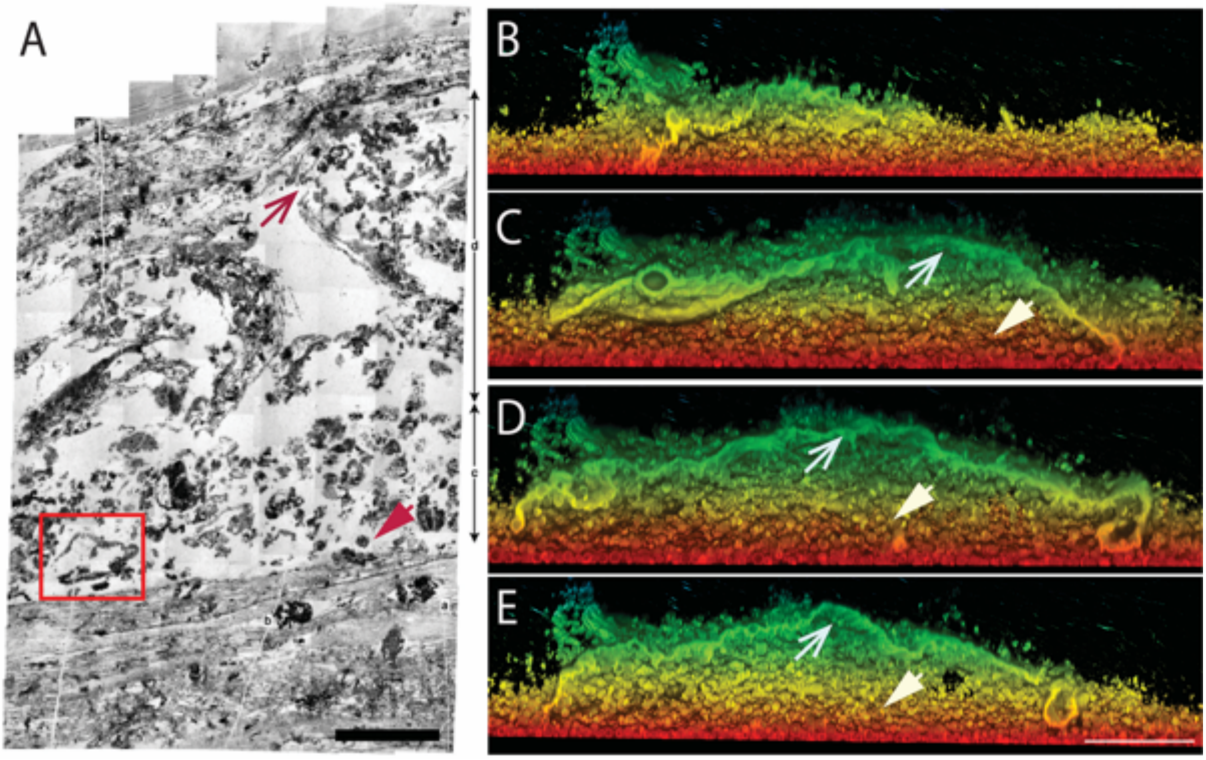


**Fig. S66. Morphological comparison between the BRC laminations and membrane debris of *EM-P.*** Image A shows α-type laminations reported from the BRC (originally published by Tice *et al.,* 2009)(59). Two filamentous layers with hollow spaces between them can be seen in image A. The hollow space in between is filled with filamentous membrane debris and spherical inclusions. Images B-E show lateral sections of *EM-P* membrane debris, as shown in Fig. S65. Open and closed arrows in all the images point to the top layer of membrane enclosure and the bottom cell layer, respectively. The hollow space in between is filled with membrane debris and spherical cells. We presume that the lysis of these cells could have produced membrane debris like the ones observed in image A. In support of this presumption, membrane debris similar to β-type laminations (indicated by the red box) was observed in *EM-P* batch cultures (shown in Fig. S71 & S74). Also see Movie S19. Scale bars: 5mm (A) and 100μm (E).

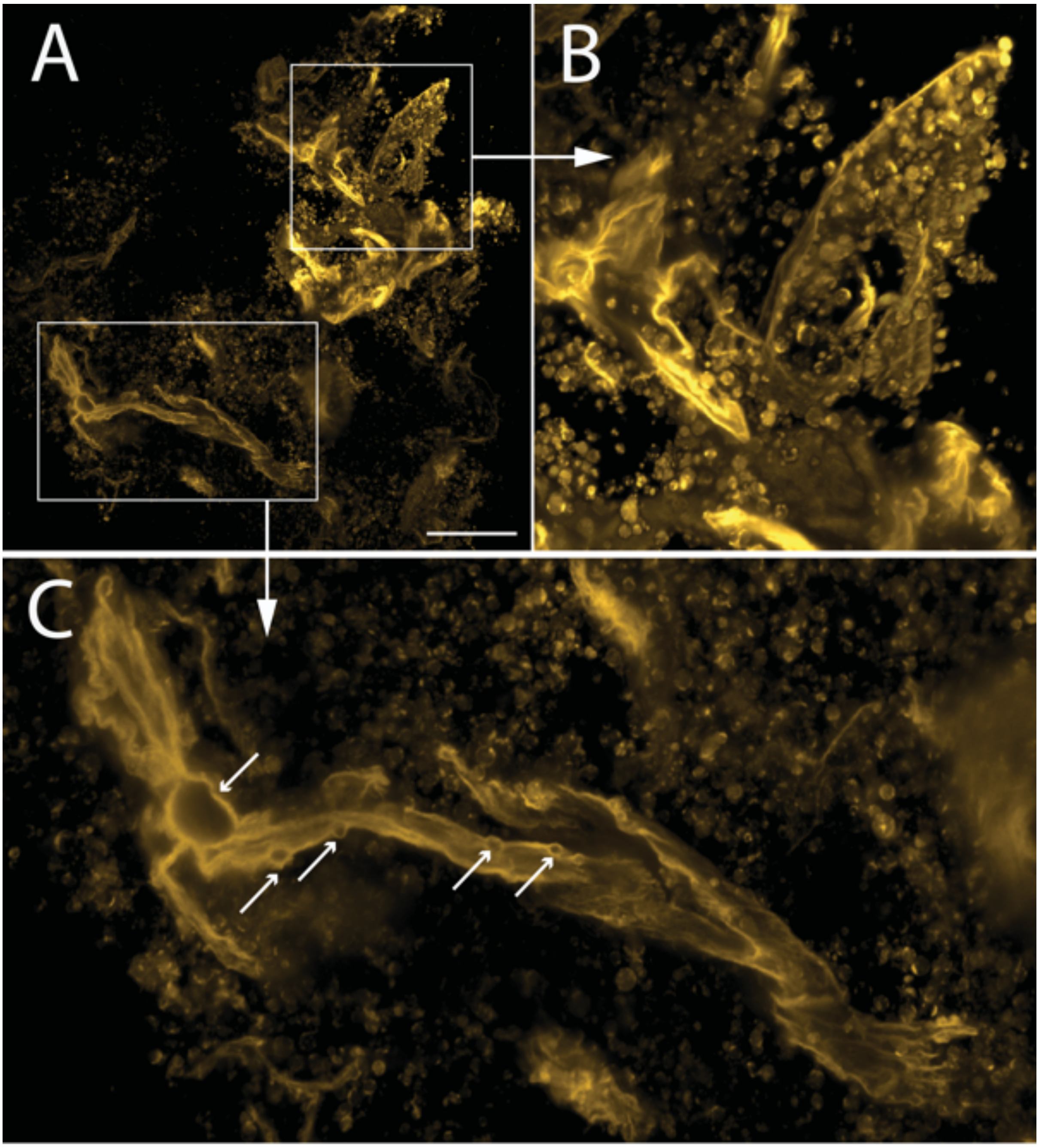


**Fig. S67. Membrane debris of *EM-P*.** The image shows *EM-P* cells covered in membrane debris. Images B & C are the magnified regions of image A. Arrows in c point to the naturally formed lenticular gaps within the membrane debris. Scale bar: 100µm (A).

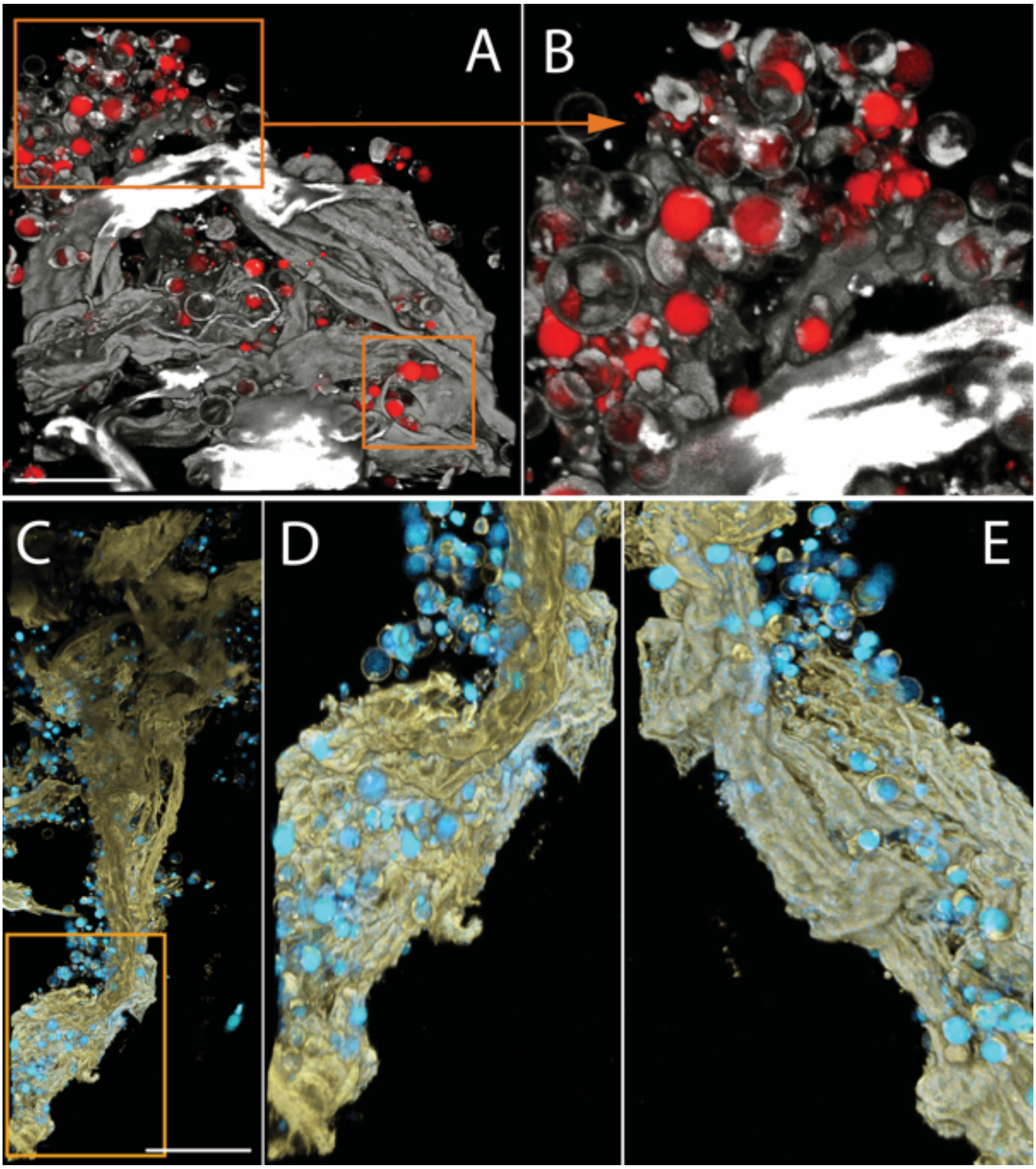


**Fig. S68. Membrane debris of *EM-P*.** The image shows *EM-P* covered in membrane debris. Image B shows the magnified regions of image A. Images D & E are the anterior and the posterior view of the membrane debris with cells. These images highlight the fabric-like texture of the membrane debris. Cells in these images were stained with FM$^{TM}$5-95 (membrane, white in A & B and grey in C-E) and PicoGreen$^{TM}$ (DNA, red in A & B and cyan in C-E). Scale bar: 50µm (A & C).

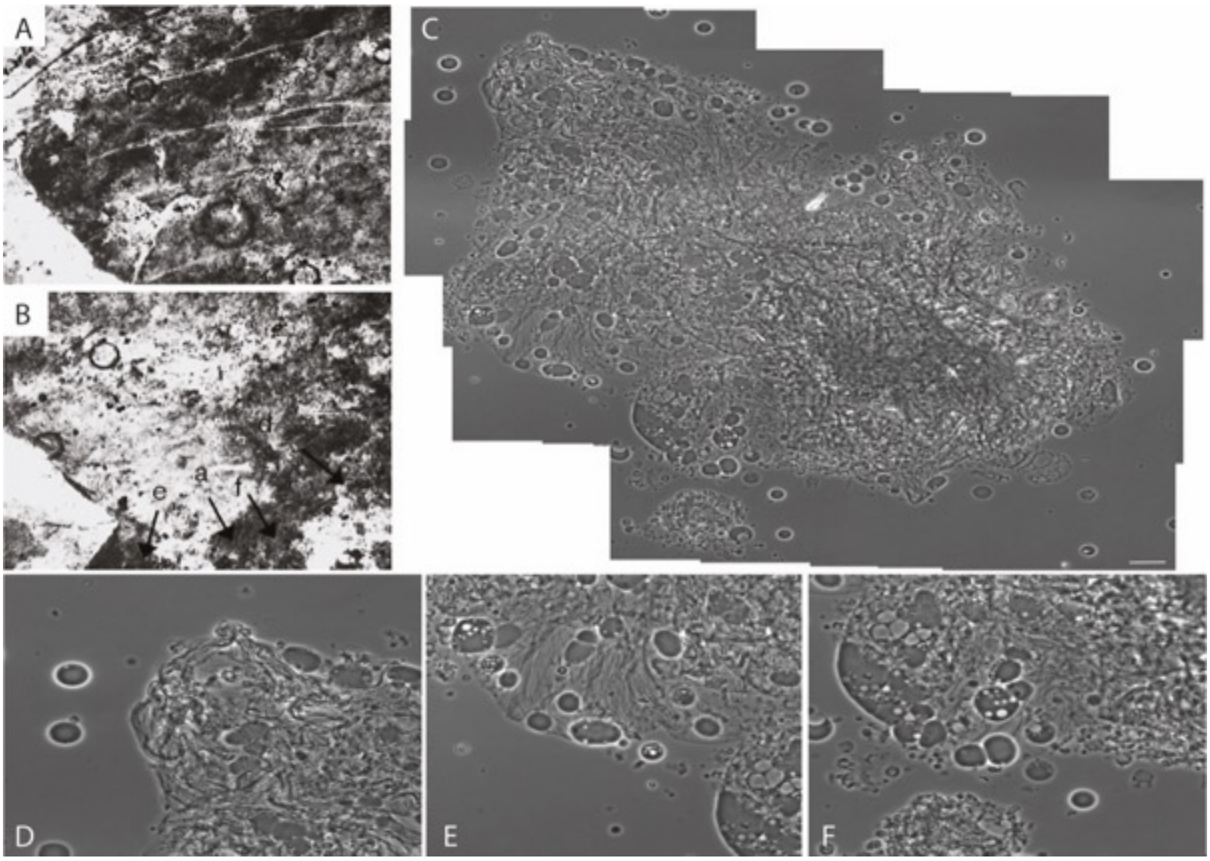


**Fig. S69. Morphological comparison between *EM-P* and the Chinaman Creek microfossils.** Images A & B are organic structures reported from the Chinaman Creek (Brasier *et al.,* 2005)(60). Image C is a composite image of morphologically analogous *EM-P* structures. Images D-F are magnified regions of image C. All the images show spherical structures with or devoid of organic carbon enclosed in a film-like structure. Scale bar: 10µm (C).

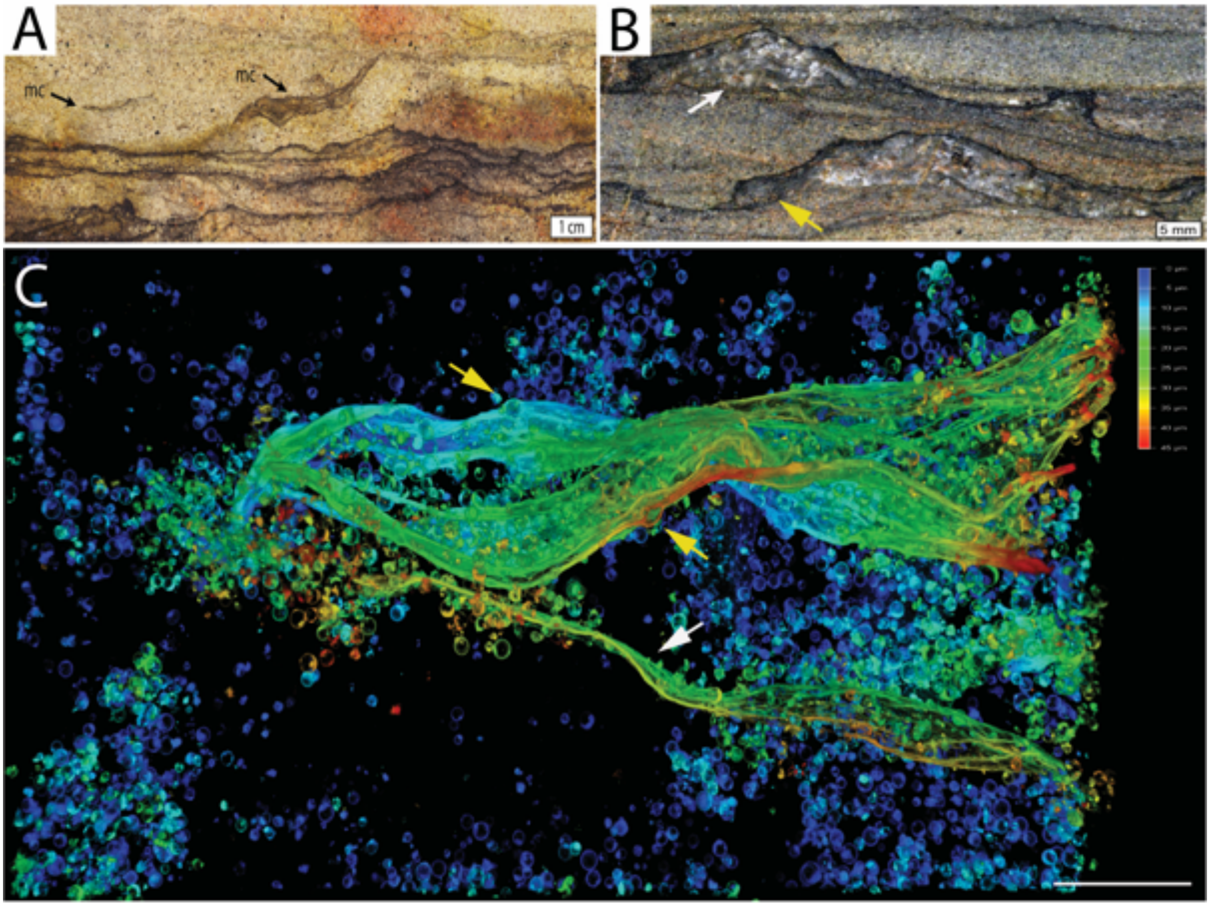


**Fig. S70. Morphological comparison of *EM-P* Membrane debris with laminated structures.** Images A & B show bifurcating laminated structures reported from the Moodies Group (originally published by Homann *et al.,* 2015)(61). Image C is a 3D-rendered confocal image of image of morphological analogous membrane debris formed by *EM-P* cells. White arrows in B point to hollow lenticular regions within the laminations, similar to the one observed in *EM-P*. Arrows in images A & C point to the bifurcation of membrane debris. Membranes were stained with Nile red, and imaging was done using a point scanning microscope. Scale bar: 50µm (C).

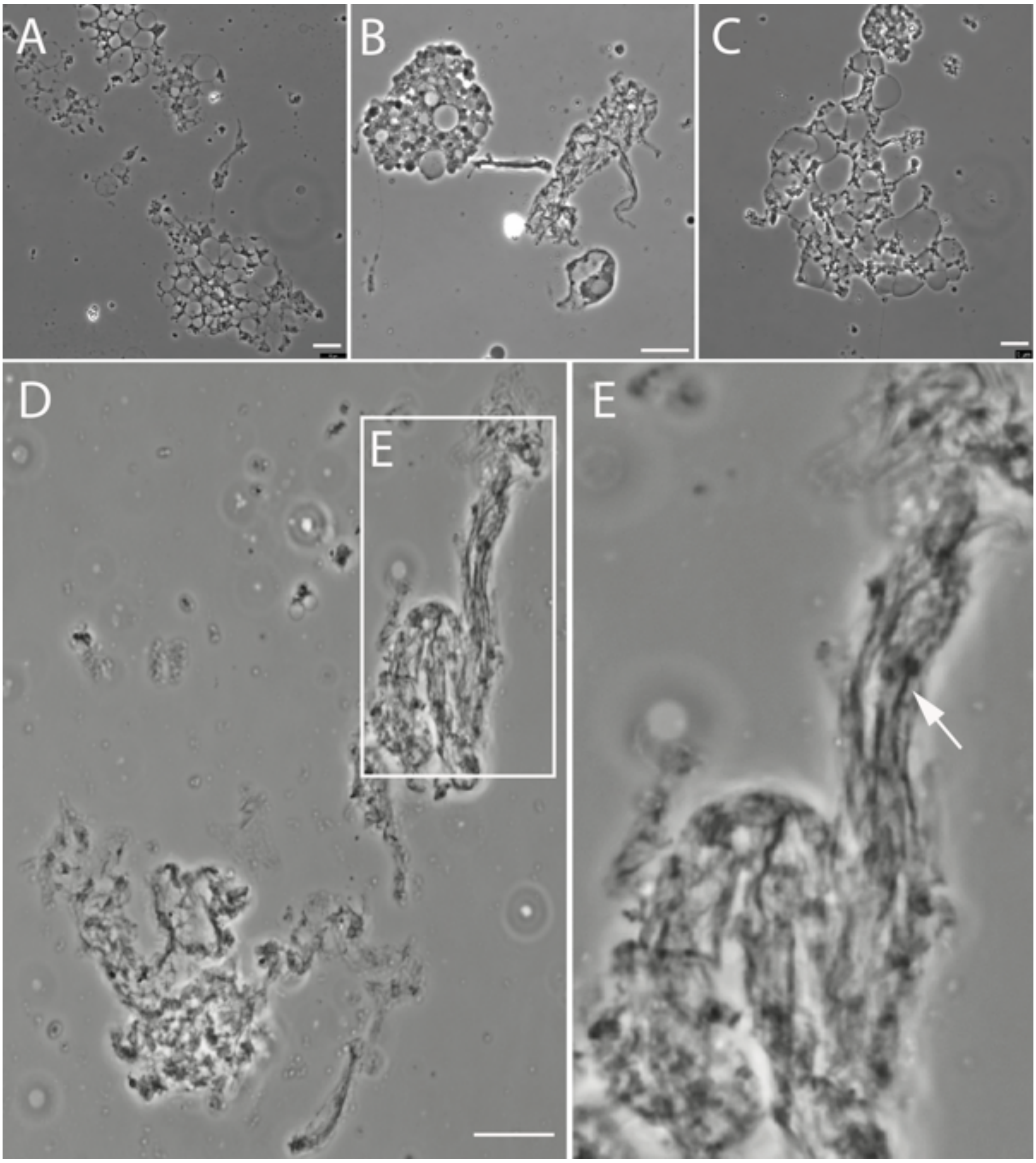


**Fig. S71. Sequential stages involved in the formation of β-laminations.** Images A-E shows the steps involved in the sequential transformation of membrane debris to β-type laminations. Image A-C shows the aggregation of vacuoles formed from the lysis of large spherical *EM-P* cells. Image D shows collapsed membrane debris formed by lateral compression or deflation of such aggregations after the release of daughter cells. The arrow in image E points to the individual membrane layers within the debris. Scale bars:10µm.

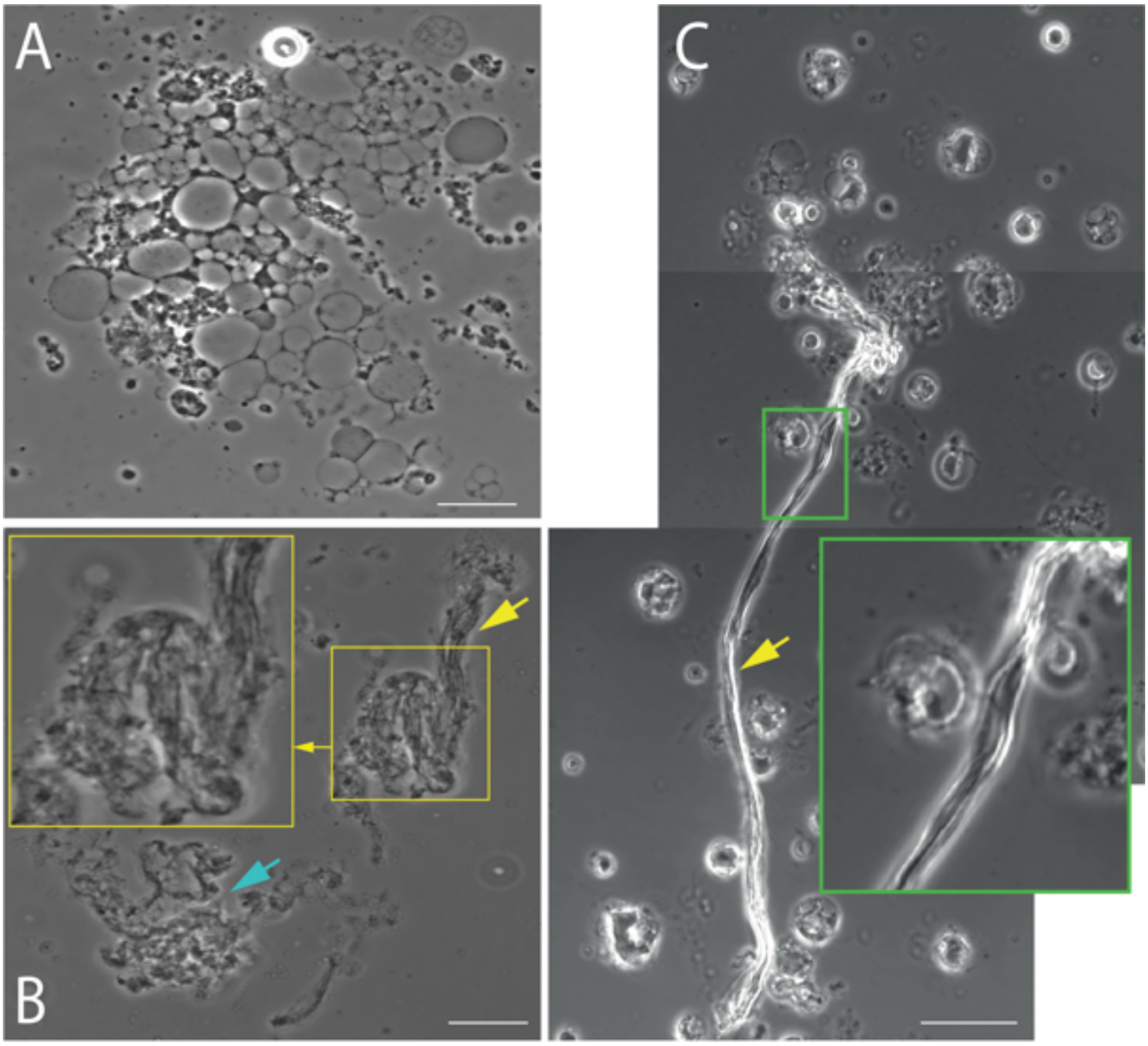


**Fig. S72. Sequential stages involved in the formation of β-laminations.** Images A-C show the steps involved in the sequential transformation of membrane debris to β-type laminations. Image A shows the aggregation of vacuoles formed from the lysis of large spherical *EM-P* cells. Image B shows collapsed membrane debris formed by lateral compression or deflation of such aggregations after the release of daughter cells. Cyan and yellow arrows point to membrane debris and compressed membrane debris, respectively. Inset in image B is the magnified region of compressed membrane debris. Image-c shows structures similar to β-type laminations. Scale bars:10µm.

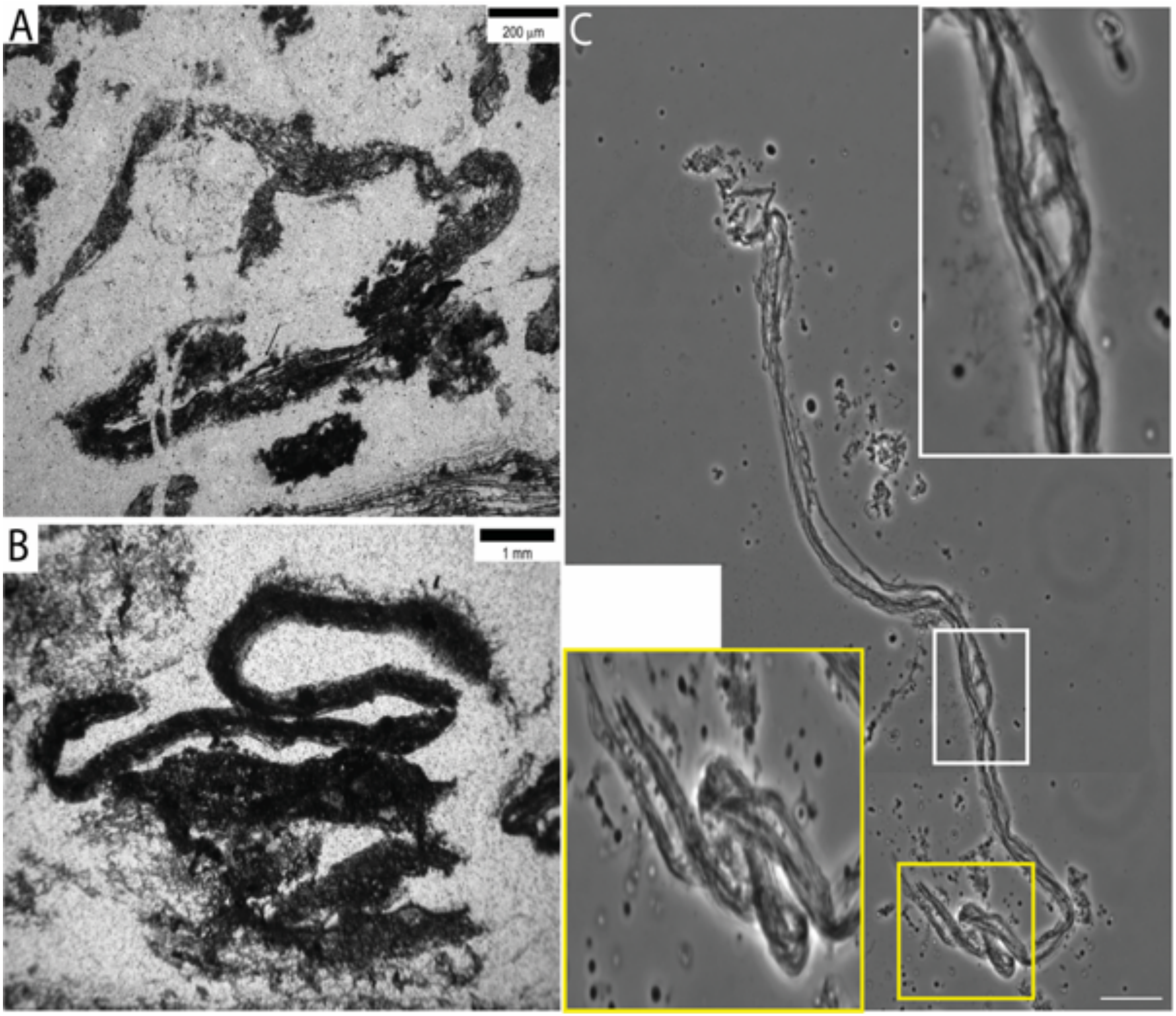


**Fig. S73. Morphological comparison of the BRC β-laminations with *EM-P's* membrane debris.** Images A & B show β-type laminations reported from the BRC (originally published by Tice *et al.,* 2004)(58). These laminations are described as rolled-up or bundled filamentous structures. Image C shows morphologically analogous membrane debris formed by *EM-P*. Insets in image C are magnified regions of the debris showing bundled-up individual filamentous structures. Scale bars: 10μm (C).

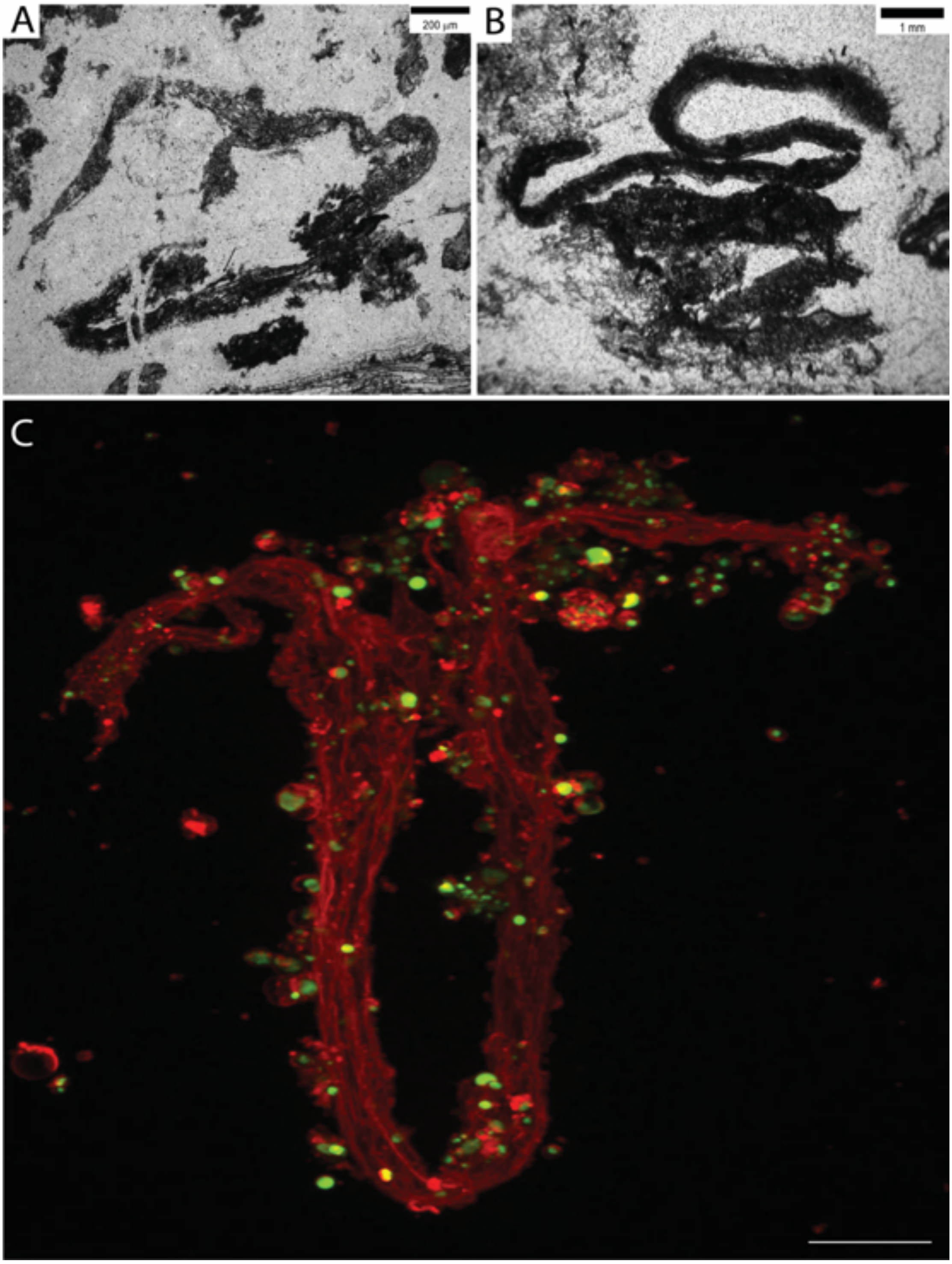


**Fig. S74. Morphological comparison of the BRC β-laminations with *EM-P's* membrane debris.** Images A & B show β-type laminations reported from the BRC (originally published by Tice *et al.,* 2004)(58). Image C shows a STED image of morphologically analogous membrane debris formed by *EM-P*. Spherical daughter cells (evidenced by the presence of DNA) are still attached to the membrane debris, which can be seen in the image. In the image, the membrane was stained with FM5-95 (red), and DNA was stained with PicoGreen (green). Scale bar: 10μm (C).

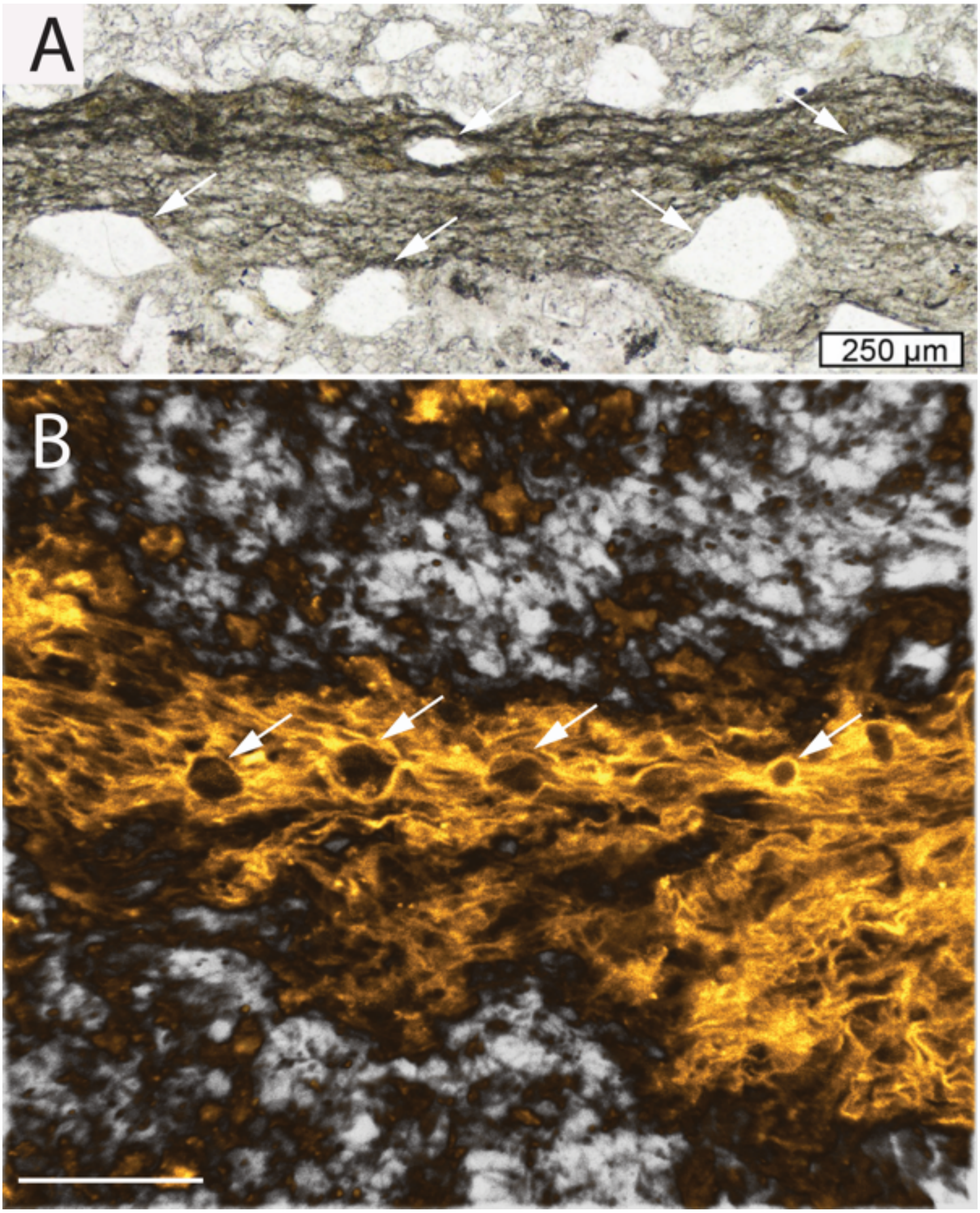


**Fig. S75. Morphological comparison of the Moodies Group laminations with *EM-P's* membrane debris.** Image A shows α-type laminations reported from the Moodies Group (originally published by Homann *et al.,* 2018)(49). Image B is a 3D-rendered confocal image of *EM-P*'s cell debris. Arrows in images A & B point to the lenticular gaps within the membrane debris. Scale bar: 20µm (B).

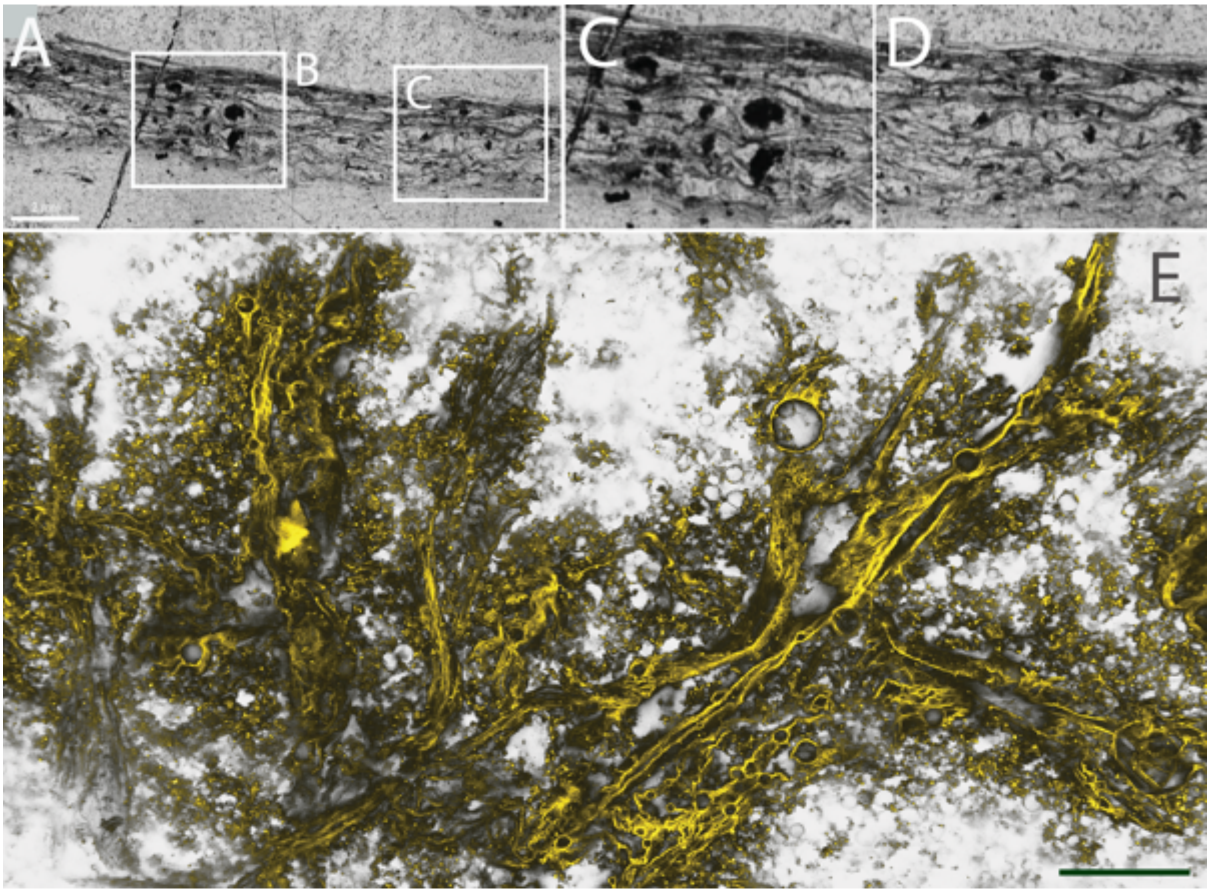


**Fig. 76: Morphological comparison between laminated structures reported from the Moodies Group and structures formed by *EM-P.*** Images A-C are laminated structures reported from the Moodies Group (originally published by Hickman-Lewis *et al.,* 2021)(4). They show filamentous structures with lenticular gaps. Image C is a 3D-rendered confocal image of *EM-P*'s membrane debris. Filamentous membrane debris bifurcating, forming spherical/lenticular gaps, can be seen in several regions. Some spherical\lenticular gaps were hollow, and some had a honeycomb pattern within them, indicating the presence of large spherical *EM-P* cells with intracellular vesicles (Fig. 8, S77 & S78). Membranes were stained with Nile red, and imaging was done using a STED microscope. Scale bar is 50µm.

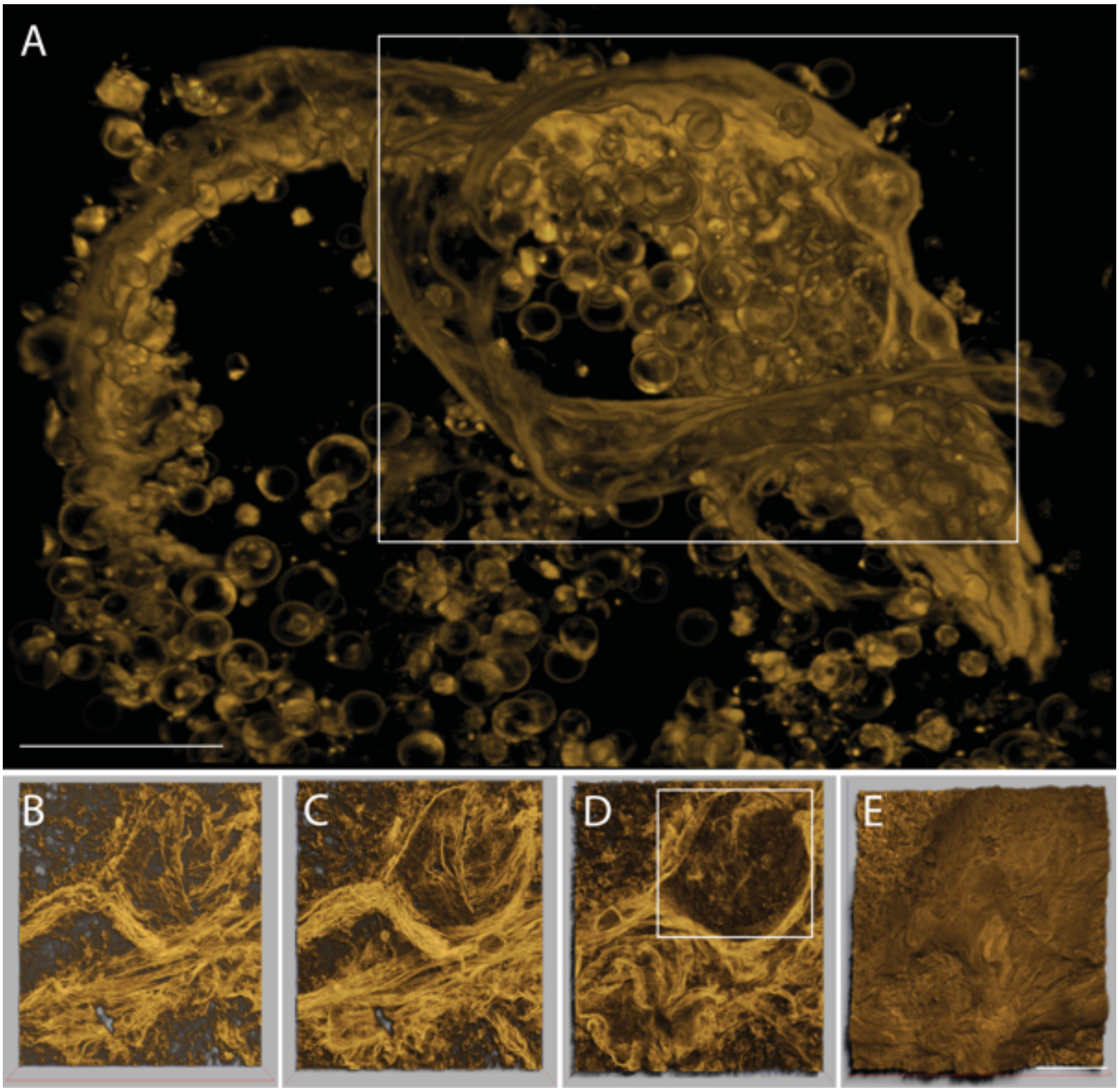


**Fig. S77. Lenticular membrane debris of *EM-P*.** Image A shows spherical *EM-P* cells covered in fabric-like membrane debris. Images B-E show an *EM-P* biofilm at different focal planes (bottom to top). The boxed region in image D shows the honeycomb pattern within the lenticular gap (also see Fig. S78 & Movie 21). Scale bar: 20µm.

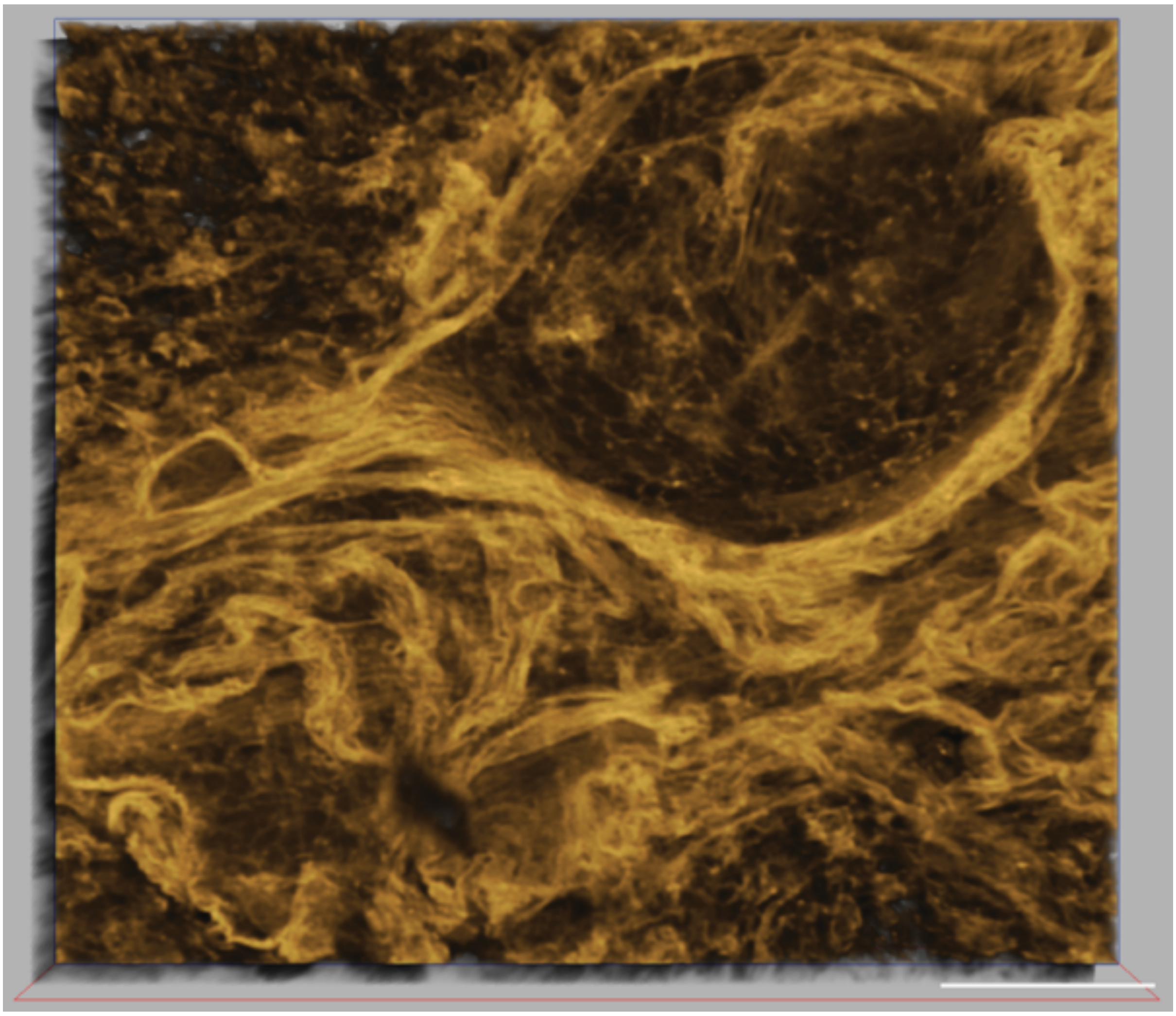

**Fig. S78. Lenticular membrane debris of *EM-P*.** The image shows the membrane debris produced by the lysis of *EM-P* cells. A honeycomb pattern within the lenticular structure suggests the presence of individual cells or *EM-P*s vesicles during the time of their formation. (also see Movie 21). Scale bar: 20µm.

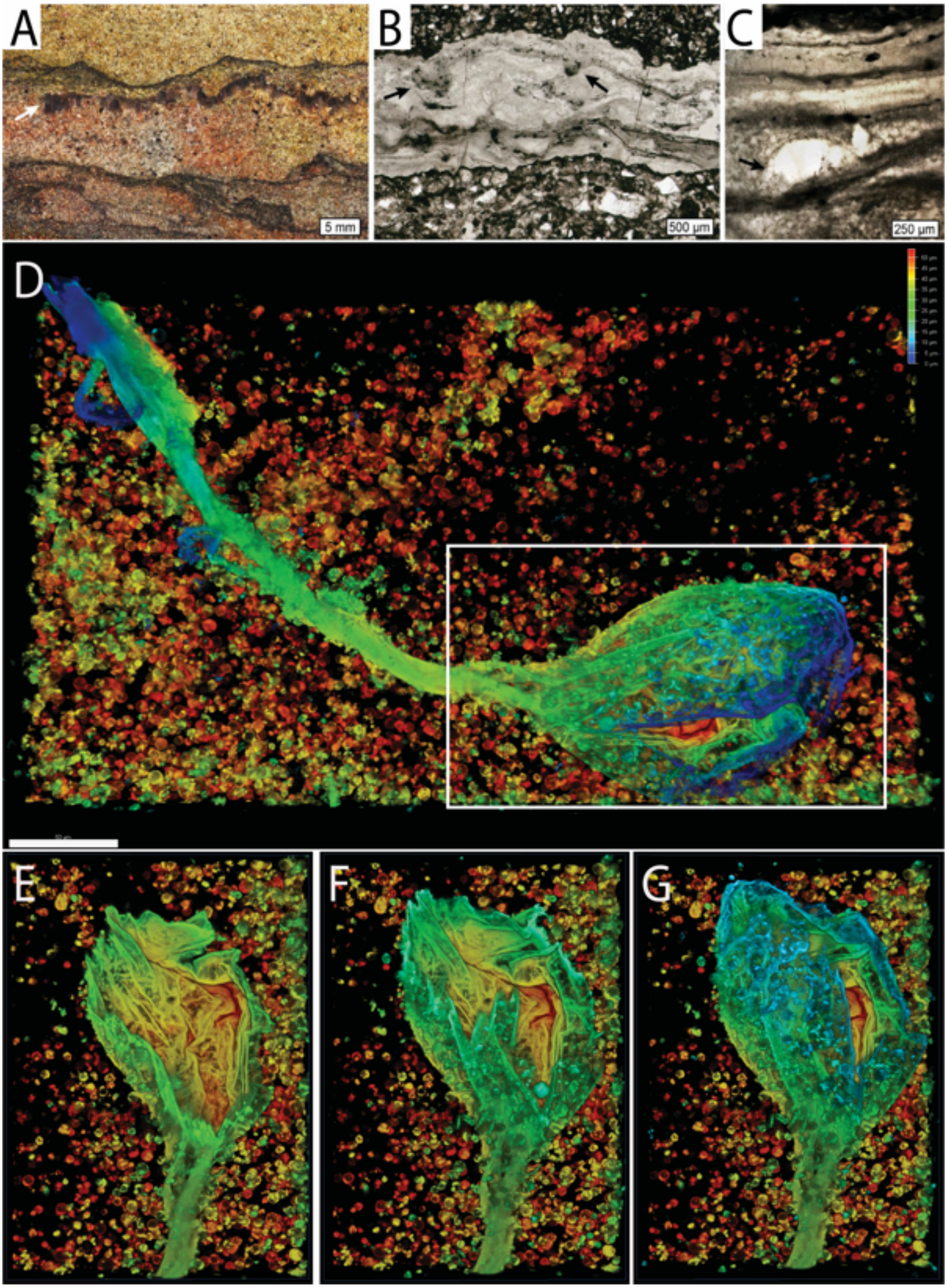


**Fig. S79. Morphological comparison of the Moodies Group laminations with *EM-P's* membrane debris.** Image A-C shows laminations reported from the Moodies Group (originally published by Homann *et al.,* 2018)(49). Image D is a 3D-rendered confocal image of *EM-P*'s membrane debris exhibiting hollow lenticular gaps (Movie 20). Images e-g show the hollow lenticular structure at different Z-axis positions. Membranes were stained with Nile red, and imaging was done using a STED microscope. Scale bar: 50µm.

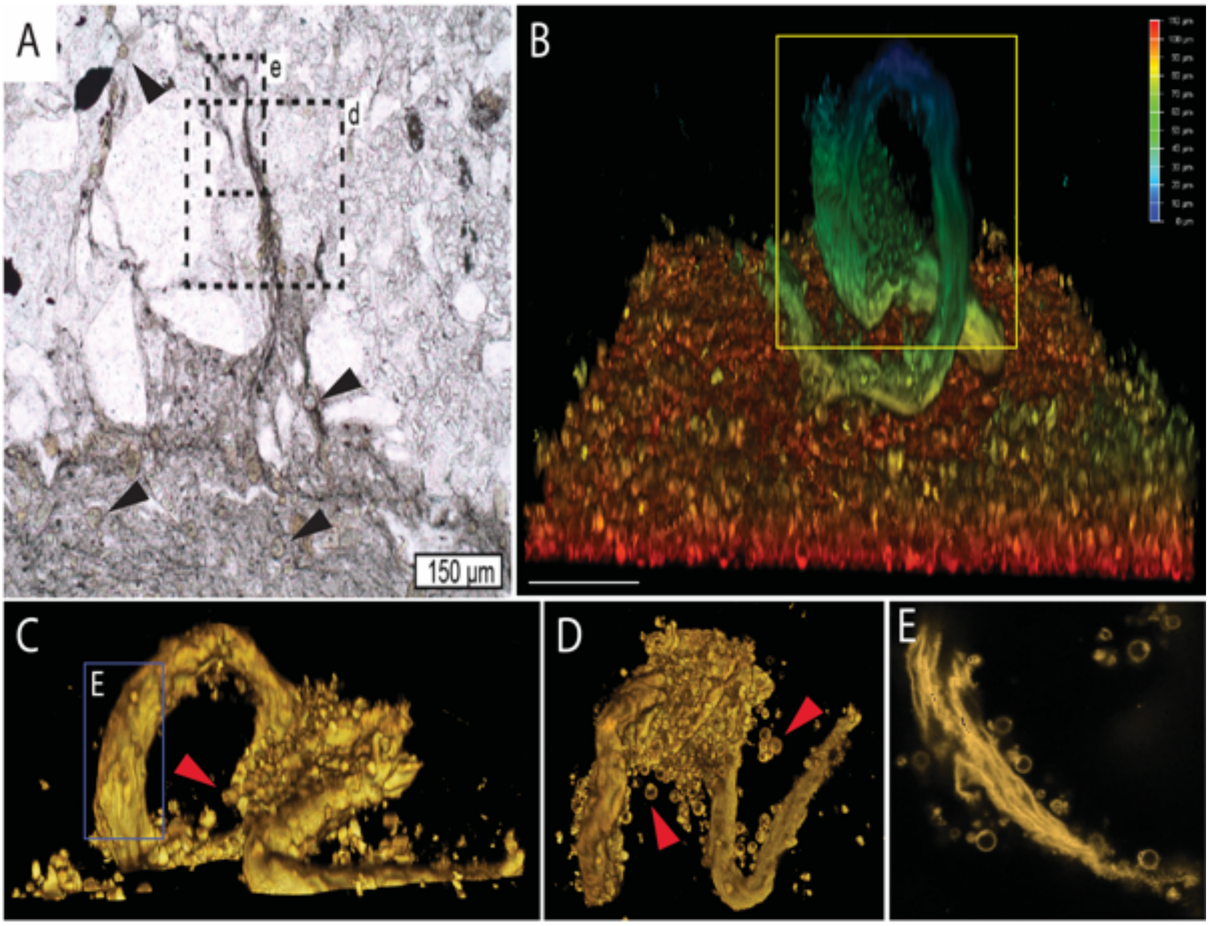


**Fig. S80. Morphological comparison of the Moodies Group laminations with *EM-P's* membrane debris.** Image A show β-type laminations reported from the Moodies Group (originally published by Homann *et al.,* 2018)(49). Laminations from the Moodies Group were shown to form raised filamentous structures (A). Arrows in the image point to spherical structures within the laminations. Images B-D show 3D-rendered confocal images of morphologically analogous *EM-P's* membrane debris. Image B shows filamentous membrane debris of *EM-P*, that rose above layers of spherical cells. Images C & D are close-up images of raised filamentous membrane debris of *EM-P* from different viewing angles. Arrows in these images point to spherical cells attached to the membrane debris. Image E is a non-3D rendered image showing individual membrane layers within the debris, with spherical inclusions attached. Scale bar: 20µm (B).

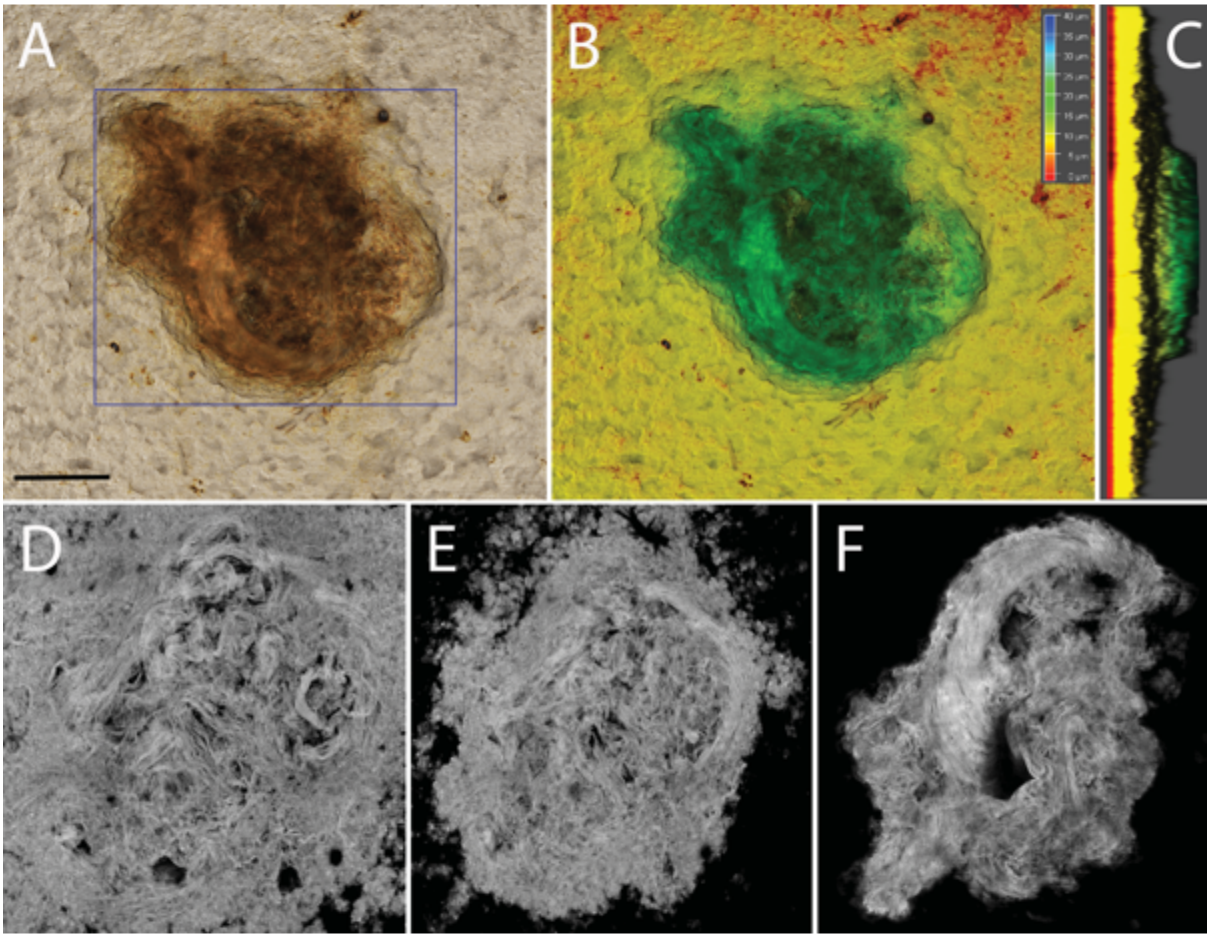


**Fig. S81. Swirl-like structures of *EM-P*'s membrane debris.** Images A-F show a 3D-rendered STED image of membrane debris observed in late-growth stages of *EM-P* (1–2-month-old). This debris typically formed raised mound-like structures. Images A & B show the top view of such structures, and image C shows the side view. Images d-f are images of such mounds at different depths (bottom to top). The formation of swirls can be seen in images E & F. In the image, membrane debris is stained with Nile red. A morphological comparison of these structures with Archaean laminations is shown in Fig. S82. Scale bar: 50μm (A).

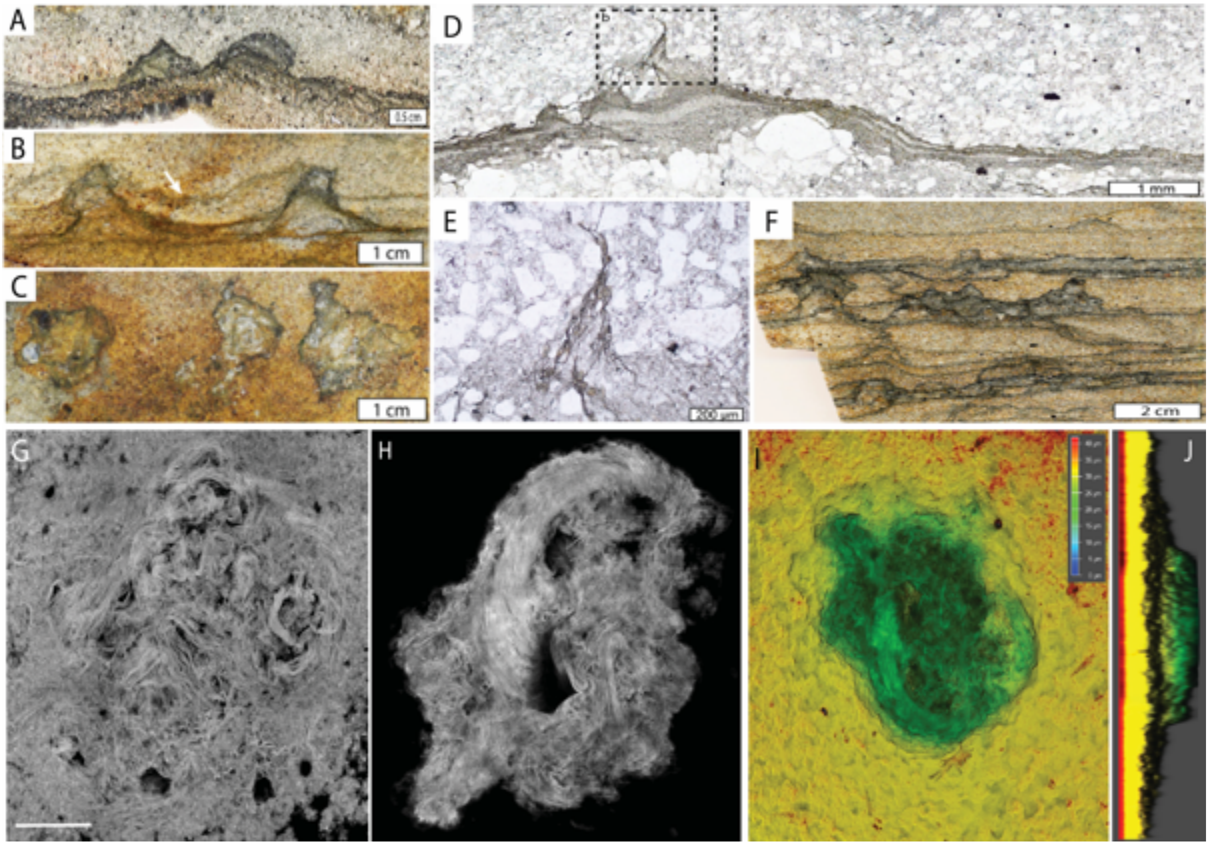


**Fig. S82. Morphological comparison of the Moodies Group laminations with *EM-P's* membrane debris.** Images A-F shows raised mound-like structures reported from the Moodies Group (Homann *et al.,* 2018)(49). Image C shows the top view of such mounds. These mounds were shown to have filamentous extensions at their peaks. Images G-J are morphologically analogous structures formed by *EM-P*. In these images, the membrane debris is stained with Nile red. Images I & J show the top and side view of these raised mounds (depth coloring). Scale bar: 50µm (G).

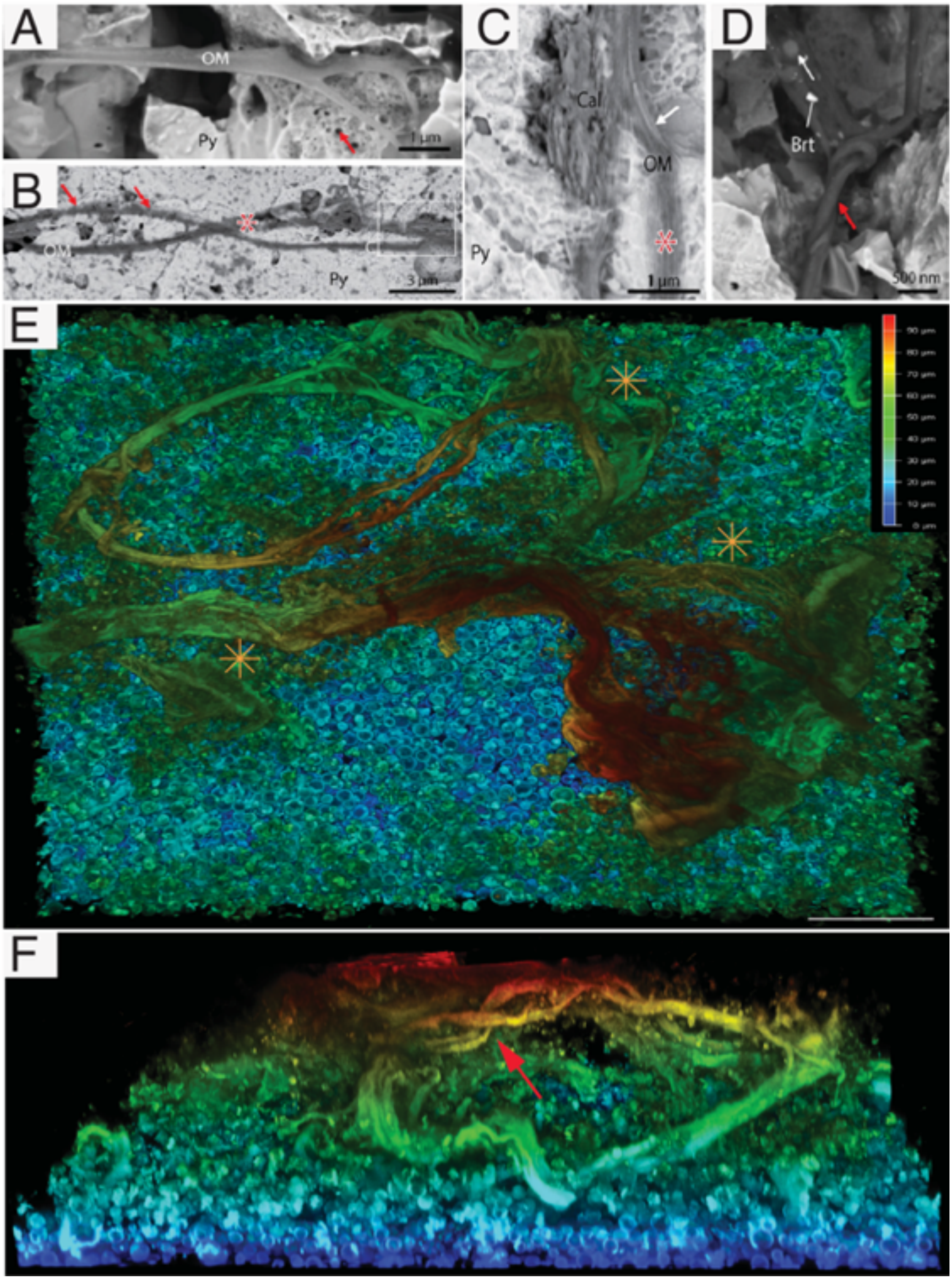


**Fig. S83. Morphological comparison of laminations reported from the BRC with *EM-P's* membrane debris.** Images A & B show β-type laminations reported from the BRC (originally published by Tice *et al.,* 2009)(59). Images E & F are frontal and lateral view of morphologically analogous structures formed by *EM-P*. Arrows in these images point to helical structures observed in microfossils and *EM-P*. Star signs in images B, C, and E point to regions showing the filamentous nature of the membrane debris. In the image, the membrane was stained with Nile red. Scale bars: 50µm (E).

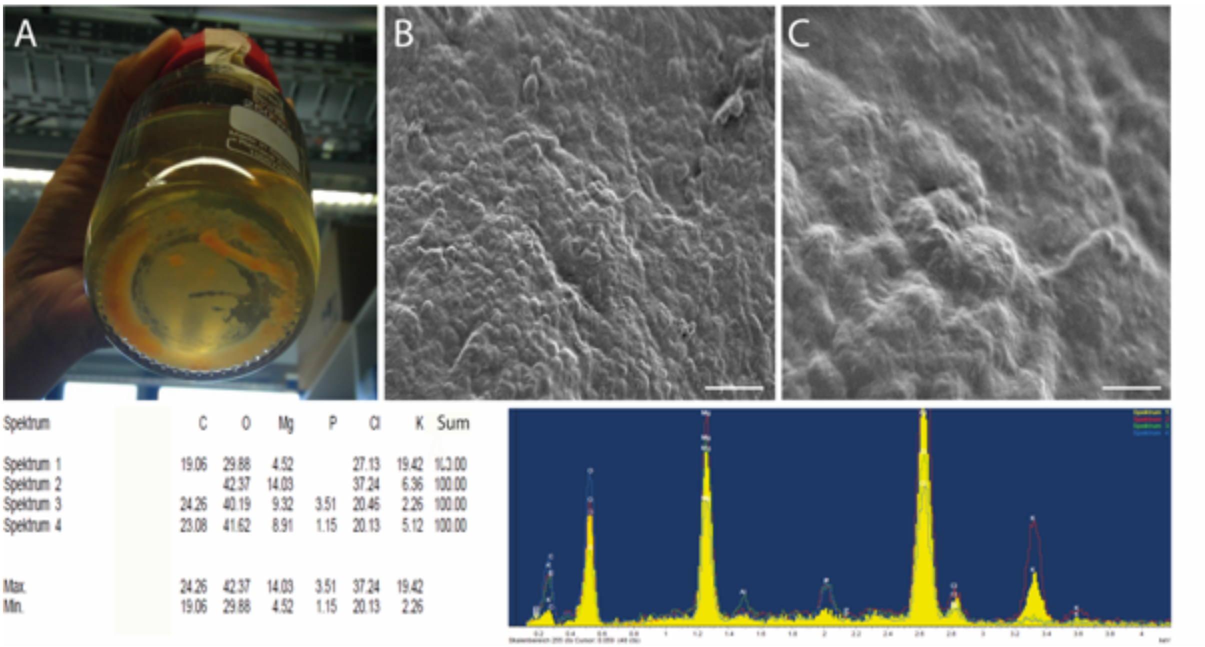


| Spektrum | C | O | Mg | P | Cl | K | Sum |
|---|---|---|---|---|---|---|---|
| Spektrum 1 | 19.06 | 29.88 | 4.52 | | 27.13 | 19.42 | 100.00 |
| Spektrum 2 | | 42.37 | 14.03 | | 37.24 | 6.36 | 100.00 |
| Spektrum 3 | 24.26 | 40.19 | 9.32 | 3.51 | 20.46 | 2.26 | 100.00 |
| Spektrum 4 | 23.08 | 41.62 | 8.91 | 1.15 | 20.13 | 5.12 | 100.00 |
| Max. | 24.26 | 42.37 | 14.03 | 3.51 | 37.24 | 19.42 | |
| Min. | 19.06 | 29.88 | 4.52 | 1.15 | 20.13 | 2.26 | |

**Fig. S84. Chemical nature of *EM-P*'s crust.** Image A shows a 30-month-old *EM-P* culture at the bottom of the bottle. Over the course of incubation, *EM-P* biofilm transformed into a brittle wafer due to salt encrustation. Images B & C show SEM images of *EM-P*. Scale bars in these images are 2µm. The elemental composition of the solidified mat is shown in D (determined by SEM-EDX).

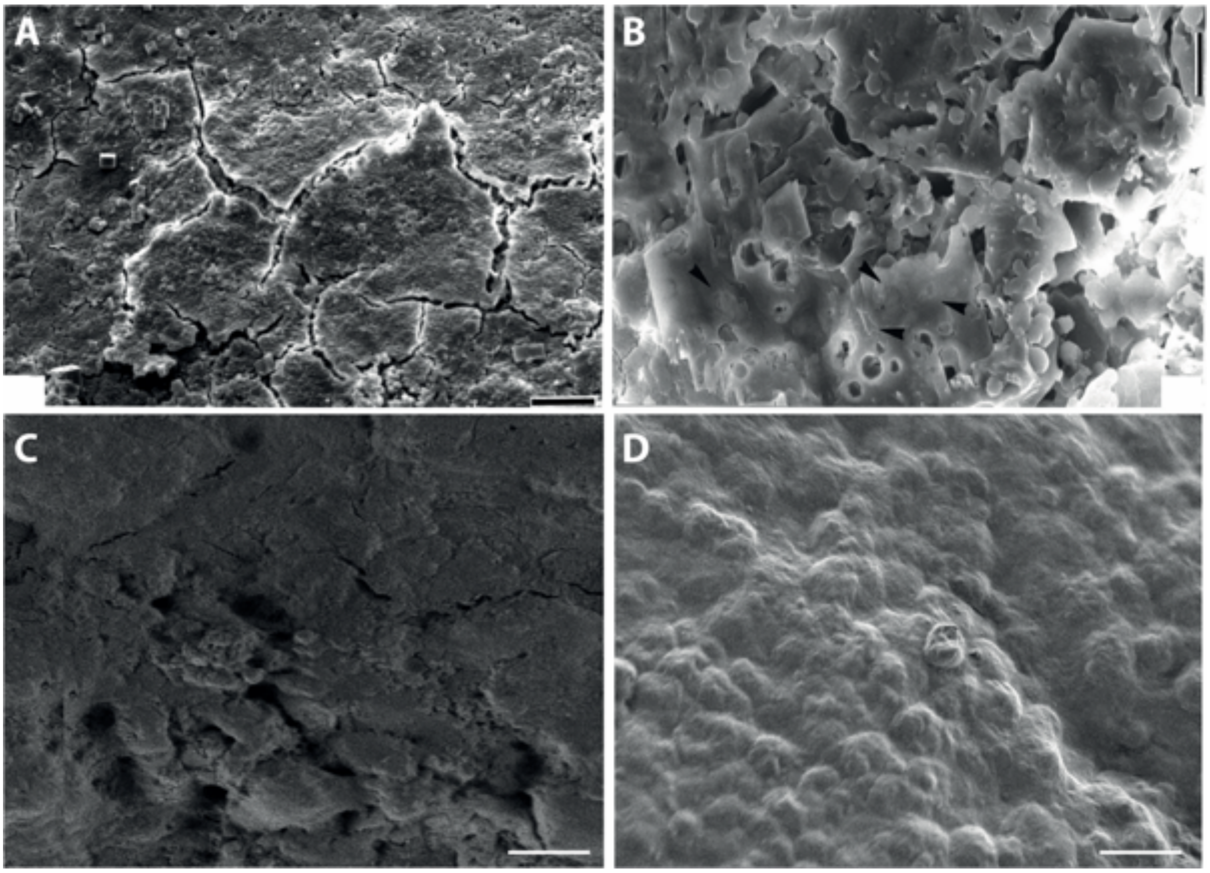


**Fig. S85. Morphological comparison of the Kromberg microfossils with *EM-P*.** Images A & B show mineral-encrusted microbial mats (A) and spherical microfossils (B) within these encrusted mats reported from the Kromberg Formation (originally published by Westall *et al.,* 2001)(62). Scale bars in images A & B are 20µm. C & D show SEM images of morphologically analogous salt-encrusted *EM-P* cells after 30 months of incubation. The elemental composition of such microbial mats and individual cell morphologies are shown in Fig. S84. Scale bars in images C & D are 2µm.

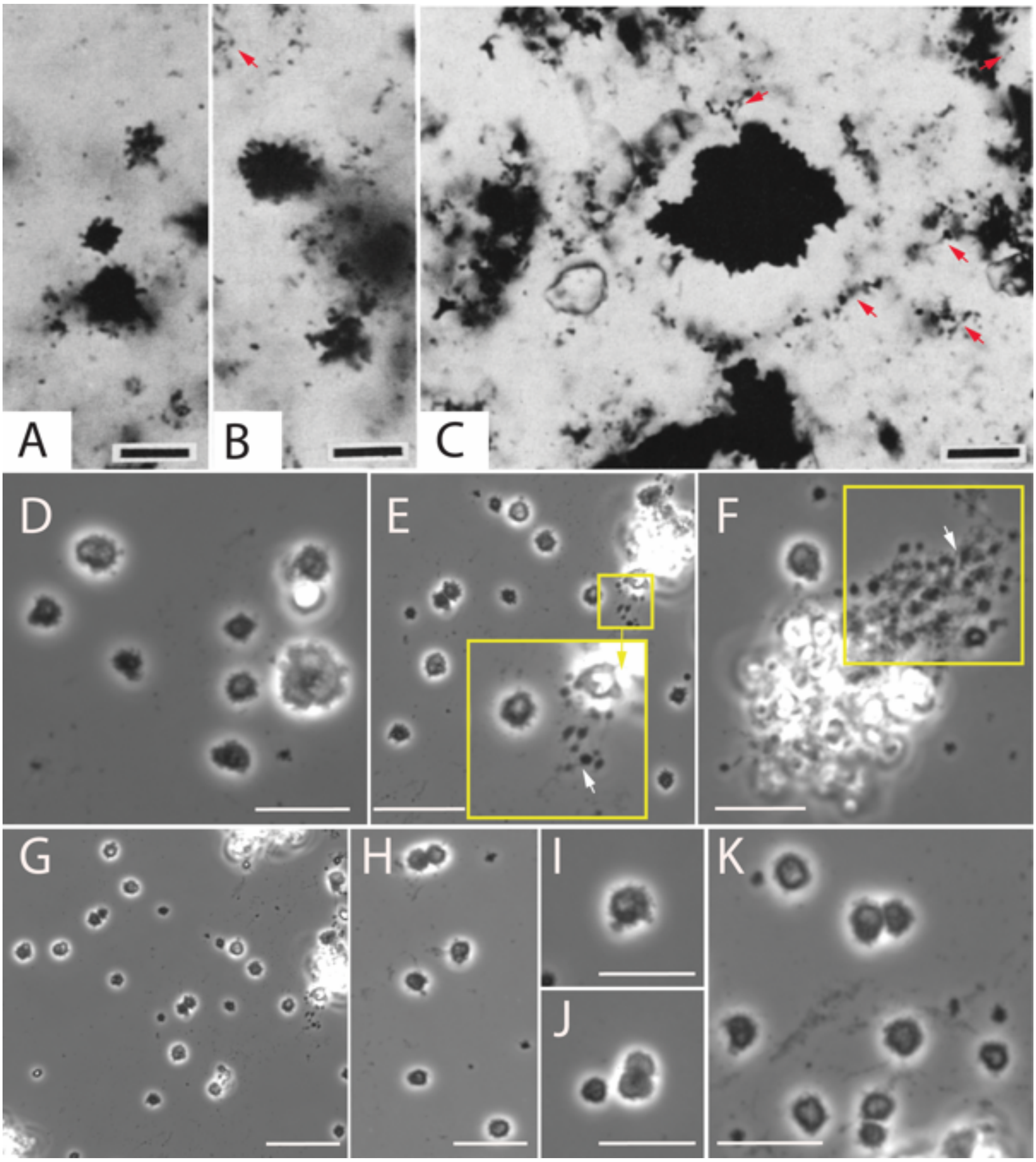


**Fig. S86. Morphological comparison of the North-Pole microfossils with *EM-P*.**

Images A-C show stellate shaped structures with filamentous projections reported from the North-Pole locality (originally published by Buick *et al.,* 1990)(45). Scale bars: 100µm. D-K are phase-contrast images of morphologically analogous *EM-P* cells. The stellar-shaped structure could have been the biologically induced mineral formed on the surface of the cells, as shown in Fig. S84. The filamentous structures were the daughter cells growing out of the crust when encrusted cells were transferred into fresh media. Arrows in images C, E & F point to a filamentous string of daughter cells extending out of cell clumps (Movie 22).

Images D-K also show salt-encrusted individuals or cells in dyads undergoing binary fission. Scale bars: 10µm.

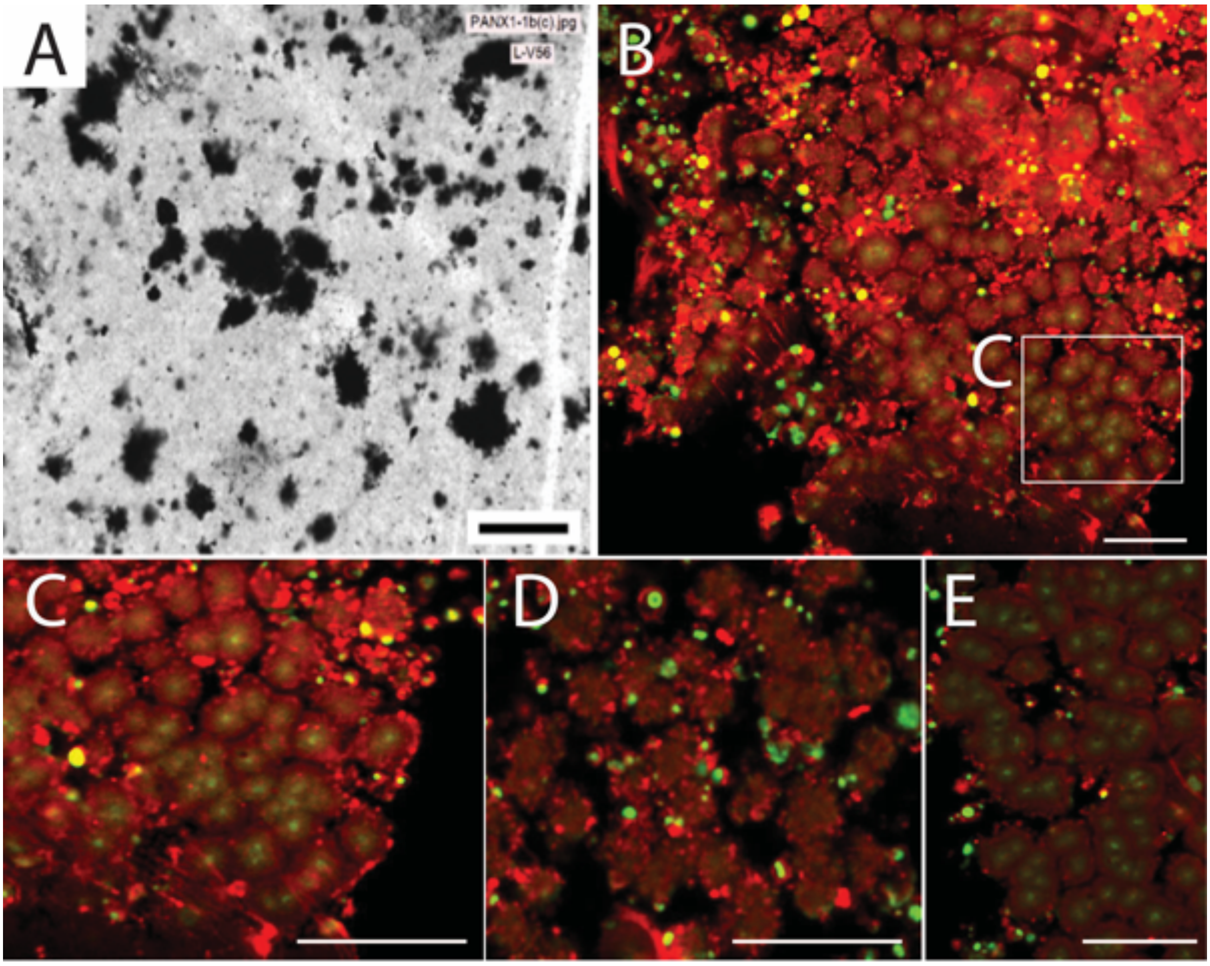


**Fig. S87. Morphological comparison of the SPF microfossils with *EM-P*.** Image A shows stellate microfossils with filamentous projections reported from the SPF (originally published by Sugitani *et al.,* 2013)(39). Scale bar: 200µm. Images B-E are spinning-disk confocal images of morphologically analogous *EM-P* cells. Cells in these images are stained with membrane stain, FM™5-95 (red) and PicoGreen™ (DNA, green). Like microfossils, *EM-P* cells revived from the above-described crust (Fig. S86) formed filamentous daughter cells from below the stellate structures. Scale bars: 10µm.

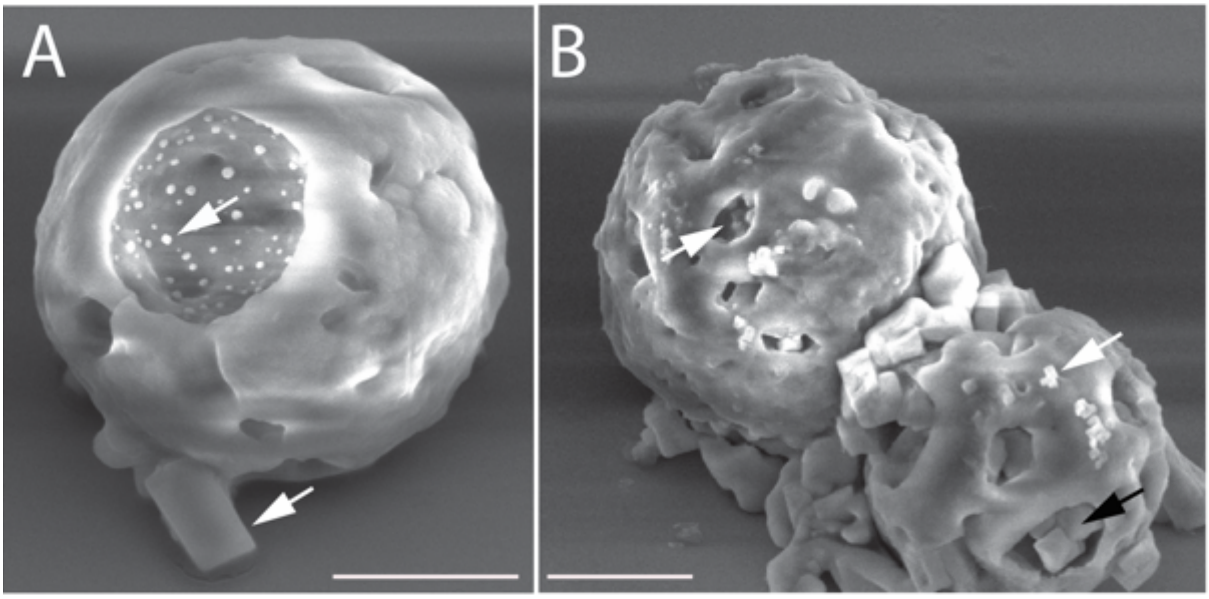


**Fig. S88. Association of salt with *EM-P* & *RS-P*.** Images A & B are SEM images of *EM-P*. Image-a shows *EM-P* cells forming membrane invaginations that resemble an endocytosis vesicle. These images *EM-P* with salt crystals within the membrane vesicles (closed arrows). Scale bars: 2μm.

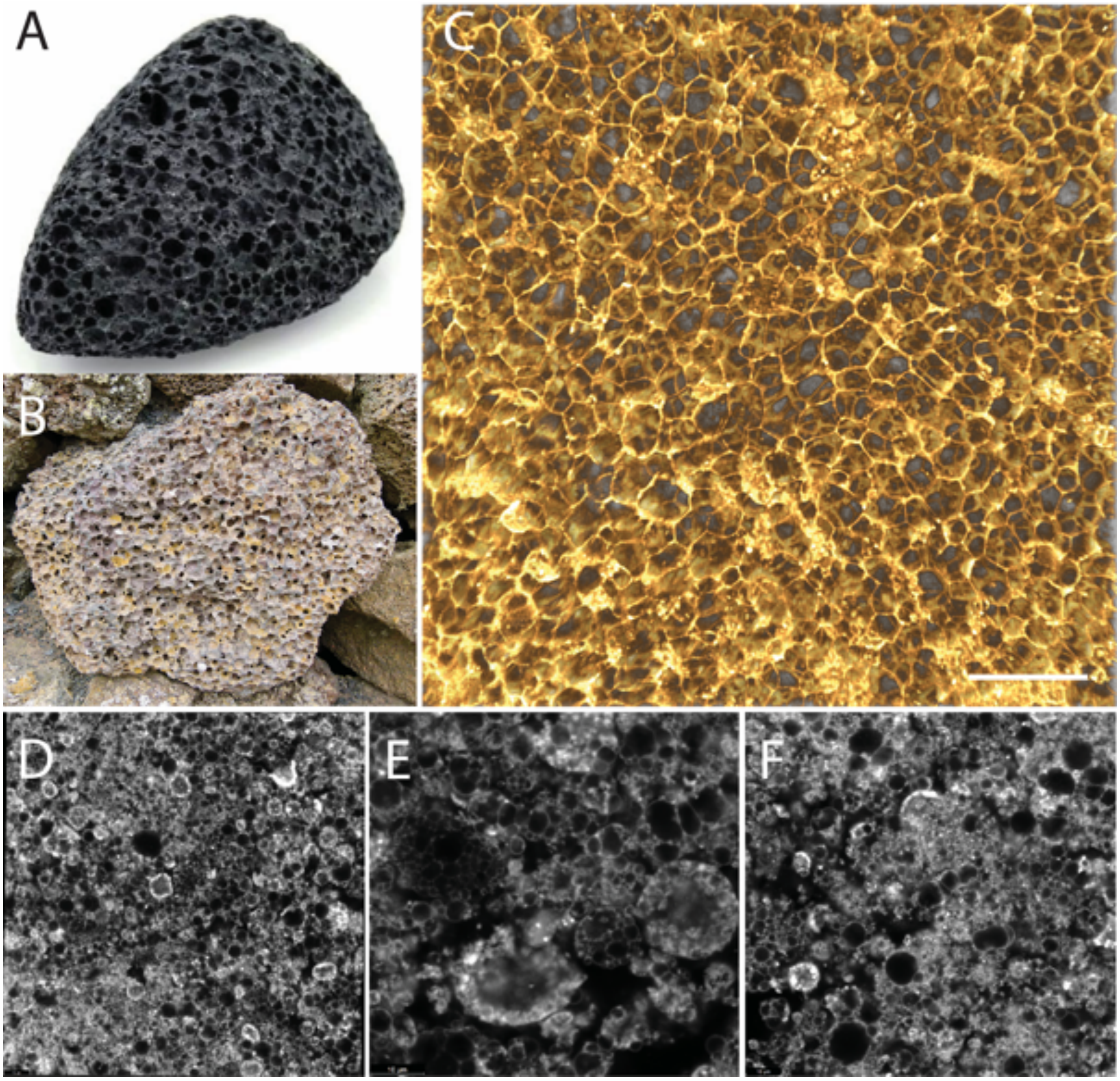


**Fig. S89. Morphological comparison of volcanic pumice with honeycomb-like structures observed in *EM-P* incubations.** Images A & B are the images of volcanic pumice (World Wide Web). Images C-F are the images of *EM-P* biofilm. Scale bars: 20µm (C) and 10 µm (D-F).

**Supplementary movies:**

**Movie 1.** Movies show an *EM-P* cell with an intracellular daughter cell within its vesicle. Scale bar: 5µm.

**Movie 2-4.** Movies show a gradual increase in the number of daughter cells within *EM-P*s intracellular vesicles and a corresponding decrease in the volume of the cell cytoplasm. Scale bar: 5µm.

**Movie 5-9.** Movies show sequential stages involved in *EM-P* cell lysis and the release of intracellular vacuoles containing daughter cells. Movies also show the formation of cell debris during the release of ICVs. Scale bar: 5μm.

**Movie 10-12.** The movie shows cell debris of *EM-P* cells with tiny daughter cells attached to them. Scale bar: 5μm.

**Movie 13-16.** Movies show sequential stages involved in the detachment and fragmentation of these strings of daughter cells into individual daughter cells (Movie 15). Movie 16 show the constant movement of the daughter cells, which should have provided the kinetic energy required for the fragmentation of strings of daughter cells into individual daughter cells. Scale bar: Movie 13 & 14 (5μm), Movie 15 (10μm).

**Movie 17.** The movie shows cell debris of *EM-P* cells with tiny daughter cells attached to them. Scale bar: 5μm.

**Movie 18.** The movie shows cell debris being formed within the biofilm. The lateral view of the biofilm shows the membrane debris being pushed out of the biofilm. Cells in this movie are stained with membrane stain FM$^{TM}$5-95 and imaged using a STED microscope.

**Movie 19.** The movie shows a layer of *EM-P* cells and membrane debris that was formed over this layer. Several spherical cells can be seen attached to the membrane debris and in the free space between the cell layer and the wavy membrane. Membrane debris was often noticed to have lenticular gaps. Cells in this movie are stained with membrane stain FM$^{TM}$5-95 and imaged using a STED microscope.

**Movie 20.** The movie shows layered membrane debris of *EM-P* forming hollow lenticular structures. Such structures should have formed by random folding of membrane debris as no indication of trapped cells was observed in these structures. Scale bar: 20μm.

**Movie 21.** The movie shows solidified *EM-P* biofilm. Hollow lenticular structures could be seen within the layers of membrane debris. Such structures should have formed honeycomb patterns within these lenticular gaps suggesting the presence of one's intact *EM-P* cells within these structures. Scale bar: 20µm.

**Movie 22:** The movie shows a string of *EM-P* cells growing out of the stellar-shaped salt-encrusted cells. Scale bar: 10µm.
